# Cryo-EM structure of CLCA1 identifies CLCA1 as a founding member of a novel metzincin family

**DOI:** 10.1101/2025.10.18.683246

**Authors:** Elisabeth Nyström, Sjoerd van der Post, Doireann Bradley Barrett, Grete Raba, Thaher Pelaseyed, Mihai Oltean, Ana S. Luis, Sergio Trillo-Muyo

## Abstract

Calcium-activated chloride channel regulator 1 (CLCA1) is implicated in several diseases, especially mucus-associated airway diseases, but its molecular function and regulation have remained unclear. By determining the structure of CLCA1 by negative stain electron microscopy and cryo-EM, we could confirm that CLCA1 forms large oligomeric complexes which adopts a compact domain organization comprising a metallohydrolase (MH), von Willebrand type A (VWA), β-sheet-rich (BSR), inhibitory (ID), and fibronectin type III-like (FnIII-l) domains. The unusually large MH domain bears hallmarks of metzincins but is distinguished by several unique features including an atypical active site zinc-coordination environment and a second Zn^2+^ -coordination site. Unlike classical metzincins, CLCA1 lacks a pro-domain; instead, a C-terminal inhibitory loop occludes the MH active site, providing an alternative mechanism of autoinhibition. The adjacent VWA domain, resolved in its closed state, is poised for conformational change upon ligand binding, suggesting a route for allosteric regulation of protease activity. Structural and functional assays support a role for CLCA1 in cleaving glycosylated substrates, leading to alterations in mucin architecture consistent with a regulated function in mucus remodeling. Together, these data establish CLCA1 as the founding member of a new eukaryotic metzincin family, here termed CLCAsins, with unique regulatory mechanisms.

## Introduction

The protein calcium activated chloride channel regulator 1 (CLCA1) has been implicated to play a role in several diseases, most noticeably mucus-associated airway diseases such as Th2-driven asthma, chronic obstructive pulmonary disease (COPD) and cystic fibrosis (CF), but also in the expulsion of parasite infection, cancer and pneumonia as reviewed in (Liu and Shi 2019; Patel, Brett, and Holtzman 2009). The physiological function of CLCA1 is still under debate; on the one hand CLCA1 has been suggested to regulate the calcium activated chloride channel TMEM16A by stabilizing TMEM16A on the cell surface (Sala-Rabanal et al. 2015; Sala-Rabanal et al. 2017). On the other hand, CLCA1 has been suggested to modify mucus properties by inducing structural rearrangement due to enzymatic cleavages of secreted mucins (Nystrom et al. 2018; Nystrom et al. 2019). Thus, further studies are needed to determine the overlap between the suggested functions and their physiological relevance.

CLCA1 belongs to a larger family of conserved CLCA proteins. It is found throughout the metazoan part of the animal kingdom, commonly with several paralogs within each species in vertebrates. CLCA-like proteins have also been observed in prokaryotes, which have been suggested to have gained the gene by horizontal gene transfer from eukaryotes (Lenart et al. 2013). Most of these CLCA proteins have a conserved domain structure; an N-terminal metallohydrolase domain (MH) followed by a Cysteine-rich domain (C) and von Willebrand type A domain (VWA), and a C-terminal fibronectin type III (FnIII)-like domain. Additionally, following the VWA domain is a predicted β-sheet rich (BSR) region, and all CLCA paralogs except CLCA1 have a membrane spanning helix in the most C-terminal region (Patel, Brett, and Holtzman 2009; Sala-Rabanal et al. 2017).

The 914 amino acid product of *CLCA1*, as well as other CLCA members, is autocatalytically cleaved in the endoplasmatic reticulum into a larger N-terminal and smaller C-terminal product (NTP and CTP, respectively), which remain associated via yet unknown interactions (Bothe et al. 2011; Nystrom et al. 2019; Pawlowski et al. 2006; Yurtsever et al. 2012). Intermolecular disulfide bonds drive further oligomerization, and CLCA1 is secreted into the intestinal mucus layer or pathological airway mucus as ∼1 MDa oligomers. (Fernandez-Blanco et al. 2018; Nystrom et al. 2019).

The autocatalytic cleavage of CLCA1 is catalyzed by the conserved zinc-dependent MH domain within the NTP of CLCA1. The active site **HE**WA**H**LRWGVF**D** motif (with predicted zinc-coordinating amino acids in bold) is typical for that of the zincin family of proteases, and CLCA1 has been compared to matrix metalloproteases (MMPs) and a disintegrin and metalloproteinases (ADAMs), due to this similarity (Stocker et al. 1995; Pawlowski et al. 2006; Yurtsever et al. 2012). The autocatalytic cleavage does not appear to be crucial for folding or secretion (Bothe et al. 2011; Nystrom et al. 2018; Yurtsever et al. 2012), but several studies have indicated that the physiological function of CLCA1 is exerted by the NTP, and that the CTP thus might play a regulatory function (Nystrom et al. 2019; Sala-Rabanal et al. 2015; Yurtsever et al. 2012). However, the function of the different domains of CLCA1 remain poorly understood, and only its VWA domain has been studied at a structural level (Berry and Brett 2020).

By determining the structure of CLCA1 by cryogenic electron microscopy (cryo-EM) and negative staining electron microscopy (NS-EM), we highlight several characteristic features of CLCA1 indicating that CLCA1 represent a novel family of metzincins, which we have termed CLCAsins with a potentially unique regulatory mechanism of its enzymatic activity. Furthermore, the VWA domain undergoes a conformational shift upon ligand-binding which might be propagated to the MH domain for activation. The study opens up new avenues for further investigation of the regulatory mechanisms determining the activity and function of CLCA1 in health and disease, which will inform future studies of CLCA1 as a potential therapeutic target.

## Results

### Interactions in the C-terminal end drives CLCA1 oligomerization

Transiently expressed CLCA1 secreted from CHO-S cells eluted as large molecular-weight oligomers from size exclusion chromatography (SEC), in line with previous observations (Nyström JBC 2019 (Figure S1A)). Oligomers of CLCA1 were also visible by NS-EM and cryo-EM, in which CLCA1 appeared as flower-like structures (Figure 1A and 1F). Similar flower-like assemblies were detected in mucus collected from wild-type, but not Clca1-deficient mice, confirming that these structures represent the endogenous oligomeric organization of CLCA1 (Figure 1B–C).

**Figure 1:**
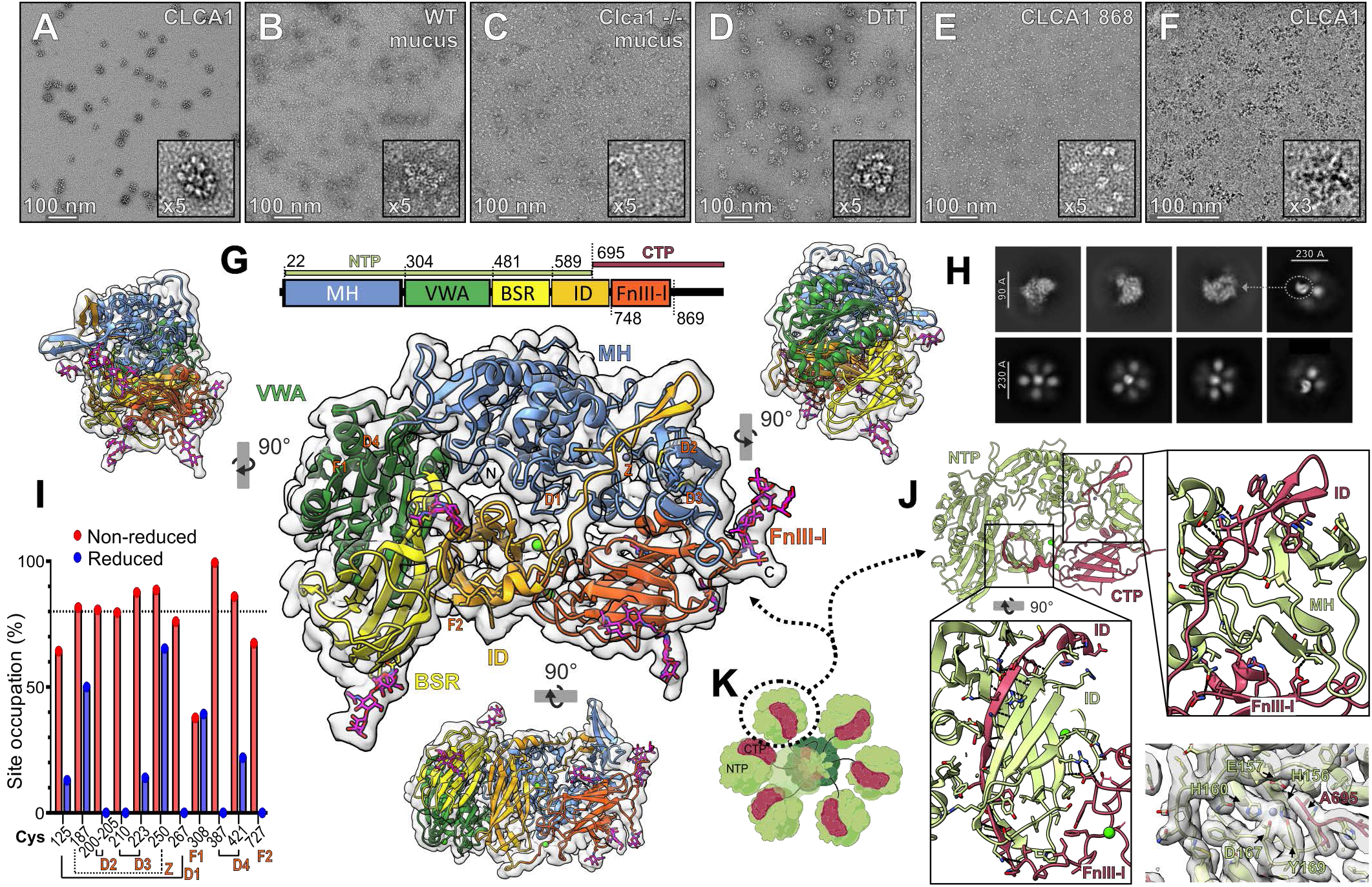
CLCA1 structure by NS-EM and cryo-EM. **1A-E**: Negative-staining electron microscopy (NS-EM) micrographs of recombinantly expressed and purified CLCA1 (A), WT or Clca1-/-colonic mucus (B and C respectively), DTT-treated recombinant CLCA1 (D), recombinantly expressed and purified CLCA1-886 (E). A 5× zoomed-in view is shown in the bottom right box. **F**: Cryo-EM micrograph of purified CLCA1 with a 3× zoomed-in box. **G**: CLCA1 Cryo-EM map and model. Schematic representation of CLCA1 at the top showing the N- and C-terminal cleavage products generated by autocatalytic processing (NTP and CTP, respectively); the position of the first residue in each domain or region is indicated. The CLCA1 model covering amino acid 22-868 is shown into the electron density map in cartoon representation, with cysteines, glycosylated asparagines, and glycans displayed as sticks. Disulfide-bonded (D), free (F), and zinc-coordinating (Z) cysteines are labelled, as are the N- and C-termini. A central frontal view of CLCA1 is shown reduced by 50% and rotated 90° clockwise (left) and anticlockwise (right) around the y-axis, and 90° clockwise around the x-axis (bottom). MH; metallohydrolase (blue), VWA; von Willebrand factor type A domain (green), BSR; β-sheet rich (yellow), ID; Inhibitory domain (orange), FnIII-l; Fibronectin type III-like (red). **H**: Representative cryo-EM 2D classes obtained from oligomer picking (bottom and top right) and monomer picking (top left). **I:** Cysteine site occupancy after differential alkylation in either reduced or non-reduced conditions. Solid lines; disulfide-bonded pairs (D1 to D4), dashed line; zinc-coordinating cysteines (Z); F; free cysteines (F1 and F2). Dashed line at 80% occupancy is arbitrarily to indicate fully occupied cysteines. **J:** Representation of interactions between the NTP (green) and CTP (red) and the density map of the autocatalytic site (insert). **K:** Schematic drawing of the proposed oligomerization of CLCA with the NTP and the FNIII-l domain from the CTP interacting in distinct petals, with the C-terminal tails extending to and interacting in a core. The number of petals may not be accurate.

The molecular structure of CLCA1 was solved at 2.99 Å resolution by cryo-EM (Figure S2, Table S1). This revealed that each “flower petal” consisted of a CLCA1 monomer, for which the structure for amino acids 22-868 could be solved, leaving the structure of the most C-terminal end undefined (Figure 1G). The interaction surface between individual petals could not be resolved, as inclusion of more than one monomer within the extraction box permitted accurate alignment of only one, while the others remained blurred (Figure 1H). We thus speculated that the most C-terminal ends hidden within the oligomeric structure mediated the intermolecular interactions (Figure 1K). This was partly confirmed by expression of a truncated CLCA1 lacking the most C-terminal end (CLCA-868), which was found as monomers both by SEC and NS-EM (Figure 1E and S1A).

### Mixed roles of cysteines in CLCA1

The CTP of CLCA1 contain two cysteines, Cys727 and Cys884, and CTPs of CLCA1 have been shown to interact by intermolecular disulfide bonds (Nystrom et al. 2019). This disulfide bond is most likely driven by homodimerization of Cys884 as Cys727 appeared as a free thiol in the structure (Figure S1E) and disulfide-mediated oligomerization of the CTP was disrupted in CLCA1-868 (Figure S1B). The cysteine site occupancy in Cys727 was confirmed by mass spectrometry by a differential alkylation strategy. The majority of cysteines involved in disulfide bonds in the structure were found to be 80– 100% occupied by mass spectrometry, with occupancy decreasing to 0–20% upon reduction with DTT. However, site occupancy for Cys727 was found to be approximately 70% (Figure 1I). Parallel analysis of nonreduced and alkylated CLCA1 indicated that occupied thiols on Cys727 to a large extent were modified by glutathione and cystine adducts, rather than involved in disulfide bonds. These reducible modifications are considered artifacts of secreted protein production introduced by the excess of glutathione and cysteine present in the culture media and are compatible with the observed electron density (Zhong et al. 2017)(Fig S1D). Similar to Cys727, Cys125 was found to be only partially occupied in non-reducing conditions. Although the disulfide bond is observed in the cryo-EM map, its electron density is relatively weak, further suggesting incomplete occupancy (Fig S1D).

Differential alkylation analysis also indicated a free cysteine in position 308. Cys308 has previously been shown to be unpaired (Berry and Brett 2020) and was primarily found as a free thiol or with cysteine adducts (Figure 1I and S1D). Furthermore, Cys187 and Cys250 remained un-reactive to alkylating agents upon reduction which was found to be due to metal coordination (further discussed below). Cys884 could not be analyzed by mass spectrometry because it lies in an *O*-glycosylated region (Figure S1C) and was not detected in our proteomics analysis.

Disulfate-mediated dimerization has been suggested to contribute to the oligomerization of CLCA1 (Nystrom et al. 2019). However, reduction with DTT did not cause disruption of full-length CLCA1 (CLCA1-FL) oligomers neither in NS-EM or SEC, indicating that interactions other than disulfide-bonds mediated CLCA1 oligomerization (Figure 1D and S1A). Although the structure was not solved experimentally, AlphaFold3 predictions of the C-terminal end suggest intermolecular hydrophobic interactions involving an α-helix in the C-terminal tail, corresponding to the membrane spanning helix in membrane bound orthologs of CLCA1, which could facilitate the oligomerization (Figure S1C).

### Global domain architecture and glycosylation of CLCA1

The cryo-EM structure largely confirmed the predicted domain boundaries of CLCA1; a MH followed by a VWA domain, a BSR region which was revealed to consist of two independent domains, a BSR domain and an Inhibitory domain (ID), and a FnIII-like (FnIII-l) domain in the CTP (Figure 1G) (Yurtsever et al. 2012). Each domain contacts the next, while the terminal FnIII-l domain connects to the initial MH domain, thereby conferring a compact conformation. Additionally, the ID domain interacts with all other domains in the molecule. Contrary to previous predictions, the ID transverse the autocatalytic site *via* a flexible loop. Thus, the NTP and CTP remained tightly associated despite autocatalytic cleavage, due to interactions between the ID and MH domains, but primarily through strong interactions within the ID, where a β-strand from the CTP contributes to an NTP β-sandwich (Figure 1J) (further discussed below).

CLCA1 has eight predicted *N*-glycan consensus sequences of which seven are within the amino acid range covered by the resolved structure (Gupta and Brunak 2002). The density map indicated the presence of seven *N*-linked glycans on Asn503, Asn585, Asn770, Asn804, Asn810, Asn831 and Asn836 of which the composition could be partly resolved ranging from a single GlcNAc to ManGlcNAc(2) with a core fucose (Figure 1G and S3C). Fucosylation at Asn503 and Asn836 was confidently modelled, while at Asn585 and Asn804 only weak density was observed due to the lack of stabilizing interactions, with no evidence at the other positions. However, the inherently flexible nature of both *N*- and *O*-linked glycan modifications limits the ability to resolve glycosylation by cryo-EM. Thus, complementary glycopeptide analysis by mass spectrometry was used for identification of both the composition and abundance of *N*-glycans at three positions, Asn503, Asn585 and Asn770 (Table S2). Asn503 and Asn585 were predominantly occupied with the complex glycans NeuAcHex(5)HexNAc(4) and FucHex(5)HexNAx(4) respectively while at position Asn770 a high mannose structure Hex(5) HexNAc(2) was identified with varying hexose additions (Figure S3B). The discrepancy regarding fucosylation at Asn503 could be explained by an oxidation event in the peptide or by another modification, although the exact cause could not be determined. Thus, all predicted *N*-glycan sites were confirmed to be occupied, and an additional model was generated including all of them to better understand their role on the protein structure and their impact on CLCA1 interactions and substrate recognition (Figure S3A).

Previous analysis has predicted several *O*-glycosylation sites in CLCA1 (Nystrom et al. 2019). However, only two sites, Ser110 and Thr111, could be confirmed to be modified by *O*-glycosylation, either a Core 1 or single Tn antigen (Table S2). The residues were only partly modified and identified both as modified and non-modified peptides by the mass spectrometry analysis and are therefore lacking in the cryo-EM structure. Potential *O*-linked glycosylation of the mucin-like regions rich in serine, threonine and proline in the C-terminal could not be determined by either method due to the unresolved area in cryo-EM and unsuitable peptide generation for mass spectrometry analysis.

### CLCA1 is a founding member of a novel family of metzincins

As predicted by the presence of a conserved HEXXE motif, the MH domain of CLCA1 revealed a structure typical of zinc metalloproteases, including a top-located twisted b-sheet (βI-VI), a backing alpha helix (αA), the active site helix (αB), and a downstream third alpha helix (αC-’’) (Figure 2A) (Cerda-Costa and Gomis-Ruth 2014; Gomis-Ruth, Botelho, and Bode 2012). The zinc ion is coordinated by His156 and His160 from helix αB, together with Asp167, whereas Tyr169 is positioned nearby and likely contributes to stabilization of the metal-binding site (Figure 2A, B and E). Furthermore, Met238 forms a so-called Met-turn just below the active site (Gomis-Ruth, Botelho, and Bode 2012), placing CLCA1 in the Metzincin clan of proteases together with e.g. MMPs and ADAMs (Cerda-Costa and Gomis-Ruth 2014). The unusually large MH C-terminal subdomain in CLCA1 does however separate the Met-turn from the active site helix in the primary structure, which is probably why this was not recognized before.

**Figure 2:**
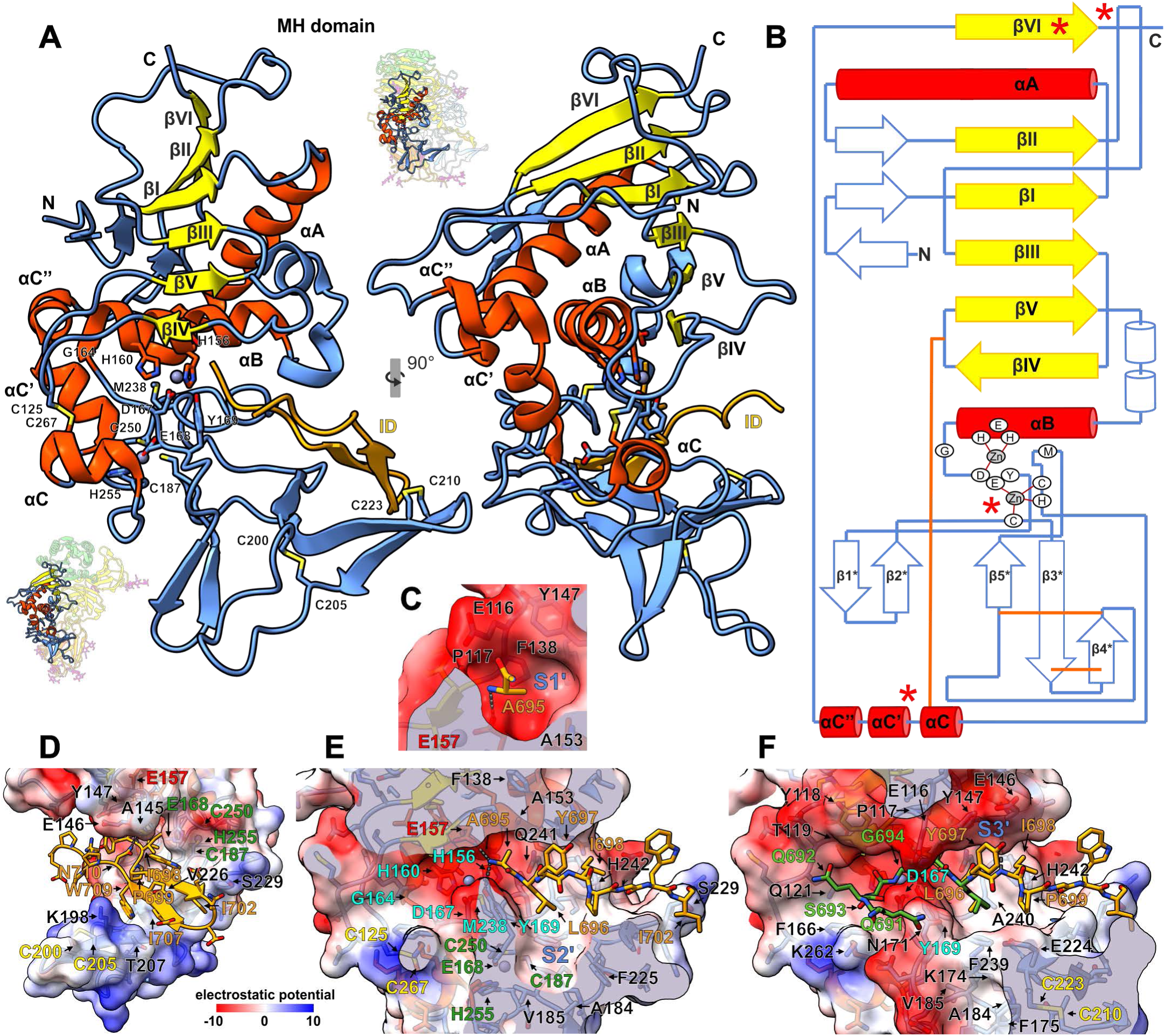
CLCA1 metallohydrolase (MH) domain. Conserved metzincin β-strands and α-helixes are shown in yellow and red, respectively, with the zinc atoms in grey and the remaining MH domain structure in blue. The inhibitory domain (ID) is shown in orange. **A:** Cryo-EM model of the CLCA1 metallohydrolase (MH) domain. The domain is displayed in the standard orientation (left) and rotated 90° anticlockwise around the y-axis (right), including the active-site insertion from the ID. Metal-coordinating residues, cysteines, and key residues of the metzincin motif are labelled, as are the N- and C-termini, and shown as sticks. The full CLCA1 molecule is shown next to each view to illustrate the relative position of the MH domain. VWA (green), BSR (yellow), ID (orange), FnIII-l (red), and N-glycans (magenta). **B:** Schematic representation of the MH domain. Disulfide bonds are shown in orange, and asterisks indicate characteristic features of the CLCAsin family. **C–F**: Details of the active site are shown in cartoon and stick representations. A semi-transparent surface representation colored by electrostatic potential is displayed for the MH domain. Interacting residues between the MH domain and the ID are labelled in black and orange, respectively; disulfide-bonded cysteines in yellow; key active-site residues in cyan; the general base (E157) in red; and residues coordinating the second zinc ion in dark green. Detail of A695 occupying the S1ʹ pocket (C), overview of the ID inhibitory loop interaction (D), close-up of the ID interaction within the active site, with the plane cut to better visualize the depth of the S2ʹ pocket and the second zinc-binding site (E), and AlphaFold model of P non-prime positions (green) in the inhibitory loop aligned with the cryo-EM structure (F). Putative interacting residues are labelled in green, also providing a clearer view of the residues defining the S2ʹ and S3ʹ pockets.

A structural comparison of the MH-domain using the non-redundant DALI PDB25 server indicated highest structural homology with eukaryotic M12 proteases (Figure S4A and Table S3), especially ADAM17 (z-score 7.3, rsmd 3.5) and ADAMTS-13 (z-score 7.1, rsmd 3.9). However, homology was also predicted with several prokaryotic metzincins known to cleave *O*-glycosylated mucin or mucin-like domains, including CpaA from *Actinobacteria* (z-score 7.3, rsmd 3.5, M72), OgpA (z-score 6.1, rsmd 3.9, M11) and Amuc_1438 from *Akkermansia muciniphila* (z-score 5.6, rsmd 3.9, unclassified) (Haurat et al. 2020; Trastoy et al. 2020; Medley et al. 2022). Similarity was also found with the archaemetzincin AmzA (Waltersperger et al. 2010) (z-score 5.8, rsmd 3.8, M54).

It was previously shown that CLCA1 has a DEY motif around the third metal-coordinating residue (Asp167) (Lenart et al. 2013). This motif is also found in M64 clan of bacterial proteases which are characterized by their ability to cleave near the *O*-glycosylated hinge region of IgA1 and IgA2m (Kobayashi et al. 1987). These proteases, also known as igalysins (Cerda-Costa and Gomis-Ruth 2014), constitute the metzincin family most closely related to CLCAsins, as they share distinctive variations of the zinc-binding motif, including an additional residue preceding the characteristic glycine (HEXXHXX<u>X</u>GXX<u>DEY</u>). In deposited structures of M64 proteases, such as the putative protease Bacova_00663 from *Bacteroides ovatus* (Joint Center for Structural Genomics 2010) and the recently published *Thomasclavelia ramosa* IgA peptidase (Ramirez-Larrota et al. 2025), the glutamate residue participates in the coordination of a second Zn²⁺ ion (Figure S4B). Archaemetzincins also coordinate a second zinc ion, but use a cysteine residue in place of the glutamate (Waltersperger et al. 2010). In both families, three additional cysteines complete the coordination: one located two positions upstream of the methionine in the Met-turn, another immediately preceding the αC helix, and the third within the first turn of this helix. In contrast, in CLCA1, the relative position of the second zinc ion is markedly different (Figure S4C), determined by the large region between αB and αC, as well as by the distinctive truncation of αC into three fragments, of which only the short fragment αCʹ occupies the position equivalent to αC in metzincins. Moreover, in CLCA1 one of the cysteines is replaced by a histidine. The first cysteine (Cys187) is located 51 residues upstream of the Met-turn, whereas Cys250 and His255 lie 21 and 16 residues upstream of αCʹ, respectively (Figure 2A, B, E and S4B). Notably, Cys250 and His255 are positioned directly at the base of αCʹ, suggesting that they may contribute stabilization effects similar to those of the last two coordinating cysteines in archaemetzincins and igalysins. The resolution achieved around the second Zn^2+^ in the cryo-EM map, together with the homology to the M64 clan, was sufficient to confidently build the current model. Furthermore, differential alkylation analysis confirmed the presence of a coordinated metal ion, as reflected by the decreased reactivity of Cys187 and Cys250 toward iodoacetamide (Figure 1I). Reduction in the presence of EDTA did not result in a complete displacement of the metal group indicating a high coordination strength. In line with this, other metzincins, e.g. ADAM17, contain a stabilizing disulfide bond at the base of αC indicating a structural role.

In reported structures, bacterial igalysins lack a prodomain, likely because their activity against *O*-glycosylated regions makes the production of an already active enzyme non-deleterious to the bacterium. However, metzincins are normally expressed as inactive zymogens where a cysteine or aspartate in an N-terminal pro-domain forms an extra coordination with the active site Zn^2+^, preventing entry of the solvent molecule needed for the nucleophilic attack (Cerda-Costa and Gomis-Ruth 2014; Guevara et al. 2010). Activation of the zymogen requires proteolytic cleavage and removal of the pro-domain, inducing a so called “cysteine or aspartate switch”. However, similar to igalysins, CLCA1 also lacks a classical prodomain; the N-terminal segment preceding the MH domain is too short to be able to interact with the active site cleft, in line with previous predictions (Yurtsever et al. 2012). The cryo-EM map shows density for the first amino acid after signal peptide cleavage, confirming its position adjacent to the VWA-BSR domain interface and distant from the catalytic zinc (Fig. 1G). Instead, the β6-β7 extension in the Inhibitory domain, which contains the autocatalytic cleavage site, occludes the active site and is kept in place by several interactions (Fig. 2A, D), highlighting the importance of maintaining the enzyme inactive until proteolytic activation is required (further presented below).

Another distinguishing feature of CLCA1 compared to previously analyzed metzincins includes a sixth β-strand (βVI) in the twisted β-sheet, arising from the C-terminal end of the MH domain, looping back behind the domain, and aligning in parallel with the β-sheet (Figure 2A-B, S4A). A similar arrangement is found in the bacterial *O*-glycoprotease StcE (Yu, Worrall, and Strynadka 2012), but the sixth strand adopts an antiparallel orientation. Igalysins also contain an additional antiparallel β-strand, whereas in this family it originates from the N-terminal region (Figure S4). Furthermore, as mentioned previously, the region between αB and αC of the MH in CLCA1 is unusually large. It spans 118 residues, including five β-strands (β1*-β5*), creating a deep active site cleft with a protruding thumb-like structure. It is stabilized by three disulfide bonds, Cys200-Cys205 between β3* and β4*, Cys210-Cys223 between the β4* and β5*, and Cys267-Cys125 which connects the αC helix with the βIV-βV loop (Figure 2A-B). The previously predicted C-domain (Yurtsever et al. 2012) is thus a part of the MH C-terminal subdomain and not a separate domain. This is in contrast to ADAMs and ADAMTSs which harbors a C-domain separated from the MH-domain by an elongated disintegrin domain (Seegar and Blacklow 2019; Gomis-Ruth, Trillo-Muyo, and Stocker 2012). Sequence alignment indicate that these unique features of CLCA1-MH are conserved throughout the CLCA-family with a few exceptions (Figure S5). For example, several CLCA1 orthologues have an asparagine rather than an aspartate in the third active-site Zn-coordination position. However, several of these have been shown to undergo Zn-dependent autocatalytic cleavage, indicating that they are expressed as active enzymes (Anton et al. 2005; Plog et al. 2009; Bothe et al. 2011).

Furthermore, the MH is a knotted domain (Kolesov et al. 2007). Protein knots are rare topological features that can influence folding pathways and, in some cases, confer enhanced stability and resistance to mechanical or thermal unfolding (Taylor 2000; a Beccara et al. 2013; Sulkowska et al. 2008a). Under mechanical stress, knots tighten and exhibit characteristic resistance to unfolding as well as a distinctive response to stretching (Sulkowska et al. 2008b).

MH forms a 3_1_ knot (trefoil knot) between Ile39 at βI and Gln301 at βVI. βVI directs the C-terminus through the βII–βIII loop, forming this unusual topology (Figure 2A–B). The side chain of Gln301 keeps the loop in place, forming hydrogen bonds with the side chain of Thr87 and the main chains of Pro82, Glu83, and Trp85 (Figure 3C). The loop is further stabilized by intra–main chain interactions between Ile81 and Ile302, as well as between Val92 and both Gln301 and Leu299 within βVI. Moreover, it interacts with Glu45 in the βI–αA loop, whose carboxyl group forms a hydrogen bond with the amide hydrogen of Lys86, while its aliphatic chain interacts with Trp85. Numerous additional hydrophobic interactions stabilize the structure, including contacts involving residues Tyr91, Val92, Pro94, and Tyr99 in the βII–βIII loop with Leu80 and Ile81 in βII, Leu299 and Leu300 in βVI, and Ile302 in the C-terminal loop connecting to the VWA domain.

**Figure 3:**
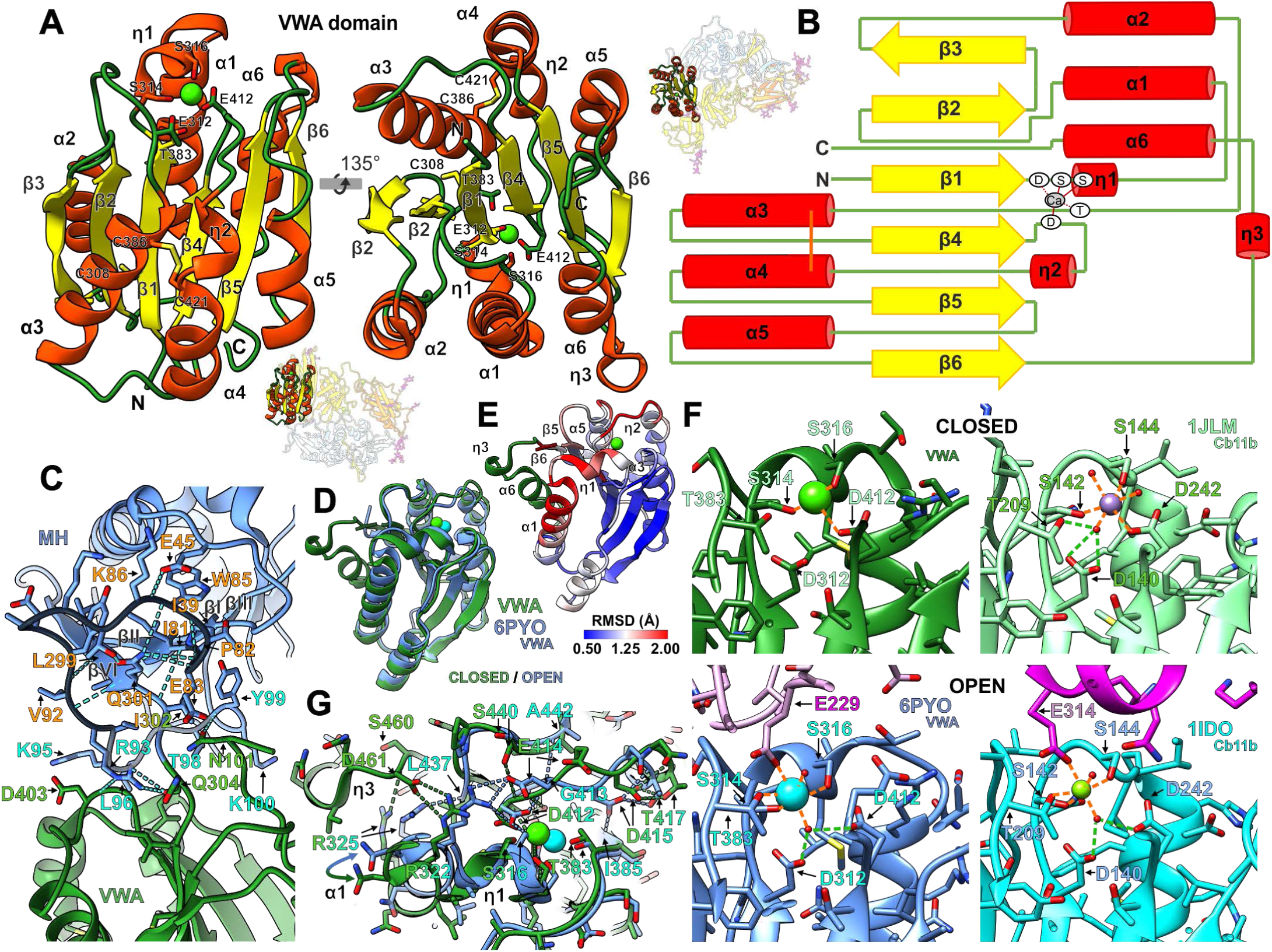
CLCA1 von Willebrand type A (VWA) domain in open and closed conformation. **A:** Cryo-EM model of the CLCA1 metallohydrolase VWA domain. β-strands are shown in yellow; α-helixes and 3₁₀ helixes (η) in red; the calcium ion in light green; and loops in green. The domain is displayed in two views, rotated 135° clockwise around the x-axis. Calcium-coordinating residues and cysteines are labeled, as are the N- and C-termini and the secondary structure elements, and shown as sticks. The full CLCA1 molecule is shown next to each view to illustrate the relative position of the VWA domain. MH (blue), BSR (yellow), ID (orange), FnIII-l (red), and N-glycans (magenta). **B:** Schematic representation of the VWA domain. Disulfide bonds are shown in orange. The same color code as in (A) is used. **C:** Interaction region between the MH (blue) and VWA (green) domains. Interacting residues in the MH domain are labelled in cyan and those in the VWA domain in green, and shown as sticks. The knot loop is shaded in black, and key residues stabilizing it are also shown as sticks and labeled in orange. Hydrogen bonds are represented by dashed cyan lines. **D-E:** Alignment of the VWA domain from the current structure in the closed conformation (VWA; green) with the previously reported open conformation (PDB ID: 6PYO; blue). Superposition is shown in (D), and the closed VWA domain colored by alignment root mean square deviation (rmsd) in (E). The region not resolved in the open-conformation structure and therefore not aligned is shown in green. Key secondary structure elements are indicated. **F**: Details of the MIDAS motif and ion-binding region. Closed (top; green) and open (bottom; blue for chain A and pink for chain B) conformations of VWA domains from CLCA1 (left) and CD11b (right). PDB IDs are indicated. Stick representation is shown. Water molecules are represented as red spheres. Calcium ions in CLCA1 structures are shown in green or cyan, whereas in CD11b structures, manganese (closed conformation) is shown in purple and magnesium (open conformation) in green. Metal coordination is depicted as orange dashed lines, and coordination of metal-bound waters as green dashed lines. Residues involved in metal coordination are labelled. **G:** Detail of (D) shown in stick representation. Dashed lines indicate the distinct hydrogen bonds characteristic of each conformation. Residues involved in the transition between the open and closed conformations are indicated. The movement of the α1 helix is highlighted by a double-headed arrow.

### The autocatalytic cleavage region occludes the active site

Substrate specificity for proteases is commonly determined by the recognition of three to five amino acids up- and downstream of the scissile bond (P3-P3’) in the substrate to pockets in the active site cleft (S3-S3’), with S1’generally exerting most substrate specificity in metalloproteases (Gomis-Ruth, Botelho, and Bode 2012). For metzincins, substrate binding is generally determined by the βIII-βIV loop, the βIV “upper wall”, βIV-βV loop, and the “floor” determined by the Gly-turn immediately following the active helix, the catalytic zinc with its coordinating imidazoles and the Met-turn (Stocker and Bode 1995). In CLCΑ1, the βV-αB loop is bulgier than in most other metzincins and Phe138 and Tyr147 are further delimiting the substrate binding cleft, and does together with Glu116 and Pro117 in the βIII-βIV loop, Gln241 in position 6 after the Met-turn and Ala153 in α2 make up a deep hydrophobic S1’pocket (Figure 2C, E and F). S2’ is also hydrophobic but wider, the upper part is delimited by Phe239 and Ala240 in position 4 and 5 after the Met-turn and Tyr169 (Figure 2F). S2’ extends for 15 Å till the aromatic ring of Phe175. The bottom region of the cavity is formed by Lys174, Ala184, Val185 and Phe225 (Figure 2E-F). The S3’ upper wall is formed by Glu146 and Tyr147 in the βV-αB loop.

However, potential substrate binding is conditioned by the presence of the Inhibitory domain β6-β7 extension. After autocatalytic cleavage, the neo-C-terminus of the NTP loses interaction with the active site and moves away from it; no electron density could be observed in the area where the region from Gly680 to Gly694 is expected to be located. In contrast, the cryo-EM map shows that the neo-N-terminus of the CTP (Ala 695 in P1’to Pro714) remain firmly attached to the MH domain, occupying the catalytic site (Figure S1E). The active site is occluded as a result of the neo-N-terminus at Ala695 forming a hydrogen bond with both the catalytic base (Glu157) and the tyrosine from the DEY motif (Tyr169). Ala695 and Leu696 lateral chains occupy the S1’ and S2’ hydrophobic pockets, respectively. Tyr697 specifically interacts with the S3’ upper wall, and a hydrogen bond is formed between the main chains of Ile698 in P4’ and Glu146. Furthermore, the hydrophobic face of a β-hairpin (Pro699, Ile702, Ile 707 and Trp709) interacts with the β3*-β5* sheet in the MH (Lys198, Thr207, Val226 and Ser229. Figure 2D). In the C-terminal region, an additional hydrogen bond between the side chain of Asn710 and the carbonyl oxygen of Ala145 contributes to the tight binding. In total, the interface area spans for 858 Å^2^ and shows a solvation free energy gain of −11.4 kcal/mol.

The lack of density from Ser693 and Gly694 in P1 and P2 position in the autocatalytic cleavage site prevents analysis of interactions on the non-primed side of the scissile bond. However, an AlphaFold2 prediction was used to infer their possible conformation (Figure 2F). The S1 position is open and only limited by the conserved Thr118. The hydrophobic wall created by the aromatic ring could accommodate hydrophobic resides or the aliphatic chain of Lys and Arg. In the autocatalytic cleavage site of CLCΑ1, a glycine (Gly694) is found in P1 position. However, in most other CLCAs, a basic sidechain is found in P1 position (Figure S5). Thr119 in βIV, Gln121 in the βIV-βV loop, Phe166 preceding the third zinc binding residue and Lys262 in αC appear to delimit S2 which is invariantly small polar in CLCAs autocatalytic cleavage sites. In CLCA1, the hydroxyl group of Ser693 could establish hydrogen bonds with the side chain of Gln121 and the main chain of Thr119. The glutamines at positions P3 and P4 could also contribute to the binding of the Inhibitory domain through interactions involving their aliphatic chains and Tyr118 and Asn171. However, it is not until position P2 that the chain turns almost 90° and makes tight contact with the active site.

The structural similarity between CLCA1 MH and glycoprotein-degrading metzincins and ADAMs suggests that CLCA1 is capable of cleaving near or at glycosylated sequences. In line with this, the S2’ pocket is large enough to accommodate a glycosylated residue and specifically bind glycans moieties through CH/π interactions. Noticeably, Phe175, Phe225 and Phe239 are mostly conserved between CLCA homologs (Figure S5). In addition, all positions around the cleavage site, with the exception of P1’, could accommodate a glycosylated residue without major steric impediments.

### Conformational alterations in the VWA domain upon ligand binding

The VWA domain is linked to the MH domain via the C-terminal part of the trefoil knot (Figure 3C). The interaction between the two domains is weak and limited to three residues in the MH βII-βIII loop. The interface is stabilized by a hydrophobic contact between Leu96 and Ile413 at VWA β5, hydrogen bonds between the guanidino group of Arg93 and the main-chain of Gln304 in the domain-linking loop and Asp403 in VWA α3-β4 loop; and a hydrogen bond between the main chain of Tyr99 and the side chain of Asn479 at the VWA C-terminus. Thus, the relative position of both domains appears to depend on additional contacts with the other protein domains (Figure 1G).

The core structure of VWA domains commonly contains a six-stranded β-sheet core surrounded by six α-helixes and three 3_10_ helixes (η). In contrast to an earlier study investigating the crystal structure of CLCA1 VWA (PDB code 6PYO), we were able to identify all these elements in the CLCA1 VWA domain, including the sixth α-helix (α6) (Berry and Brett 2020) (Figure 3A-B). The domain also contained a divalent cation in the predicted metal ion dependent adhesion site (MIDAS), coordinated by Ser314 and Ser316 in the DXSXS MIDAS motif in the β1-η1 loop and η1, and Asp412 in the β4-η2 loop (Figure 3A-B). The identity of the ion could not be determined from the structure and was annotated as Ca²⁺ due to the use of CaCl₂-containing buffers, although Mg²⁺ was shown to be a more likely ligand (Berry and Brett 2020).

The coordination of the metal differed compared to the previously published VWA structure in which the calcium ion was directly coordinated by Ser314 and Ser316 in the MIDAS motif, Thr383 situated in the α2-α3 loop, a water molecule immobilized by Asp312 in the MIDAS motif, Asp412, and Glu229(−3) in the expression tag from a second CLCA1 VWA molecule (pseudo-ligand). In the full CLCA1, the coordination by Ser314 and Ser316 remains unchanged whilst Thr382 and Asp412 exchange roles. Asp412 directly coordinates the ion, while Thr382 immobilizes a water molecule that, together with Asp312, coordinates the Ca²⁺. This water molecule is not visible in the cryo-EM structure, but its presence can be inferred from other VWA domain structures (e. g. the α-subunit of integrin CR3 (CD11b/CD18) in PDB entry 1JLM) where the involved residues adopt virtually identical positions (Figure 3F). The metal coordination differences are in line with an “open” and “closed” conformation of the VWA domain which refers to conformational shifts of the MIDAS and its surrounding loops upon ligand binding (Springer 2006; Emsley et al. 2000) (Figure 3D-G). Thus, in contrast to the previously reported isolated VWA domain structure of CLCA1 in the open conformation, full-length CLCA1 keeps the VWA domain in its closed conformation.

Structural alignment of the open and closed conformations reveals notable deviations around the MIDAS motif and Glu414, particularly in the upper part of α1, η1, the β4-η2 loop, the β5-α5 loop, and the upper region of β6 (Figure 3D-E), consistent with previous findings (Shimaoka et al. 2003). In the closed conformation, α6 interacts with the VWA hydrophobic core via Leu468, Phe472, and Leu475, orienting Asp461 toward the metal-binding site (Figure 3F). This position is stabilized by a hydrogen bond between Arg325 (α1) and the backbone carbonyl of Ser460, allowing Asp461 to form a salt bridge with Arg322, distancing it from the β4-η2 loop. When α6 is absent, α1 shifts closer, and Arg322 instead forms a salt bridge with Asp412 and Glu414. This interaction drives the rotation of Glu414, akin to conformational changes seen in integrin VWA domains upon ligand binding. In the open conformation, α1 is further stabilized by Arg325 forming a hydrogen bond with the backbone carbonyl of Leu437 (β5-α5 loop), propagating the conformational change into this region.

In the β4-η2 loop, the hydrogen bond between the side-chain of Ser440 and the main chain of Asp412 observed in the closed conformation is replaced by a series of intra-main chain hydrogen bonds involving Ala442, Gly413, and Asp415 in the open conformation, facilitating Glu414 rotation. Indeed, this loop, which contains Glu414, shows the most pronounced conformational changes.

Asp412, at the loop’s N-terminus, maintains a similar position in both conformations; however, α1 displacement in the closed state brings the MIDAS motif closer to Asp412, enabling direct metal coordination. Moreover, in the open conformation, Asp412 is restrained by the previously mentioned salt bridge with Arg322, as well as by hydrogen bonds with the amide hydrogen of Glu414 and the side chain of Ser316.

Asp415 also plays a key role in the conformational change. Asp415 side chain forms hydrogen bonds with Thr417 main and side chain at the η2 N-terminus, but in the closed conformation, it also interacts with Ile385 bringing η2 and α3 closer together. This interaction does not account for the different metal coordination mechanism observed in the nearby Thr382, which, like Asp412, does not undergo significant changes in its relative position but is affected by the displacement of α1 and η1.

In Integrins, transition between the open and closed state induce a large conformational change by shifting the α6-terminal helix up to 10 Å “downwards”, allowing the altered position of α1 (Emsley et al. 2000; Shimaoka et al. 2003; Springer 2006). The α6 electron density is missing from the previously described structure of CLCA1 VWA, suggesting that it extends from the core, allowing an open conformation. However, the relative position of the VWA, especially α6, in relation to the rest of the protein in full CLCA1 might restrain the conformational alterations needed for the transition from the closed to open state similar to what has been suggested in regards of the VWA (inserted-) domain in Integrin a2b1 (Emsley et al. 2000) (Figure 1E; further discussed below).

### BSR and the Inhibitory domains

The region following the VWA has previously been referred to as an unassigned β-sheet-rich (BSR) domain (Yurtsever et al. 2012). However, the region is composed of two domains, which we tentatively refer to as BSR and Inhibitory domain (Figure 1G and I). The BSR domain adopts a β-jellyroll fold, consisting of two parallel β-sheets. The first comprises five β-strands (β1, β8, β3, β6 and β5), while the second consists of three β-strands (β2, β7 and β4) (Figure 4A-B). In ADAMTS13, the spacer domain shares the same connectivity as the BSR domain but forms an extended β-sandwich through the addition of two β-strands in the β5-β4 region (Figure S6A-B). The interface between the two β-sheets in the BSR domain is strongly hydrophobic, as is typical for this fold. The external face of the first β-sheet is also fairly hydrophobic; Leu517 and Leu519 in β3, Met550 and Tyr552 in β6, and Val543 in β5 form a hydrophobic patch, whereas β1 and β8 are primarily composed of short polar amino acids. In contrast, the second β-sheet is hydrophilic and shielded by a complex *N*-glycan placed at Asn503 (β2). The proximal GlcNAc interacts with Trp501 in β2 and Ser567 in β7, while the fucose interacts with Lys565 in β7 and Trp532 in β4 (Figure 4G and S3C). Thus, only the external face of the first β-sheet and the lateral faces of the hydrophobic core are available for protein-protein interaction.

**Figure 4:**
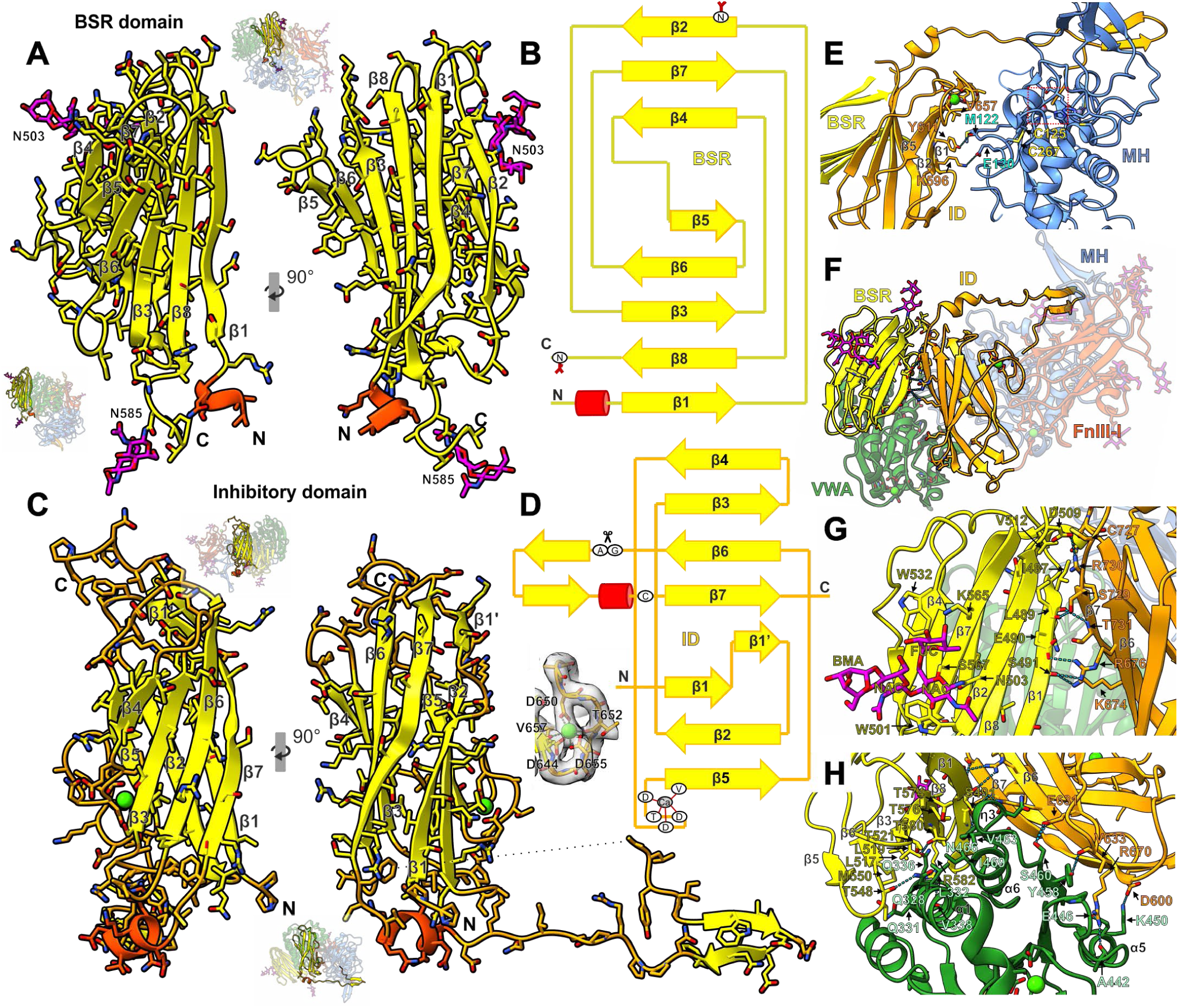
β-sheet rich and Inhibitory domain of CLCA1. **A:** Cryo-EM model of the BSR domain in cartoon and stick representation. β-strands and loops are shown in yellow, α-helices in red, and *N*-glycans in magenta. The domain is displayed in two views, rotated 90° clockwise around the y-axis. *N*-glycosylated asparagine residues are labelled, as are the N- and C-termini and the secondary structure elements. The full CLCA1 molecule is shown next to each view to illustrate the relative position of the BSR domain. MH (blue), VWA (green), BSR (yellow), ID (orange), FnIII-l (red), and N-glycans (magenta). **B:** Schematic representation of the BSR domain. *N*-glycosylated asparagine residues are marked. The same color code as in (A) is used. **C:** Cryo-EM model of the ID domain in cartoon and stick representation. β-strands are shown in yellow, α-helices in red, calcium in green, and loops in orange. The domain is displayed in the same orientation as the BSR domain (A) in two views, rotated 90° clockwise around the y-axis. The N- and C-termini and the secondary structure elements are labelled. The full CLCA1 molecule is shown next to each view to illustrate the relative position of the ID domain. The same domain color code as in (A) is used. A close-up of the calcium-binding site is shown together with the cryo-EM map, with residues involved in coordination indicated. **D:** Schematic representation of the ID domain. The CLCA1 autocatalytic cleavage site is indicated. The same color code as in (C) is used. **E:** Detail of the interaction region between the MH (blue) and ID (orange) domains. Active-site residues are shown as sticks and highlighted by a red dashed box. Interacting residues not involving the inhibitory loop are labelled and also shown as sticks, as are the cysteines stabilizing the MH interacting loop. **F-H:** Detail of the VWA–BSR–ID domain interactions. Hydrogen bonds are shown as dashed lines. The same domain color code as in (A) is used. Key interacting residues are labeled and shown as sticks. Overall view of the interaction (F); detail of the BSR–ID interface and interactions between the *N*-glycan at asparagine 503 and the BSR domain (G); and VWA interactions with the BSR and ID domains.

A short linker, shielded by a second *N*-glycan (Asn585), connects the BSR to the Inhibitory domain. The ID adopts a β-sandwich conformation with a prominent extension between β6 and β7, and an ion binding site at β4-β5 loop coordinated by Thr652 and Val657 backbone carbonyl oxygens and the carboxyl groups of Asp644, Asp650 and Asp655 (Figure 4C-D). The first β-sheet consist of three β-strands (β1, β2, and β5) and the second includes five (β1’, β7, β6, β3 and β4). The hydrophobic core between the two β-sheets is concealed by the close contact between β7 and β1 on one side, and the β4-β5 loop on the other. The external faces of both β-sheets are prominently hydrophilic.

The external face of the first β-sheet interacts with the βIV-βV loop in the MH domain, whose opposite face contributes to the substrate-binding pocket. The interaction is limited to a hydrogen bond between the side chain of Tyr661 in β2 and the main chain of Met122, further stabilized by hydrophobic interactions among their side chains and Val657 in β5, as well as a salt bridge between Lys596 and Glu130 (Figure 4E). Though, the main interaction between the MH and the Inhibitory domain is mediated by the extended region between β6 and β7. It spans from Gly680 to Cys727 and inserts directly into the MH active site, where it is cleaved yet remains bound, thereby inhibiting enzymatic activity (see “Substrate binding”). The residues Glu715-Lys724 are visible in the cryo-EM map so they were modelled despite the low resolution achieved in the area (Figure S1E). The model shows that the β6-β7 extension of the Inhibitory domain is connected to β7, in contrast to its N-terminal region. However, the structural details of this area should be interpreted with caution. The low resolution suggests that it is a flexible linker that does not engage in significant interactions with other CLCA1 domains.

The BSR and Inhibitory domains interact through the proximal region of their edge β-sheets (β1 and β2 in the BSR, and β1 and β7 in the Inhibitory domain) adopting a V-shaped conformation (Figure 4F-H). Limited hydrophobic interactions are stabilized by hydrogen bonds and salt bridges, primarily involving β1 and, to a lesser extent, the β2-β3 loop of the BSR domain, and β6 and β7 of the Inhibitory domain (whose connecting segment is cleaved). In β7 of the Inhibitory domain, Arg730 forms a salt bridge with Asp509 in the β2-β3 loop; Thr731 establishes a main-chain hydrogen bond with Gln488, and the side chain of Ser729 forms a hydrogen bond with Leu489. Both Gln488 and Leu489 are located in the β1 strand of the BSR domain. Lys674 and Arg676, located in β6 of the Inhibitory domain, extend their side chains towards the BSR domain and interact with the β1 strand. Both form hydrogen bonds with the main chain of Ser491, while Arg676 additionally establishes a salt bridge with Glu490 (Figure 4G).

The V-shape formed by the interaction between the BSR and the Inhibitory domain further interacts with VWA, positioning η3 and α6 at its vertex (Figure 4H). The BSR domain stabilizes α6 and the C-terminal loop through hydrogen bonds formed between the side chains of Ser491 and Asn466, and between Arg485 and Ser476. Meanwhile, Glu631, Val633 and Gly635, located in the β3-β4 loop of the Inhibitory domain form hydrogen bonds with Ser460 in the β6-η3 loop, Tyr458 in β6 and with Gln462 in η3, respectively. The VWA-BRS interaction also involves η3, α1 and α2 in VWA, and the external side of the BRS first β-sheet. This interaction leads to burial of the above-mentioned hydrophobic patch in the BRS surface; a hydrophobic interaction core is formed by residues within the VWA helices, including Ile469 in α6 and Leu332 and Leu335 in α1. Additional hydrogen bonds involving α1 and α1-β2 loop with the β5-β6 loop (Gln328-Thr548) and β8 (Gln336-Thr580 and Thr337/Val338-Arg582) further contribute to interface stabilization.

In contrast, the direct interaction between the VWA and the Inhibitory domain is predominantly hydrophilic and significantly weaker. The external face of the second β-sheet of the Inhibitory domain interacts with η3, α6, β6, α5, β5, and their connecting loops. Asp600 in the Inhibitory domain β1’ forms a salt bridge with Lys450 in α5, while Arg670 establishes a salt bridge with Glu446, also in α5, and forms hydrogen bonds with Glu446 and Ala442 (Figure 4H). As mentioned earlier, Ala442 is located in the β5-α5 loop, in close proximity to the MIDAS motif and Glu414.

Thus, the BSR and the Inhibitory domains may together contribute to further inhibition of CLCA1 than the direct interaction between the MH and the ID; interactions between BSR, ID and VWA, specifically involving VWA η3 and α6, may prevent the VWA open-closed conformational change and ligand binding (discussed above).

### FnIII-like domain

The CLCA1 FnIII-l domain exhibits a structural arrangement similar to the Inhibitory domain, sharing the same β-strand connectivity, with the exception that β1 remains uninterrupted (Figure 5A-B).

**Figure 5:**
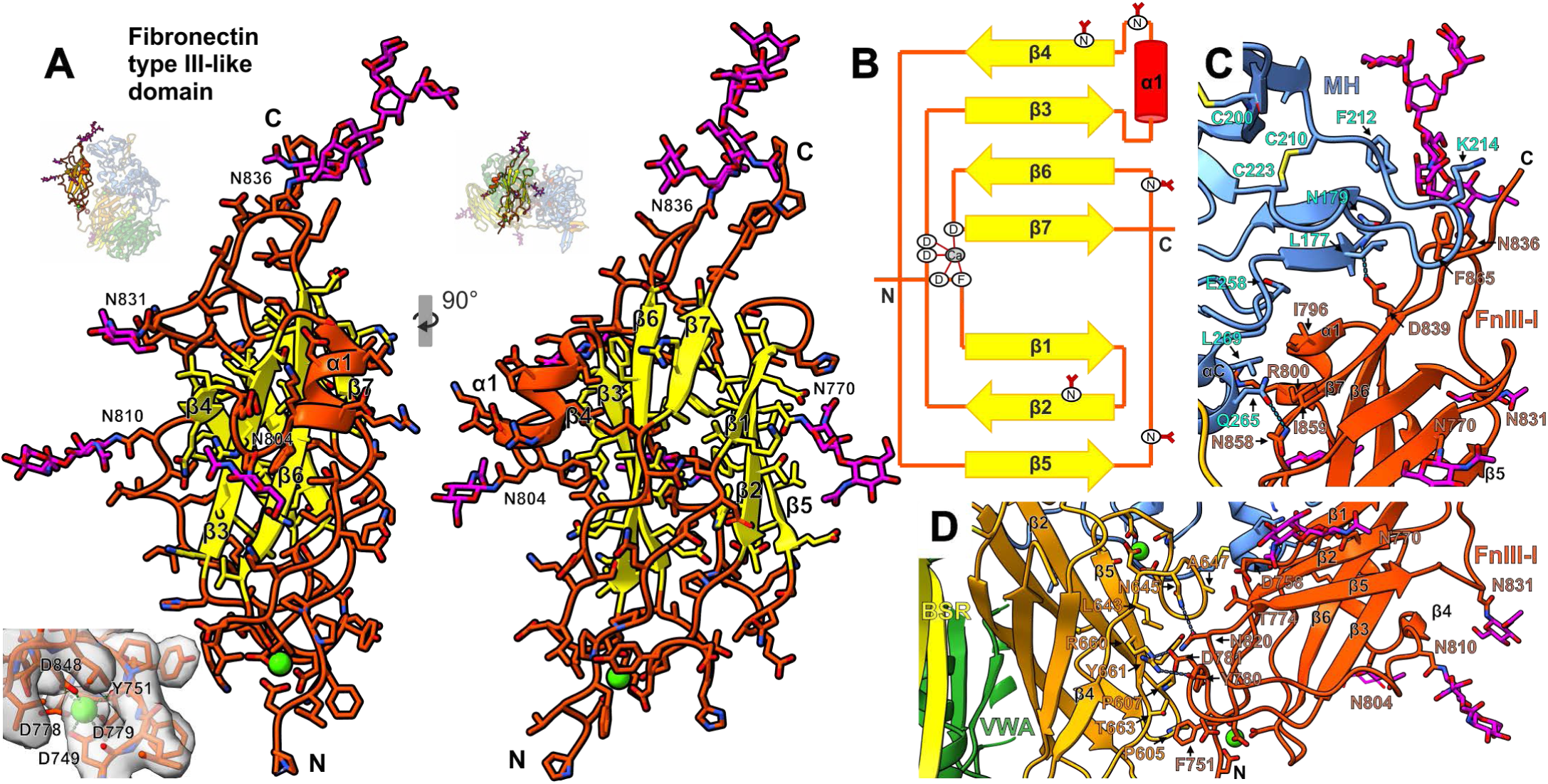
The Fibronectin type III-like (FnIII-l) domain of CLCA1. **A:** Cryo-EM model of the FnIII-l domain in cartoon and stick representation. β-strands are shown in yellow, α-helices and loops in red, calcium in green, and N-glycans in magenta. The domain is displayed in the same orientation as BSR and ID domains (Figure 4A and 4C) in two views, rotated 90° clockwise around the y-axis. *N*-glycosylated asparagine residues are labelled, as are the N- and C-termini and the secondary structure elements. The full CLCA1 molecule is shown next to each view to illustrate the relative position of the FnIII-l domain. MH (blue), VWA (green), BSR (yellow), ID (orange) and N-glycans (magenta). A close-up of the calcium-binding site is shown together with the cryo-EM map, with residues involved in coordination indicated. **B:** Schematic representation of the FnIII-l domain. *N*-glycosylated asparagine residues are marked. The same color code as in (A) is used. **C-D:** Detail of the interaction region between the MH (blue) and FnIII-l (red; N-glycans in magenta) domains (C), and between the ID (orange) and FnIII-l domains (D). Interacting residues are labelled and shown as sticks, as are the cysteines stabilizing the MH interacting loop. Hydrogen bonds are shown as dashed lines, and calcium ions are represented as green spheres.

These structural similarities suggest an evolutionary relationship between the two domains, which may have been overlooked due to the insertion between β6 and β7 in the ID.

The FnIII-l domain lacks the RGD (Arg-Gly-Asp) sequence, a characteristic motif found in FnIII domains involved in cell adhesion. In fact, its protein recognition capacity appears to be strongly limited by extensive *N*-glycosylation, a feature more commonly associated with Ig-like domains. It contains five *N*-glycans located at β2 (Asn770), the α1-β4 loop (Asn804), β4 (Asn810), and the β5-β6 loop (Asn831 and Asn836). *N*-glycans at Asn804, Asn810, and Asn831 project toward the solvent, whereas Asn770 and Asn836 orient toward the Inhibitory and MH domains, respectively (Figure 5 and S3).

The connecting loops are generally long and disordered, and include an ion-binding site at the bottom of the β-sandwich as well as an α-helix insertion between β3 and β4, which lies on the top of the external face of the second β-sheet (Figure 5A-B). The ion, presumably Ca²⁺, is coordinated by the Phe751 backbone carbonyl oxygen and by the carboxyl groups of four aspartic acid residues: Asp749 in the loop connecting ID and FnIII-l, Asp778 and Asp779 in the β2-β3 loop, and Asp848 in the β6-β7 loop. These first two regions are precisely involved in the interaction with the Inhibitory domain which in some CLCA1 orthologs results in the burial of a free cysteine at the equivalent position Gly754 (Figure S5). Additionally, the connecting loops at the bottom of the first β-sheet interact with the ion-binding loop (β4-β5) of the ID. The interactions are mainly hydrophobic and involve Phe751, Thr774 and Tyr780 in FNIII and Leu643, Ala647, Tyr661 and Thr663 in the Inhibitory domain. Interaction stabilization is mediated by hydrogen bonds between Asp781 and Arg660, and between Asn820 and Asn645, located in FnIII-l and the Inhibitory domain, respectively (Figure 5D).

FnIII-l interacts back with the MH domain, resulting in CLCA1’s characteristic compact structure. The interaction is also predominantly hydrophobic and involves two distinct regions (Figure 5C). The first region involves FnIII-l α1 and the β6-β7 loop, as well as MH αC, and is stabilized by a hydrogen bond between Asn858 and Gln265. The second region comprises the β5-β6 loop including the *N*-glycan at Asn836, the upper parts of β6 and β7, and the C-terminal loop of FnIII-l. It also includes the unusually large segment of the MH domain between αB and αC, specifically the β1*-β2* β-hairpin and the long β4*-β5* loop stabilized by the Cys200-Cys205 disulfide bond. Both regions adopt a conformation resembling opposing β-hairpins arranged in a clamp-like shape that accommodates FnIII-l in the central part. Only one hydrogen bond is observed between the side chain of Asp839 and the main chain of Asn179.

### CLCA1 activity

To investigate the regulatory mechanism of CLCA1 we aimed to express and enzymatically profile truncated variants of CLCA1. In the past, expression of truncated variants has been unsuccessful, probably due to incorrect folding (Nystrom et al. 2019). Based on the experimental structure we modified the domain borders to truncate CLCA1 after position 302, 478, 584, and 868. Of these, only CLCA1-868 was expressed as a stable construct. CLCA1-584 was secreted at very low levels, indicating a chaperone function of the CTP during protein folding in the endoplasmic reticulum. CLCA1-302 and 478 were both found to be highly unstable after purification, which might indicate self-degradation in the absence of the Inhibitory domain (data not shown).

Based on the similarity between CLCA1 and prokaryotic proteases able to degrade *O*-glycosylated proteins, we screened CLCA1 against several *O*-glycosylated substrates in the presence or absence of the sialidase BT0455, using StcE and OgpA as positive controls. As expected, StcE was able to cleave all substrates except IgA (fetuin only after desialylation) whereas OgpA was able to cleave all substrates (MUC2-N and IgA only after desialylation). However, no proteolytic activity was found for CLCA1 against either C1inh, Fetuin, IgA1 or the C-terminal end of MUC2 (MUC2-C) (Figure 6B, S7B-D). As expected, only very little CLCA1 activity was seen against recombinantly expressed and purified MUC2-N (Figure 6A). However, more profound degradation of N-terminal MUC2 was seen upon incubation with either human or murine semi-purified and denatured colonic MUC2 (Figure 6C-D). CLCA1 did not induce full degradation of human colonic MUC2 oligomers, as seen by SDS-AgPAGE analysis, whereas StcE fully dissolved these oligomers, indicating that CLCA1 might play a regulatory rather than disruptive role in MUC2 processing.

**Figure 6:**
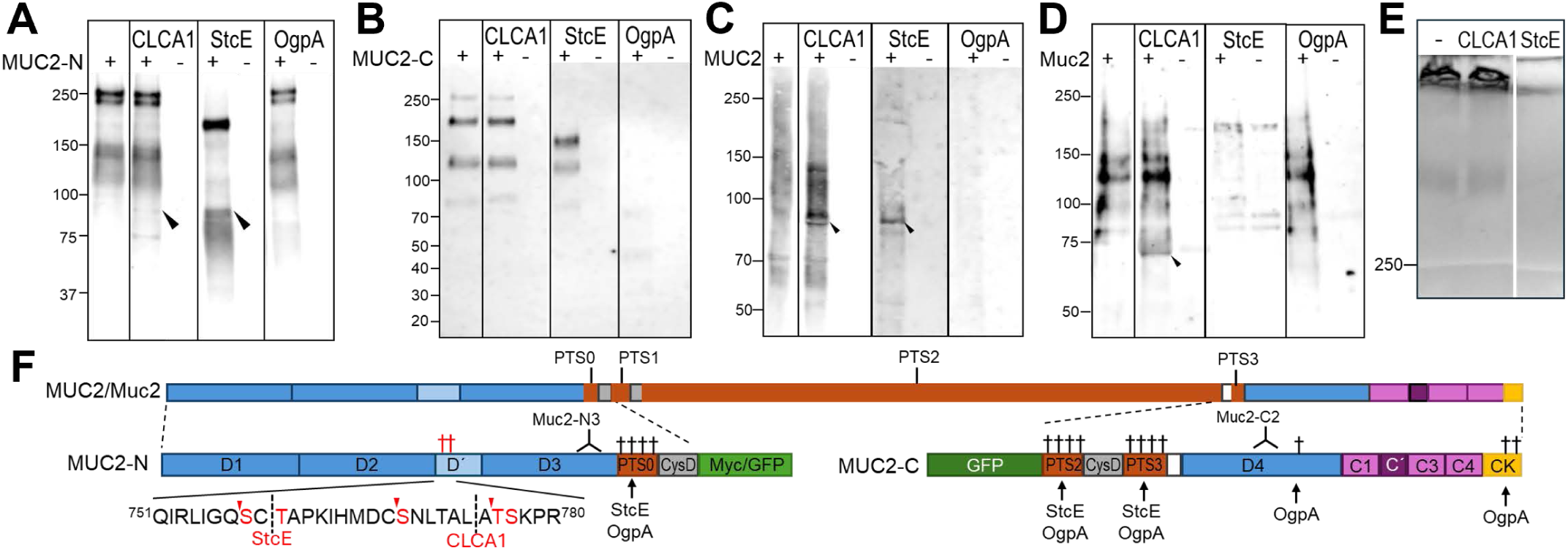
CLCA1, StcE and OgpA-mediated proteolysis of MUC2. **A-D:** Western blot analysis of MUC2-N (A) or MUC2-C (B) terminal constructs or semi-purified human (C) or murine (D) MUC2 after overnight incubation in the presence of CLCA1, StcE or OgpA as indicated. Protease and substrate only controls were included. Arrowheads indicate cleaved CLCA1 mediated or similar fragments. **E:** Alcian blue stained SDS-AgPAGE for analysis of non-reducible MUC2 oligomers from semi-purified human colonic mucus in the presence or absence of CLCA1 and StcE. **F**: Schematic representation of native Mucin-2 and terminal constructs used for proteolysis assays. Von Willebrand factor type D assemblies (D1-D4), the heavily *O*-glycosylated regions rich in proline, threonine and serine (PTS0-3), CIS domains, von Willebrand factor type C domains (C1-C4) and the cystic knot (CK) are indicated. † denotes known (black) or predicted (red) *O*-glycosylation sites. Red letters in the TIĹenlargement indicated *O*-glycosylated residues. Known (black) och suggested (red) cleavage regions or sites for StcE, OgpA and CLCA1 are marked. Red arrow heads in the TIĹ insert denoted predicted OgpA cleavage sites.

Noticeably, both for MUC2-N and human colonic MUC2, CLCA1 and StcE produced similar sized products. The observed N-terminal fragment was smaller than the expected product from StcE-mediated cleavage in the first PTS domain, indicating the existence of a glycosylated region in the central part of the MUC2 N-terminal region. Using the NetOGlyc prediction server (Steentoft et al. 2013), *O*-glycosylation was predicted at several sites in MUC2-N with high probability in the region between amino acid 769 to 786 which lie within the von Willebrand factor type D’ domain, specifically TIĹ (Table S4). Closer examination of this region revealed a potential consensus cleavage site for StcE at Gln757-Ser758 (Shon et al. 2021) and a previously suggested cleavage site of CLCA1 (Nystrom et al. 2019) between Leu774 and Ala775 with potential glycosylation of Thr776 and Ser777 in P2’ and P3’ respectively, in addition to several potential cleavage sites for OgpA (Figure 6F). Alignment analysis revealed that the suggested cleavage site for CLCA1 is poorly conserved both between orthologs and paralogs to MUC2 (Figure S7E). In rodent Muc2, a hydrophobic amino acid would be placed in P2’, consistent with the autocatalytic cleavage site. Thus, glycosylation in P2’ might be tolerated rather than required for CLCA1. However, the substrate and cleavage site for CLCA1 needs to be further deduced. *O*-glycan prediction of MUC2 orthologs did however reveal that the pattern with highly scored *O-*glycosylation prediction in the TIĹ domain is consistent between the different homologs.

Taken together, the data indicates that CLCA1 is able to cleave the N-terminal region in native (but denatured) MUC2, potentially in *O*-glycosylation dependent manner. Furthermore, *O*-glycosylation of N-terminal region of MUC2 might have a regulatory role for proteolysis, similar to what has been described for ADAMS (Goth et al. 2015).

## Discussion

By determining the structure of CLCA1 by cryo-EM we provide evidence that CLCA1 is a representative member of a novel eukaryotic family of metzincins closely related to the igalysins, which we propose to name CLCAsins. CLCA1 exhibits several distinctive features compared to classical metzincins, including a characteristic mode of coordinating a second Zn²⁺, the formation of a 3₁ knot that constitutes the first reported knotted metzincin domain, an extended β-sheet formed by the parallel insertion of a C-terminal β-strand, and a novel mechanism of zymogen formation. Instead of the classical pro-peptide regulation, CLCA1’s proteolytic activity appears to be inhibited by an extended loop from the Inhibitory domain that occludes the active site cleft, a mode of latent regulation distinct from the cysteine-switch typical of eukaryotic metzincins.

A striking parallel exists between CLCA1 and ADAMTS13, a metzincin that cleaves the von Willebrand factor (VWF) which is highly homologous in the terminal ends to MUC2 and other mucins representing the major protein component of the CLCA1 environment. Similar to CLCA1, ADAMTS13 lacks a functional propeptide; its short pro-sequence is dispensable for secretion and enzymatic activation (Majerus et al. 2003). Instead, ADAMTS13 is regulated through substrate-induced allosteric activation: under shear stress, binding of its substrate VWF to several exosites within the domains following the metalloprotease domain—including the Spacer domain, which is homologous to the BSR domain—triggers conformational changes that relieve autoinhibition and activate proteolysis (Petri et al. 2019). By analogy, we propose a similar mechanism in CLCA1, where binding of substrate via its VWA domain could induce conformational transitions in the BSR/Inhibitory domains that displace the inhibitory loop from the active site, thereby activating proteolytic function (Figure 7A).

**Figure 7:**
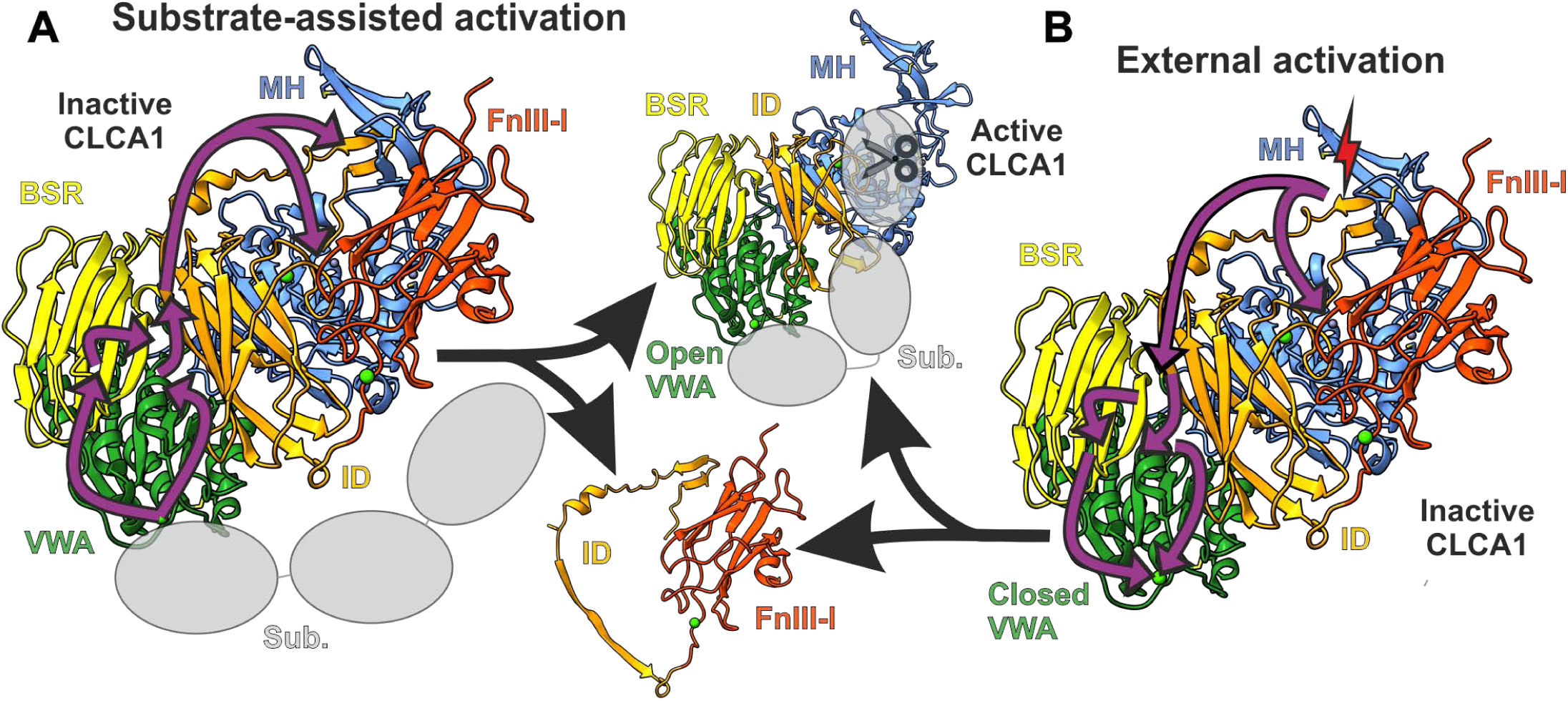
Models of CLCA1 allosteric activation. Substrate-assisted activation (left): binding of the VWA domain to a substrate induces the VWA open conformation, disrupting its interface with the BSR and ID domains and leading to the release of the active N-terminal part (NTP). External activation (right): proteolysis or redox events affect the ID domain, transmitting conformational changes to the VWA–BSR–ID interface and triggering the release of the active NTP. In this state, the VWA domain adopts an open conformation that can subsequently bind the substrate.

In an alternative model, removal of the BSD and Inhibitory domain precedes activation. The BSR and Inhibitory domains adopt a V-shaped conformation interacting with VWA α6. This is analogous to the lipase BlEst2 from *Bacillus licheniformis* which is autoinhibited by its C-terminal domains II and III and becomes activated upon proteolytic cleavage and removal of these domains (Nakamura et al. 2024) FEBS J.) (Figure S6C-F). Removal of BSR and ID would lead to the adoption of the VWA open conformation described in the previously reported crystallographic structure. This would in turn allow VWA to bind the substrate, while the MH domain, once released from the Inhibitory domain, would mediate its cleavage (Figure 7B). Although this is in line with our previous observation of a truncated N-terminal product of CLCA1 in intestinal mucus (Nystrom et al. 2019), an external cleavage between the VWA and the BSR, or between the BSR and the ID is unlikely without preceding conformational changes as the first linker is buried by the MH domain and the second is shielded by an *N*-glycan. Testing this model directly was impeded experimentally: truncated constructs of CLCA1 either failed to express or were unstable, likely due to intramolecular chaperone roles of the BSR and Inhibitory domains.

ADAMTS13 also regulates VWF non-enzymatically; free cysteines in the C-terminal CUB domains of ADAMTS13 reduces exposed cysteines in the VWC-domain of VWF, thereby preventing lateral disulfide bonds and oligomerization of VWF (Yeh et al. 2010). Given the high level of similarity between the terminal ends of VWF and MUC2 (Gallego et al. 2023), and our identification of partly reduced exposed cysteines in CLCA1 indicate that a similar mechanism might be possible between CLCA1 and MUC2. However, thiol-based interactions between CLCA1 and MUC2 needs further investigation.

Based on the homology between CLCA1/MUC2 and ADAMTS13/VWF, we propose a model wherein conformational alteration in MUC2, perhaps due to mucus dynamics or environmental shear, exposes a binding site to the CLCA1 VWA domain. Binding induces conformational changes in CLCA1, releasing its inhibitory loop (through displacement or proteolysis of BSR/Inhibitory domains), thereby activating the MH domain to cleave the substrate. Reversible disulfide bridges could modulate the CLCA1-MUC2 interaction, adding an additional layer of regulation.

The extended S2’pocket and similarities to bacterial *O-*glycoproteases further indicate that CLCA1 might act as a glycoprotease, adding yet another potential mechanism of substrate regulation. The features of the S2’binding pocket are largely conserved, but has slight variations that might relate to species specific glycosylation patterns of the substrates. However, compared to many bacterial *O-* glycoproteases with broad activity, CLCA1 appears to be very tightly regulated to be active only at the right site and the right time. However, substrate specificity and the mechanism for regulation needs further investigation.

The proposed involvement of CLCA1 in several diseases has raised the possibility to use CLCA1 as therapeutic target. Identification of its unique structural features will help directing development of specific compounds to alter the physiological function of CLCA1. Proteases are commonly targeted by compounds against the active site. However, the potential involvement of conformational changes in the VWA as a requirement for ligand binding poses and alternative approach, by which the VWA can be allosterically locked in its closed position as previously been suggested for integrins (Luo, Carman, and Springer 2007).

In conclusion, by determining the molecular structure of CLCA1 by cryo-EM we have identified unique features placing CLCA1 as a founding member of a novel family of metzincins, CLCAsins, which a potential novel mechanism of activity regulation. The distinctive features of CLCA1 described herein appears to be largely conserved throughout the CLCA family, providing evidence that CLCA1 can be considered a model protein for this family. Our findings can guide future research on development of therapeutic targeting of CLCA1 in mucus associated disease.

## Methods

### CLCA1 constructs

*CLCA1* (AF127036) in a pcDNA3.1 vector (Pawlowski et al. 2006) was used for expression of full-length CLCA1. Truncation and insertion of a His6-tag after Gly478 was introduced by QuikChange Site-directed mutagenesis kit (Agilent Technologies) according to manufacture’s instruction with 50 ng pcDNA3.1-CLCA1 as parental vector, and forward primer 5’-ggccctttcatcaggacatcatcaccatcaccactagtaatggagctgtctctc-3’ and reverse complementary primer. *CLCA1* (AF039400) truncated after amino acid 302, 584 and 868, with C-terminal His6-tags in pcDNA3.1 vectors were ordered from GenScript.

### Expression and Purification of recombinant proteins

CLCA1 vectors were transiently expressed in FreeStyle-CHO-S cells (RRID:CVCL_D604) by the Mammalian Protein Expression Facility at the University of Gothenburg and spent medium was collected at 68-96h post transfection. Cells and debris were removed by centrifugation and the remaining supernatant was dialyzed into 50 mM HEPES 150 mM NaCl (pH 7.4). CLCA1 truncates were purified by a combination of HisTrap EXCEL Ni Sepharose affinity chromatography (Cytiva) (CLCA1 302, 584 and 868) or PureCube Ni-INDIGO (Cure Biotech) (CLCA1 478), ion exchange chromatography and size exclusion chromatography. Ion exchange chromatography was performed on a Mono Q™ 5/50 GL anion-exchange column (Cytiva) at a flow rate of 1 mL/min using 50 mM HEPES, pH 7.4 as Buffer A and 500 mM NaCl, 50 mM HEPES, pH 7.4 as Buffer B. Samples were concentrated using VivaSpin 6 concentrators (3 kDa MWCO; Sartorius). For size-exclusion chromatography, a Superose 6 10/300 GL column (Cytiva) was used at a flow rate of 0.3–0.5 mL/min. All chromatographic procedures were performed using an ÄKTA Purifier system (GE Healthcare).

Full-length (FL) CLCA1 was expressed without any tag, and the spent medium was prepared for further purification in the same manner as described previously, but dialyzed into 50 mM HEPES, 20 mM NaCl, pH 8.0. FL_CLCA1 was first purified by anion-exchange chromatography using a HiPrep™ Q XL 16/10 column (Cytiva), then dialyzed into 50 mM HEPES, 20 mM NaCl, pH 7.4, and further purified on a Mono Q™ 5/50 GL anion-exchange column (Cytiva). In both chromatographic steps, bound proteins were eluted with a NaCl gradient from 20 mM to 500 mM. Fractions containing FL_CLCA1 as the major component were concentrated using VivaSpin 6 concentrators (3 kDa MWCO; Sartorius) and finally purified by size-exclusion chromatography on a Superose™ 6 10/300 GL column (Cytiva). *Bacteroides thetaiotamicron* sialidase BT0455 was expressed in *E. coli* strain TUNER (Novagen) and purified as described before (Ndeh et al. 2025). The recombinant protein was dialyzed into 10 mM MES, 200 mM NaCl, pH6.5. StcE was prepared as described in (Javerfelt et al. 2025).

### Mucus extraction

Mucus for negative stain electron microscopy was collected from co-housed and littermated *Clca1^+/+^* and *Clca1^-/-^*(Clca1^tm1Htzm^, RRID:MGI:3802573) on C57Bl/6N genetic background. Mucus for Muc2 isolation and proteolytic activity assay was prepared from C57bl/6N mice. All mice were kept in a SPF facility under standard conditions, in full compliance with Swedish animal welfare legislation (ethical permission 6269/24). After dissection, the colon was longitudinally opened, fecal pellets removed and mucus was scraped with a blunt spatula and collected into 150 µl Krebs-Ringer solution.

Human colonic tissue for mucus collection was obtained from brain-dead organ donors, at the end of the organ procurement procedure, in collaboration with Sahlgrenska University Hospital, Gothenburg, Sweden, in compliance with the human research ethical committee in Gothenburg, Sweden (ethical permit 2020-02677 with complement). After resection, the tissue was kept on ice. Mucus was collected by gentle scraping.

Insoluble Muc2 was extracted by incubating mucus samples in Extraction buffer (6 M guanidinium chloride, 5 mM EDTA, 10 mM NaH_2_PO_4_ pH 6.5) with (murine) or without (human) 1x cOmplete™ EDTA-free Protease Inhibitor Cocktail (Roche) at 4°C with gentle shaking overnight. The samples were spun at 17 000 g, 10 °C, 30 minutes and the supernatant discarded. The extraction was repeated once for 3 hours (murine mucus) or 8 hours (human mucus), after which the centrifugation was repeated. The resulting pellet was reduced for 2 hours (murine) or 5 hours (human) at 37°C in 200 µl reduction buffer (6 M GuHCl, 0.1 M Tris, 5 mM EDTA, pH 8.0) freshly supplemented with 10 mM (human) or 50 mM (murine) dithiothreitol (DTT). Reduction was quenched by addition of 2.5 times excess of iodoacetamide for 2.5 hours (murine) or overnight (human) at room temperature. The solution was centrifuged at 10,000 rpm, 4 °C for 30 min to remove any residual insoluble material. Finally, solubilized mucins were dialyzed (SnakeSkinTM 10 kDa dialysis tubing) against deionized water at 4 °C for 12 (murine) or 48 (human) hours, changing the water every 4-8 hours. Human mucins were lyophilized and prior to use resuspended in water (0.5% w/v). The protein concentration was determined by reading the absorbance at 280 nm using NanoDrop (ThermoFisher Scientific).

### Negative staining Electron Microscopy

Purified CLCA1, including full-length (FL_CLCA1) and truncated forms, was diluted to 20 µM in 50 mM HEPES, 150 mM NaCl, 5 mM CaCl₂, pH 7.4 immediately before grid preparation. Mucus samples were dissolved in the same buffer and normalized to a final protein concentration of 0.08 mg/mL, as determined by absorbance at 280 nm using a NanoDrop spectrophotometer. Carbon-coated copper grids (CF300-CU-50; Electron Microscopy Sciences) were glow-discharged for 40 s at 15 mA. A 5 µL aliquot of the protein solution was applied to each grid and incubated for 30 s. Excess sample was removed by blotting with filter paper, and grids were washed twice with distilled water. Grids were then stained with 2% (w/v) uranyl acetate for 30 s, blotted to remove excess stain, and air-dried at room temperature.

Micrographs were collected on a Thermo Fisher Talos L120C transmission electron microscope operated at 120 kV. Images were recorded at a nominal magnification of 73,000 x. Representative micrographs were used for qualitative assessment of particle integrity and distribution.

### Single particle cryo-EM

Full-length CLCA1 was diluted to a final concentration of 2 µM in 50 mM HEPES, 150 mM NaCl, 5 mM CaCl₂, pH 7.4. The sample was applied to Quantifoil Cu 1.2/1.3 300 mesh copper grids (SPT Labtech) that had been glow-discharged at 15 mA for 40 s with negative polarity using a GloCube Plus glow discharge system (Quorum). Grids were plunge-frozen in liquid ethane using a Vitrobot Mark IV (Thermo Fisher Scientific) operated at 100% humidity and 4 °C.

Data were collected on a Titan Krios transmission electron microscope (Thermo Fisher Scientific) operated at an acceleration voltage of 300 kV and equipped with a K3 Summit 6k × 4k direct electron detector (Gatan). Images were recorded in EPU (Thermo Fisher Scientific) using aberration-free image shift (AFIS) mode at a calibrated pixel size of 0.86 Å/pixel. Each movie consisted of 40 frames, with an average electron dose of ∼1.3 e⁻/Å² per frame and a total dose of approximately 50 e⁻/Å². The nominal defocus range was −0.5 to −2.5 µm.

Micrographs were imported into cryoSPARC v3.2 (Punjani et al. 2017), motion-corrected using Patch Motion Correction, and subjected to CTF estimation with Patch CTF. Micrographs were manually inspected, and poor-quality images were excluded prior to further analysis. Representative data collection and processing parameters are summarized in Supplementary Table S1.

A first round of manual particle picking was performed on representative micrographs, followed by 2D classification to select high-quality particle classes. These classes were then used as templates for automated particle picking. Extracted particles (box size: 256 pixels) were subjected to 2D classification, and poorly defined or junk particles were removed.

The remaining particles were used for ab initio reconstruction into four classes, followed by heterogeneous refinement. Particles from the best-resolved class were then subjected to a second ab initio reconstruction with two classes and subsequent heterogeneous refinement. The best class from this round underwent non-uniform refinement, followed by local refinement, global CTF refinement, another non-uniform refinement, and local refinement to improve map quality.

To further assess conformational variability, the refined particles were used in an additional heterogeneous refinement using the previously obtained high-quality model together with a lower-quality reference from the earlier two-class heterogeneous refinement. The best class from this step was subsequently refined through non-uniform refinement and a final local refinement imposing C2 symmetry.

The final FL-CLCA1 map was reconstructed from 337,181 particles and achieved an overall resolution of 2.99 Å, as determined by the gold-standard Fourier shell correlation (FSC) at 0.143.

### Model Building

An initial model of FL-CLCA1 was generated using AlphaFold2 (Jumper et al. 2021). The predicted model was fitted into the experimental cryo-EM density map using Molrep (Vagin and Teplyakov 2010) and manually adjusted in Coot v0.9.8.1 (Emsley et al. 2010). Iterative refinement was performed using the real-space refinement tool in Phenix v1.21.2-5419 (Liebschner et al. 2019) in combination with manual rebuilding in Coot. Model validation was carried out with MolProbity (Williams et al. 2018).

Structural analysis and figure preparation were performed using PyMOL v2.5 (Schrodinger 2015) and UCSF ChimeraX 1.9 (Meng et al. 2023). Extended *N*-glycans were added using GLYCAM-Web (Grant et al. 2025). The CLCA1 C-terminal region was modelled using AlphaFold3 (Abramson et al. 2024).

### Protolytic assays

1 uM CLCA1 or StcE, or 1U OgpA (opeRATOR, Genovis) was added to approximately 50 ug mucus in 25 mM HEPES, 75 mM NaCl and 2.5 mM CaCl2 and incubated at 37°C overnight.

Native IgA1 (Abcam, ab91020), recombinant human C1 inhibitor (Gibco, 17830493), fetuin from fetal bovine serum (Merck, F3385), MUC2-C (Trillo-Muyo et al. 2025) or MUC2-N (Ambort et al. 2012) was incubated with CLCA1, StcE or OgpA at 100nM (for both substrate and enzyme, 0.1 U for OgpA) in 50 mM HEPES, 150 mM NaCl, 5 mM CaCl_2_, with or without 0.1-1 µM BT0455, an unspecified sialidase from *Bacteroides thetaiotaomicron* (Briliute et al. 2019; Ndeh et al. 2025), at 37°C, overnight.

Samples were reduced in tris-glycine-SDS buffer containing 50 mM DTT for 20 minutes at 37°C before gel electrophoresis using Novex Tris-Glycine Gels (4-20% or 4-12%) (ThemoFisher Scientific). SYPRO^TM^ Ruby protein gel stain (Invitrogen) was used for in-gel protein detection using GelDoc^TM^ EZ Imager (Bio-Rad) for imaging. Gels with MUC2-C and MUC2-N were subsequently incubated in tris-glycine-SDS buffer for 20 min at room temperature and blotted to Immobilon® -FL PVDF membrane (Millipore) at 2mA/cm^2^, 1h. Gels with mucus samples were immediately blotted to Immobilon® -FL PVDF membranes after gel electrophoresis. After blocking with 2.5% milk in PBS, MUC2 fragments were visualized with αMUC2-C2 (Axelsson, Asker, and Hansson 1998) or αMUC2-N3 (Asker et al. 1998) followed by Alexa-680 conjugated α Rabbit antibodies. Odyssey CLx and Image Studio 6.0 (LIQOR Bio) was used for visualisation and image processing.

For MUC2 oligomer analysis, 75 ug MUC2 was incubated with 1 µM CLCA1 or StcE, or 1U OgpA at 37°C overnight. After reduction and denaturation, samples were analysed by SDS-AgPAGE prepared as described before (Issa et al. 2011) and stained with Alcian Blue.

### C-terminal dimerization assay

0.5 ug purified CLCA1 or CLCA1-868 were denatured in tris-glycine-SDS buffer with or without 50 mM DTT. Proteins were separated on a 4-20% Novex Tris-Glycine Gel, blotted to Immobilon® -FL PVDF membrane using Trans-Blot Turbo (Bio-Rad) and visualized using αCLCA1-1C4 (Abcam, ab129283) at 1:2000 dilution followed by an Alexa-680 conjugated α-mouse antibody.

### *O*-glycosylation prediction

For prediction of *O*-glycosylation sites in the N-terminal region of MUC2, the sequence for the MUC2-N construct (NP_002448.4 amino acid 1-1320) was submitted to the NetOGlyc 4.0 server (version 4.0.0.13) (Steentoft et al. 2013).

### Differential protein alkylation and mass spectrometry analysis

Free thiol groups of CLCA1 (in 50 mM HEPES, 200 mM NaCl, pH 7.4) were alkylated by incubation with 20 mM iodoacetamide (Sigma-Aldrich, I6125) for 30 minutes in the dark. In parallel fully alkylated and nonreduced control samples were prepared, the completely reduced sample was treated with 10 mM DTT and 5mM EDTA and alkylated with 40 mM iodoacetamide at 56°C for 60 minutes in the dark. The pH was subsequently adjusted to 5.5 with 0.2M phosphate buffer and subsequently denatured by the addition of SDS to a final concentration of 2% (v/v).

The samples were prepared for proteomics analysis by solvent precipitation on magnetic beads according to the SP3 method (Hughes et al. 2019). Protein aggregation on 1:1 combined Sera-Mag SpeedBeads (Cytiva, 45152105050250 and 65152105050250) was induced by adding ethanol to a final concentration of 50% (v/v), followed by incubation at room temperature for 10 minutes while shaking at 1000 rpm. The beads were then washed twice with 80% (v/v) ethanol. On bead reduction and alkylation was performed by adding 5 mM TCEP (Sigma-Aldrich, C4706), 20 mM iodoacetamide-d4 (Cambridge isotope laboratories, DLM-7249), 100mM Tris pH8 in 50% EtOH and incubated for 30 min at 37°C while mixing at 750 RPM. The beads were then washed twice with 80% (v/v) ethanol and on-bead protein digestion was performed using Trypsin/LysC protease mix (ThermoFisher, A41007) at a 1:20 enzyme-to-protein ratio in 25 mM ammonium bicarbonate for 16 hours at 37 °C. After digestion, the supernatant was collected using a magnetic rack, and acidified with trifluoroacetic acid (TFA) to a final concentration of 0.5% (v/v). The samples were desalted on C18 stagetips (Rappsilber, Mann, and Ishihama 2007) packed with 2 rounds punched from a Empore C18 filter (3M, 66883), lyophilized and resolved in 20 µL 0.1% TFA for mass spectrometry analysis.

Analysis was performed by online nanoflow liquid chromatography tandem mass spectrometry Q Exactive HF-X Orbitrap mass analyzer (Thermo Fisher Scientific) coupled to an EASY-nLC 1200 (Thermo Fisher Scientific). Mobile phase A was 0.1% formic acid (Fisher Chemical, A117) and B 0.1% formic acid in 80% acetonitrile. Peptides were separated using a 40 minute gradient up to 45% mobile phase A at 250 nL/min on a 20 cm column (i.d. 75 μm, Reprosil-Pur C18-AQ, 3 μm, 120 Å) packed inhouse. The mass spectrometer was operated at acquiring MS1 scans at 240.000 resolution (at 200 *m/z*) followed by 15 data-dependent MS2 scans at 15.000 by fragmentation by higher-energy collisional dissociation (HCD) with a normalized collision energy of 27%. Unassigned and singly charged precursors were excluded, and dynamic precursor exclusion was set to 60 seconds Target AGC for MS1 was 1e6 with a scan range from 280 - 1200 *m/z* and MS2 5e5 starting scanning from 100 *m/z*.

Data analysis was performed utilizing the FragPipe v21.1 (Polasky et al. 2020) pipeline using MSfragger v4.0 for protein identification searched against the human proteome (UniProt 2210, 80581 protein entries). Searches were restricted to strictly tryptic, with methionine oxidation and cysteine carboxyamidomethylation d0/d4 set as variable modification with a minimum peptide length set to 4 – 50 amino acid residues. Peptide and protein level FDR were set to 1%.

For quantification of cysteine site occupation, the ion intensities of the monoisotopic peak were extracted for each d0 and d4-labelled peptide using Xcalibur Qual browser v2.2. The overlap in monoisotopic peaks at the selected charge state was used to correct the intensity of the d4-labeled peptide based on the natural isotope distribution. Glycopeptides were identified using FragPipe in combination with PTM-shepherd with the additional modification search settings to include the human *N*-glycan (medium 253 entries) database and IonQuant v1.10 for LFQ based peptide quantification. Additional unknown cysteine modification were identified using the default open search settings in PTM-shepherd (Geiszler et al. 2021).

### DALI searches

The DALI protein structure comparison server was used for matching structural elements of CLCA1 against other proteins (Holm et al. 2023). For structural comparison of the MH, the structure corresponding to amino acid 22-301 was compared against the non-redundant PBD25 database (2025-aug-20). Protease clan was manually curated for the top 15 hits using the MEROPS peptidase database (Rawlings et al. 2018).

### Sequence alignment

To analyze conservation of features across the CLCA family, sequence alignment of Uniprot accession A8K7I4 (CLCA1_Human), Q6PT52 (CLCA1_Macaca mulatta), Q2TU62 (CLCA1_Horse), Q9TUB5 (Clca1_Pig), F1MGZ5 (Clca1_Bovin), A6HWA3 (Clca1_Rat), Q9D7Z6 (Clca1_Mouse), F6S0F1 (Clca1_Monodelphis domestica), A0A2U3V5T1 (Clca1_Tursiops truncatus), Q9UQC9 (Clca2_Human), Q9Y6N3 (Clca3_Human), Q14CN2 (Clca4_Human), Q8BG22 (Clca2_Mouse), Q9QX15 (Clca3a1_Mouse), Q6Q473 (Clca4a_Mouse), F6TR44 (Clca2_Horse), X1WHI8 (Clca1_Zebrafish), A0A8M9Q5D8 (Clca5.2_Zebrafish), A0A8M3APN3 (Clca5.2_Zebrafish) using Clustal Omega *via* Uniprot using standard parameters. Q02817 (MUC2_Human), Q80Z19 (MUC2_Mouse), Q62635 (MUC2_Rat), P98088 (MUC5AC_Human) and Q9HC84 (MUC5B_Human) were similarly used for alignment to analyze conservation of TIĹ domain residues.

## Acknowledgements and Funding

This work was supported by the Estonian Research Council (PUTJD1227, GR), Swedish Society for Medical Research (SSMF)(PD20-0168, EN) and the Swedish Research Council (2020-02536, SvdP; 2022-00646, ST-M).

Protein production was carried out at the Mammalian Protein Expression (MPE) core facility, and grid preparation for cryo-EM and NS-EM was performed at the Centre for Cellular Imaging (CCI). Both MPE and CCI are part of the Core Facilities of the University of Gothenburg. We acknowledge Protein Production Sweden (PPS) for providing facilities and experimental support for expression and purification of recombinant proteins. PPS is funded by the Swedish Research Council as a national research infrastructure. Cryo-EM data were collected at the Swedish National Cryo-EM Facility, funded by the Knut and Alice Wallenberg, Family Erling Persson, and Kempe Foundations, SciLifeLab. The plasmids for BT0455 and StcE expression were kindly provided by David N. Bolam and Stacy A. Malaker, respectively.

## Author Contributions

EN and ST-M initiated and planned the research. ST-M designed, performed, and analyzed the structural experiments, with assistance from EN and SvDP in the analysis. EN designed, performed, and analyzed the proteolytic assays, with assistance from DBB in the experimental work. SvdP designed, performed, and analyzed the mass spectrometry experiments. MO obtained human colon samples, which GR used to purify human MUC2. TP expressed and purified StcE, AL expressed and purified BT0455. EN, SvDP, and ST-M secured funding. EN and ST-M wrote the manuscript with input from all authors, who approved the final version of the text.

## Data availability

The structural data are deposited in the PDB and EMDB under accession codes 9R2T and 53539.

Mass spectrometry data and FragPipe search outputs are deposited in the ProteomeXchange Consortium via PRIDE under the identifier PXDXXXXX (Perez-Riverol et al. 2022).

