## Supplementary Figures for "Cryo-EM structure of CLCA1 identifies CLCA1 as a founding member of a novel metzincin family"

#### Supplementary figure legends

Supplementary figure 1: **CLCA1 oligomerization and cysteines.**

**A:** Size-exclusion chromatogram of full-length (FL) CLCA1 under non-reducing (red) or reducing (DTT, blue) conditions, and of CLCA1 truncated after residue 868. Elution volumes of protein standards and their corresponding molecular weights are indicated with vertical lines: thyroglobulin dimer (T×2), thyroglobulin (T), ferritin (F), aldolase (A), and conalbumin (C).

**B:** Western blot analysis of the C-terminal cleavage product of CLCA1 from full-length (FL) or truncated (868) CLCA1 in reducing or non-reducing conditions (DTT + and – respectively). * denotes cleaved CTP, and ¤ denotes residual uncleaved CLCA1.

**C:** AlphaFold3-predicted model of the C-terminal region of CLCA1 (residues 869–914). Cartoon representation colored by confidence according to the predicted local distance difference test (pLDDT) is shown in the top right. Surface representation colored by molecular lipophilicity potential (MLP) with *O*- and *N*-glycans displayed as sticks, rotated 45° and 90° clockwise around the y-axis. The dashed arrow indicates the position of the hydrophobic face of the α-helix.

**D:** Spectral counts of cysteine-containing peptides identified with either a cysteine thiol (Cys), a cysteine forming a disulfide adduct with free cysteine present in the culture medium (Cys–Cys), or a cysteine modified by glutathione (Cys–GSH). Insets show the resolved structure with density maps for the indicated cysteines; the contour level is set by root mean square deviation (rmsd) at 3.2 for strong density features and 6.4 for weak ones. Lines indicate cysteine disulfide pairs.

**E:** Cryo-EM model and density map details around the ID domain and inhibitory loop, showing a continuous but weak density between residues E715 and K724, with no discernible density downstream of V681 or around C727. Same contour levels as in (F) were used.

Supplementary figure 2: **Cryo-EM structure of CLCA1.**

**A:** Flowchart summary of Cryo-EM processing steps. The number of micrographs used and the number of particles remaining after each sorting step for the last iteration are specified.

**B:** Representative micrograph. Scale bar is shown.

**C:** Local resolution shown at two different contour levels. Central frontal view of CLCA1 and a view rotated 90° anticlockwise around the y-axis.

**D:** Fourier Shell Correlations (FSC) for the final density map.

**E:** Representative 2D classes.

**F:** Per-particle distribution over azimuth and elevation angles for the final density map.

**G:** Conical FSC Area Ratio (cFAR) evaluated with respect to 3072 viewing directions.

**H:** FSC (Model-map).

Supplementary figure 3: **CLCA1 *N*-glycosylation.**

**A:** Cryo-EM model surface representation colored by domains. The model is rotated three times by 90° counterclockwise around the y-axis (to the right), and by 90° and 180° counterclockwise around the x-axis (to the bottom). MH (blue), VWA (green), BSR (yellow), ID (orange), and FnIII-l (red). *N*-glycans at asparagine residues 503, 585, and 770 are shown as identified in the mass spectrometry analysis and are colored blue, purple, and pink, respectively. The remaining *N*-glycans are shown following the same pattern as N503 when no fucose is observed (N810 and N831) or as N585 when fucosylation is inferred from the cryo-EM density map (N804 and N836).

**B:** N-glycans identified by mass spectrometry at positions N503, N585, and N770. A schematic representation of the most abundant glycan at each position is shown.

**C:** Cryo-EM model and density map details at a 3.2 root mean square deviation (rmsd) contour level for all seven observed *N*-glycosylation sites. Modeled glycans, as well as those inferred from the existing density, are indicated.

Supplementary figure 4: **Metallohydrolase (MH) domain alignments**
**A:** Structures of CLCA1 and related metzincins shown in standard orientation and rotated 90° clockwise around the y-axis. CLCA1 is boxed in black; high-homology metzincins (ADAM17 (PDB ID: 2DDF)) and ADAMTS13 (PDB ID: 6QIG)) are shown in blue; mucin-like domain degrading metzincins (StcE (PDB ID: 3UJZ), CpaA (PDB ID: 6O38), OgpA (PDB ID: 6Z2O), and IgAse (PDB ID: 9I4Z)) in green; and metzincins coordinating a second zinc atom in grey (AmzA (PDB ID: 2X7M), BACOVA(BO)_00662 (PDB ID: 3P1V), and IgAse). Active-site and zinc-coordinating residues are shown as sticks. N- and C-termini and helices αB and αC are indicated. Asterisks mark the distinctive extra β-strand present in some families.
**B:** Detail of the catalytic zinc and second zinc coordination in different metzincins. Helices αB and αC, the third ligand of the catalytic zinc, the adjacent characteristic aromatic residue, and the residues coordinating the second zinc ion are indicated.
**C:** Relative position of the zinc ions. Alignment of the structures in (B), where the CLCA1 backbone is shown in grey, and the zinc ions of CLCA1, IgAse, BO_00663, and AmzA are shown in blue, cyan, orange, and pink, respectively. The catalytic zinc is marked with an asterisk.

Supplementary figure 5: **CLCA1 sequence alignments.**
Clustal Omega sequence alignment of CLCA1 paralogs and orthologs in selected species denoted by their UniProt accession number. Conservation of amino acids or features discussed in the text are highlighted. Solid highlight; conserved, framed highlight; similarity, pink; S1´-S3´-delimiting residues. Color marking of CLCA1_Human denotes distinct domains; red; signal sequence, blue; MH, green; VWA; yellow; BSR (light) and Inhibitory domain (dark), orange; FnIII-like. Numbers above C denotes their position in CLCA1_Human (black) and either their disulfide partner, free cysteine (-SH) or $Zn2^{+}$-coordination (grey). O; confirmed *O*-glycosylation sites, $; *N*-glycan positions (N890 predicted but not confirmed). NB: CLCA3_Human is a truncated pseudogene and not expressed in vivo.

Supplementary figure 6: **β-sheet rich related domains.**
**A:** Cryo-EM model of the BSR domain (yellow) aligned with the ADAMTS13 spacer domain (PDB ID: 6QIG; brown). The domain is displayed in three views, rotated twice to the right by 90° clockwise around the y-axis. N-glycosylated asparagine residues and attached glycans are shown as sticks. The N- and C-termini and secondary structure elements are labelled: black for common elements, yellow for the BSR domain, and brown for the spacer domain.
**B:** Schematic representation of the domains shown in (A). Same color code.
**C:** Cryo-EM model of the BSR domain (yellow) aligned with the BlEst2 domain II (PDB ID: 6WPX; pink). Representation as in (A).
**D:** Domain organization of BlEst2. Domains I, II, and III are shown in different shades of pink. The active site is boxed, and the Lid domains 1 and 2 (L1 and L2), located in the interfacing region, are indicated.
**E:** Alignment of the BSR (yellow) and ID (orange) domains from CLCA1 with domains II (pale pink) and III (purple) from BlEst2. The "V" shape is illustrated by dashed lines.
**F:** Addition of the VWA (green) and Domain I (pink) to the alignment shown in (E). The enlarged view of the interfacing region shows that VWA α-helix 6 (α6) occupies a position equivalent to L1, and α-helix 1 (α1) to L2.

Supplementary figure 7: **CLCA1, StcE and OgpA-mediated proteolysis of MUC2**
**A:** Western blot analysis of MUC2-N incubated with or without CLCA1, StcE and OgpA in the presence of BT0455 sialidase.
**B-D:** Gel based analysis of IgA1 (B), C1inh (C) or Fetuin (D) proteolysis by CLCA1, StcE and OgpA with or without BT0455 (right panels with BT0455 as indicated). *; protease.
E: Sequence alignment of MUC2 TIL´ region from human, mouse and rat, and MUC5AC and MUC5B from human. Amino acids with NetOGlyc prediction score >0.5 are highlighted in red. Arrow indicate the suggested CLCA1 mediated cleavage site in human MUC2. Sequence highlighted in light blue; TIL´, dark  blue; E´.

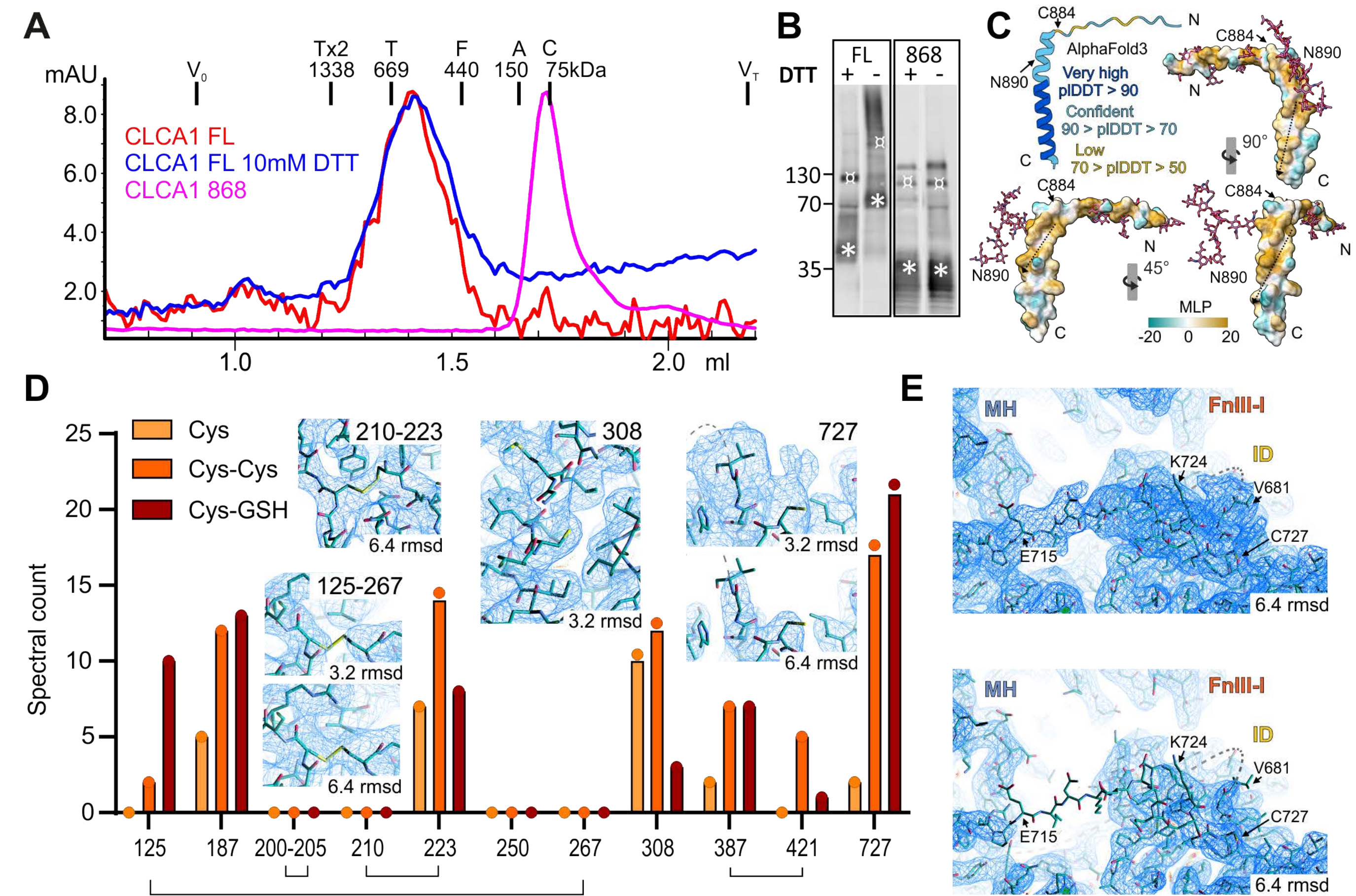

A
mAU
V0
Tx2 1338
T 669
F 440
A 150
C 75kDa
VT
CLCA1 FL
CLCA1 FL 10mM DTT
CLCA1 868
ml
B
FL
868
DTT
130
70
35
C
C884
N
N890
C
AlphaFold3
Very high pIDDT > 90
Confident 90 > pIDDT > 70
Low 70 > pIDDT > 50
90°
45°
MLP
-20 0 20
D
Cys
Cys-Cys
Cys-GSH
Spectral count
210-223
6.4 rmsd
308
3.2 rmsd
727
125-267
125
187
200-205
210
223
250
267
308
387
421
727
E
MH
FnIII-I
ID
K724
V681
C727
E715
6.4 rmsd

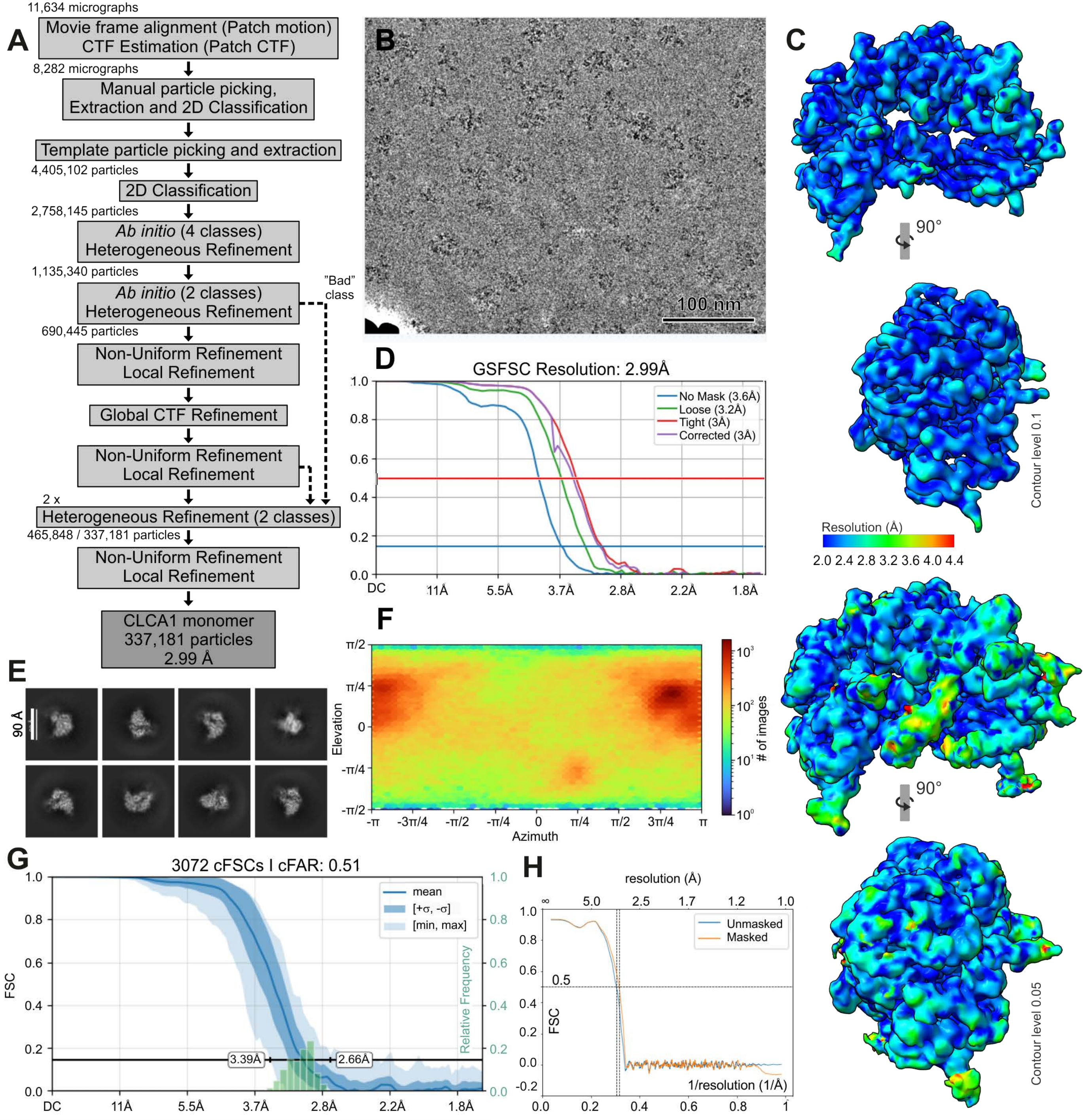

A
11,634 micrographs
Movie frame alignment (Patch motion)
CTF Estimation (Patch CTF)
8,282 micrographs
Manual particle picking, Extraction and 2D Classification
Template particle picking and extraction
4,405,102 particles
2D Classification
2,758,145 particles
Ab initio (4 classes) Heterogeneous Refinement
1,135,340 particles
Ab initio (2 classes) Heterogeneous Refinement
"Bad" class
690,445 particles
Non-Uniform Refinement Local Refinement
Global CTF Refinement
Non-Uniform Refinement Local Refinement
2 x
Heterogeneous Refinement (2 classes)
465,848 / 337,181 particles
Non-Uniform Refinement Local Refinement
CLCA1 monomer
337,181 particles
2.99 Å
B
100 nm
C
90°
Contour level 0.1
Resolution (Å)
2.0 2.4 2.8 3.2 3.6 4.0 4.4
90°
Contour level 0.05
D
GSFSC Resolution: 2.99Å
No Mask (3.6Å)
Loose (3.2Å)
Tight (3Å)
Corrected (3Å)
DC 11Å 5.5Å 3.7Å 2.8Å 2.2Å 1.8Å
E
90 Å
F
Elevation
Azimuth
### of images
G
3072 cFSCs | cFAR: 0.51
FSC
Relative Frequency
mean
[+σ, -σ]
[min, max]
3.39Å
2.66Å
DC 11Å 5.5Å 3.7Å 2.8Å 2.2Å 1.8Å
H
resolution (Å)
Unmasked
Masked
0.5
FSC
1/resolution (1/Å)

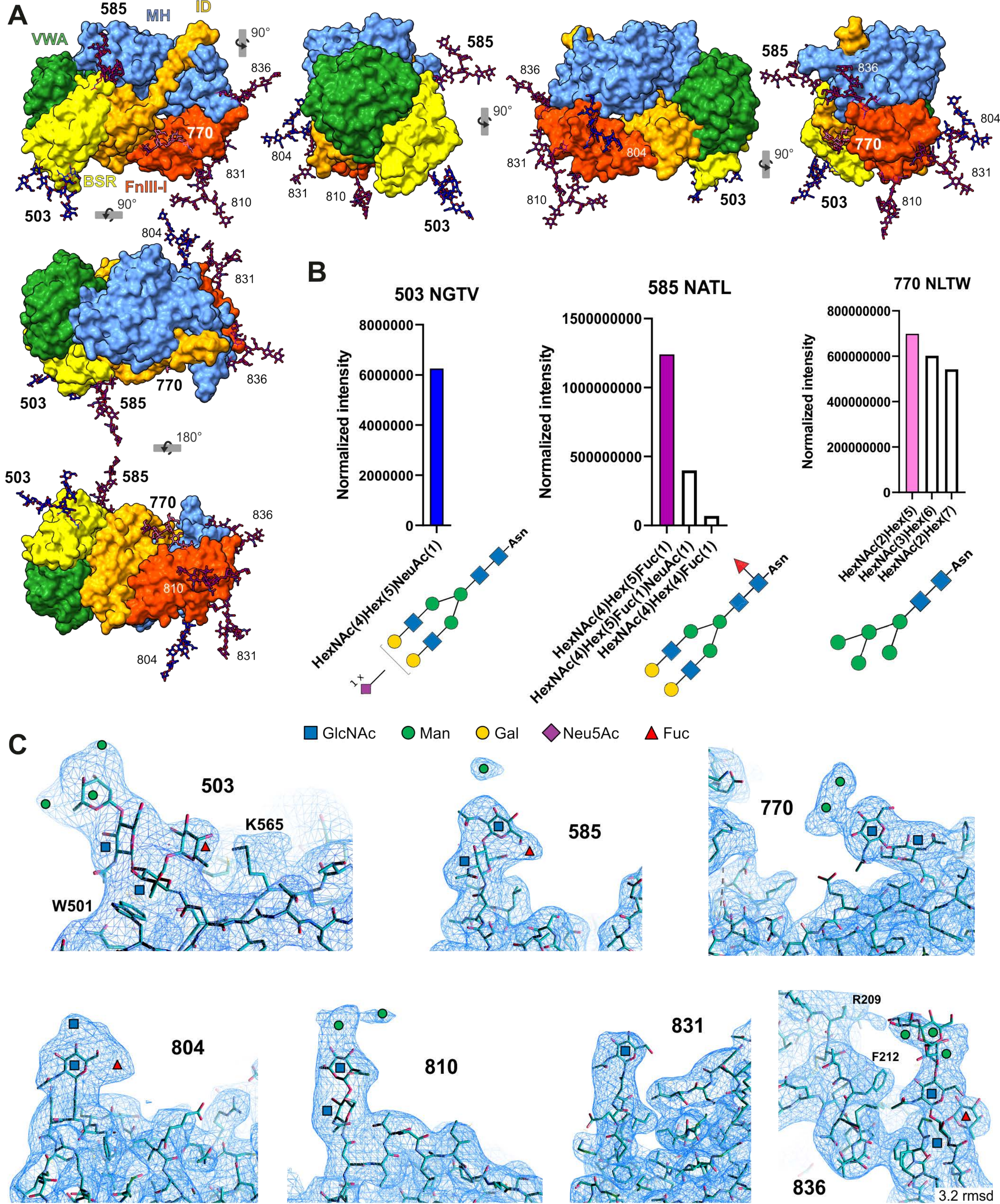
A
VWA
585
MH
ID
BSR
FnIII-I
770
503
836
831
810
804
90°
180°
B
503 NGTV
585 NATL
770 NLTW
Normalized intensity
HexNAc(4)Hex(5)NeuAc(1)
HexNAc(4)Hex(5)Fuc(1)
HexNAc(4)Hex(5)Fuc(1)NeuAc(1)
HexNAc(4)Hex(4)Fuc(1)
HexNAc(2)Hex(5)
HexNAc(3)Hex(6)
HexNAc(2)Hex(7)
Asn
1 x
GlcNAc
Man
Gal
Neu5Ac
Fuc
C
K565
W501
R209
F212
3.2 rmsd

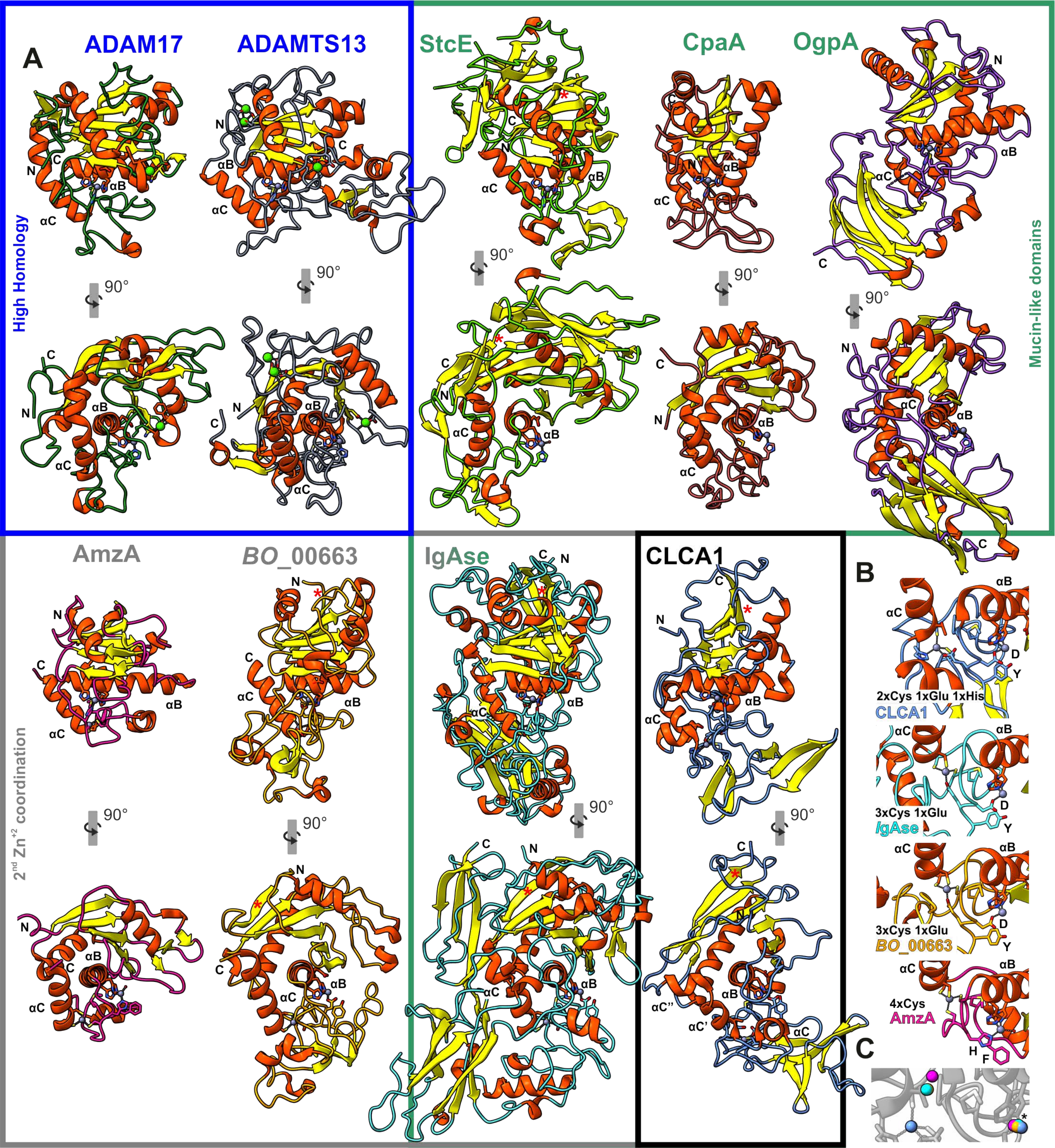
A
ADAM17
ADAMTS13
StcE
CpaA
OgpA
High Homology
Mucin-like domains
AmzA
BO_00663
IgAse
CLCA1
2nd Zn+2 coordination
B
2xCys 1xGlu 1xHis
CLCA1
3xCys 1xGlu
IgAse
3xCys 1xGlu
BO_00663
4xCys
AmzA
C
90°
αB
αC
αC'
αC''
N
C
D
Y
H
F

```
sp|A8K7I4|CLCA1_HUMAN              ------MGPFKSSVFILI--LHLLEGALSNSLIQLNNNGYEGIVVAIDPNVPEDETLIQQ  52
sp|Q6PT52|CLCA1_MACMU              ------MGPFKSSVFILI--LHLLEGALSDSLIQLNNNGYEGIVIAIDPNVPEDETLIQQ  52
sp|Q2TU62|CLCA1_HORSE              ------MGSFKSSVFILV--LHLLEGALSNSLIHLNNNGYEGIVIAIDPNVPEDETLIQQ  52
sp|Q9TUB5|CLCA1_PIG                ------MGSFRSSLFILV--LHLLEGAQSNSLIQLNGNGYEGIVIAIDPNVPEDERLIQN  52
tr|F1MGZ5|F1MGZ5_BOVIN             ------MGSFKNSVFILV--LHLLEGALSDSLIQLNNNGYEGIVIAIDPNVPEDETLIQH  52
tr|A6HWA3|A6HWA3_RAT               ------MGSLKSPVFLLV--LYLLEGVLSNSLIQLNNNGYEGIVIAIDHDVPEDEALIQR  52
sp|Q9D7Z6|CLCA1_MOUSE              ------MESLKSPVFLLI--LHLLEGVLSESLIQLNNNGYEGIVIAIDHDVPEDEALIQH  52
tr|F6S0F1|F6S0F1_MONDO             ------------------------------------------------------------  0
tr|A0A2U3V5T1|A0A2U3V5T1_TURTR     ------MGSFKNSVFILV--LHLLEGALSDSLIQLNNNGYEGIVIAVDPNVPEDETLIQQ  52
sp|Q9UQC9|CLCA2_HUMAN              MTQRSIAGPICNLKFVTLLVALSSELPFLGAGVQLQDNGYNGLLIAINPQVPENQNLISN  60
sp|Q9Y6N3|CLCA3_HUMAN              -------MVFSLKVILFL--SLLLSPVLKSSLVTLNNNGYDGIVIAINPSVPEDEKLIQN  51
sp|Q14CN2|CLCA4_HUMAN              ------MGLFRGFVFLL---VLCLLHQSNTSFIKLNNNGFEDIVIVIDPSVPEDEKIIEQ  51
sp|Q8BG22|CLCA2_MOUSE              MTHRDSTGPVIGLKLVTLLFTLSPELLFLGAGLKLKENGYDGLLVAINPRVPEDLKLITN  60
sp|Q9QX15|CA3A1_MOUSE              -------MVPGLQVLLFL--TLHLLQNTESSMVHLNSNGYEGVVIAINPSVPEDERLIPS  51
sp|Q6Q473|CLA4A_MOUSE              ------MAFSRGPVFLLL--LLYLLWGSDTSLIRLNENGYEDIIIAIDPAVPEDTTIIEH  52
tr|F6TR44|F6TR44_HORSE             MTHRDNEGLVCSLKFLILLVALSPELLSLGAGVQLQDNGYDGLLVAINPQVSEDQNLIPN  60
tr|X1WHI8|X1WHI8_DANRE             ------------MDWRTVFVVLWMLLPYPSTAIKLDGGGYVDITIAIGAKVKQDDTLIDK  48
tr|A0A8M9Q5D8|A0A8M9Q5D8_DANRE     ------------------------MLLSTSTGIKLDGDGYVDIIIAIGAKVSENNRLIDN  36
tr|A0A8M3APN3|A0A8M3APN3_DANRE     ------------MHLRSVFVLLWMLRLSTSTGIKLDGNGYIDITIAISSKVPEDDKLIDK  48

                                                                                 oo
sp|A8K7I4|CLCA1_HUMAN              IKDMVTQASLYLLEATGKRFYFKNVAILIPETWKTKADYVRPKLETYKNADVLVAESTPP  112
sp|Q6PT52|CLCA1_MACMU              IKDMVTQASPYLFEATGKRFYFKNVAILIPETWKTKADYVRPKLETYKNADVLVAESTPS  112
sp|Q2TU62|CLCA1_HORSE              IKDMVTQASPYLFEATEKRFYFKNVAILVPENWKTKPEYERPKLETYKNADVLVAEPNPP  112
sp|Q9TUB5|CLCA1_PIG                IKDMVTKASPYLFEATEKRFYFKNVAILIPASWKAKPEYVKPKLETYKNADVVVTEPNPP  112
tr|F1MGZ5|F1MGZ5_BOVIN             IKDMVTDASPYLFEATEKRFYFKNVAILIPENWKTKPEYVKPKLETYKNADVLVAEPNPT  112
tr|A6HWA3|A6HWA3_RAT               IKDMVTQASPYLFEATGKRFYFKNVAILIPENWNTKPEYKRPKLETLKNADVLVSTMSPI  112
sp|Q9D7Z6|CLCA1_MOUSE              IKDMVTQASPYLFEATGKRFYFKNVAILIPESWKAKPEYTRPKLETFKNADVLVSTTSPL  112
tr|F6S0F1|F6S0F1_MONDO             ---MVSEASNYLYGATEKRFYFKGVSILIPETWQKKPEYEKPKLETYKKADIVIDVPNPP  57
tr|A0A2U3V5T1|A0A2U3V5T1_TURTR     IKDMVTEASPYLFEATEKRFYFKNVAILIPQNWNTKPEYVKPKLETYKNADVLVAEHNPT  112
sp|Q9UQC9|CLCA2_HUMAN              IKEMITEASFYLFNATKRRVFFRNIKILIPATWKAN-NNSKIKQESYEKANVIVTDWYGA  119
sp|Q9Y6N3|CLCA3_HUMAN              IKEMVTEASTHLFHATKQRAYFRNVSILIPMTYKSKSEYLIPKQETYDQADVIVADLYLK  111
sp|Q14CN2|CLCA4_HUMAN              IEDMVTTASTYLFEATEKRFFFKNVSILIPENWKENPQYKRPKHENHKHADVIVAPPTLP  111
sp|Q8BG22|CLCA2_MOUSE              IKEMITEASFYLFNATKRRVFFRNVQILVPATWTDH-NYSRVRQESYDKANVIVAEQSEE  119
sp|Q9QX15|CA3A1_MOUSE              IKEMVTQASTYLFEASQGRVYFRNISILVPMTWKSKPEYLMPKRESYDKADVIVADPHLQ  111
sp|Q6Q473|CLA4A_MOUSE              IKGMVTKASTYLFEATEKRFFFKNVSILIPESWKDSPQYRRPKQESYKHADIKVAPPTVE  112
tr|F6TR44|F6TR44_HORSE             IKKMITEASFCLFNATKRRVFFRNIKILIPATWKAN-NYSKAKQESYEKANVIVTDWYGA  119
tr|X1WHI8|X1WHI8_DANRE             IKEMVTDGSFYLYHALDKKVYLKDATILVPSHWSCK-SCSKARTESFEKAKIKIDHAK--  105
tr|A0A8M9Q5D8|A0A8M9Q5D8_DANRE     IKDMISEGSFYLFHALDKKVYFKEATILVPSHWSCK-GCGKARTESFEKAKIKVDYA---  92
tr|A0A8M3APN3|A0A8M3APN3_DANRE     IKDMVTEGSVHLYKALDKKVYFKEVTILVPLNWNSK-GFTKARTELFETGMIRIDNPHPA  107
                                      *:: .*  *  *   : :::   **:*  :         : *  . . : :

                                               125-267
sp|A8K7I4|CLCA1_HUMAN              GNDEPYTEQMGNCGEKGERIHLTPDFIAGKKL-AEYGPQGRAFVHEWAHLRWGVFDEYNN  171
sp|Q6PT52|CLCA1_MACMU              GGDEPYTEHIGKCGDQGERIHLTPHFLAGKQL-KEYGPQGRAFVHEWAHLRWGVFDEYNN  171
sp|Q2TU62|CLCA1_HORSE              GNDQPYTEQMGKCGEKGERIYFTPDFLAGKRL-DEYGPQGRVFVHEWAHLRWGLFNEYND  171
sp|Q9TUB5|CLCA1_PIG                ENDGPYTEQMGNCGEKGEKIYFTPDFVAGKKV-LQYGPQGRVFVHEWAHLRWGVFNEYNN  171
tr|F1MGZ5|F1MGZ5_BOVIN             GNDGPYTEQMGGCGEKGDRIYLTPDFLAGKKS-RQYGPQGRAFVHEWAHLRWGVFNEYNN  171
tr|A6HWA3|A6HWA3_RAT               GNDEPYTEHIGACGERGIRIHLTPDFLAGKKQ-TEYGPQDRTFVHEWAHFRWGVFDEYNN  171
sp|Q9D7Z6|CLCA1_MOUSE              GNDEPYTEHIGACGEKGIRIHLTPDFLAGKKL-TQYGPQDRTFVHEWAHFRWGVFNEYNN  171
tr|F6S0F1|F6S0F1_MONDO             NNDAPRTDQFGQCGDKGERIHLTPDIILGKKL-KEYGPQGKILVHEWAHFRWGVFEEYNE  116
tr|A0A2U3V5T1|A0A2U3V5T1_TURTR     GNDGPYTEHMGGCGEKGNKIYLTPGFLAGKKL-LQYGPQGRVFVHEWAHLRWGVFNEYDN  171
sp|Q9UQC9|CLCA2_HUMAN              HGDDPYTLQYRGCGKEGKYIHFTPNFLLNDNLTAGYGSRGRVFVHEWAHLRWGVFDEYNN  179
sp|Q9Y6N3|CLCA3_HUMAN              YGDDPYTLQYGQCGDKGQYIHFTPNFLLTNNL-ATYGPRGKVFVHGWAHLRWGVFDEYNV  170
sp|Q14CN2|CLCA4_HUMAN              GRDEPYTKQFTECGEKGEYIHFTPDLLLGKKQ-NEYGPPGKLFVHEWAHLRWGVFDEYNE  170
sp|Q8BG22|CLCA2_MOUSE              HGDDPYTLQHRGCGQEGRYIHFTPSFLLNDELAAGYGARGRVFVHEWAHLRWGVFDEYNN  179
sp|Q9QX15|CA3A1_MOUSE              HGDDPYTLQYGQCGDRGQYIHFTPNFLLTDNL-RIYGPRGRVFVHEWAHLRWGVFDEYNV  170
sp|Q6Q473|CLA4A_MOUSE              GRDEPYTRQFTQCEEKAEYIHFTPDFVLGRKQ-DEYGDSGKVLVHEWAHLRWGVFDEYNE  171
tr|F6TR44|F6TR44_HORSE             HGDDPYTLQYRGCGKEGKYIHFTPDFLLNDDLTAGYGSRGKVFVHEWAHLRWGVFDEYNN  179
tr|X1WHI8|X1WHI8_DANRE             -LMEPRTKLYGECGKGGEYIHFTPDFLLNDSAIQMYGPRGKVFLHEWAHLRWGVYDEYNE  164
tr|A0A8M9Q5D8|A0A8M9Q5D8_DANRE     DSDDPYTNQYDECGKQGNYIHFTPNFLLSDTAVQTYGLRGKVFVHEWAHLRWGVYDETNE  152
tr|A0A8M3APN3|A0A8M3APN3_DANRE     YGDEPYTNQYGECGAEGQYIHFTPNFLRDDKLIRLYGSRGKVLVHEWAHLRWGVYDEYSE  167
                                       * *     *   .  *::** ::        **  .: ::* ***:***:::* .
```

```
                                                         187-Zn2+     200—205   210-223      223-201
sp|A8K7I4|CLCA1_HUMAN                  DEKFYLSN-GRIQAVRCSAGITGTNVVKKCQGGSCYTK-RCTFNKVTGLYEKGCEFVLQS   229
sp|Q6PT52|CLCA1_MACMU                  DEKFYLSN-GRIQAVRCSVGITGKIEVNKCQGGSCYTK-RCTFNKATGLYEKGCEFIPHS   229
sp|Q2TU62|CLCA1_HORSE                  DQKFYLSN-KEIKPVKCSADIAGKNVVNHCQGGSCATK-PCRRDRVTGLYAQECEFIPDE   229
sp|Q9TUB5|CLCA1_PIG                    EQKFYLSN-KKEQPVICSAAIRGTNVLPQCQGGSCVTK-PCRADRVTGLFQKECEFIPDP   229
tr|F1MGZ5|F1MGZ5_BOVIN                 DQKFYLSN-KRRKPVVCSEGFTGETVIQKCQGGSCVTQ-PCKLDRKTGLYEEGCEFIPHK   229
tr|A6HWA3|A6HWA3_RAT                   NEKFYLSN-GKPQAVRCSATITGKHVVRRCQGGSCVTNGKCVIDRVTGLYKDNCVFIPDK   230
sp|Q9D7Z6|CLCA1_MOUSE                  DEKFYLSK-GKPQAVRCSAAITGKNQVRRCQGGSCITNGKCVIDRVTGLYKDNCVFVPDP   230
tr|F6S0F1|F6S0F1_MONDO                 DEPFYQFN-GKNIPVKCSGAITGTNKVNACSGGSCSVR-NCRTDPQTGKLNKGCKFIPDE   174
tr|A0A2U3V5T1|A0A2U3V5T1_TURTR         DQKFYLSN-KRKKPVICPSTITGKNVLQKCQGGSCVTQ-SCKSDRVTGLYEKGCEFIPNE   229
sp|Q9UQC9|CLCA2_HUMAN                  DKPFYINGQNQIKVTRCSSDITGI---FVCEKGPCPQE-NCII---SKLFKEGCTFIYNS   232
sp|Q9Y6N3|CLCA3_HUMAN                  DQPFYISRRNTTEATRCSTRITVYMVLNECKGASCIAR-PFRRDSQTGLYEAKCTFIPKR   229
sp|Q14CN2|CLCA4_HUMAN                  DQPFYRAKSKKIEATRCSAGISGRNRVYKCQGGSCLSR-ACRIDSTTKLYGKDCQFFPDK   229
sp|Q8BG22|CLCA2_MOUSE                  DKPFYVNGRNEIQVTRCSSDITGV---FVCEKGLCPHE-DCII---SKIFREGCTFLYNS   232
sp|Q9QX15|CA3A1_MOUSE                  DQPFYMSRKNTIEATRCSTRITGTNVVHNCERGNCVTR-ACRRDSKTRLYEPKCTFIPDK   229
sp|Q6Q473|CLA4A_MOUSE                  DQPFYSASSKKIEATRCSTGITGTNRVYACQGGSCAMR-RCRTNSTTKLYEKDCQFFPDK   230
tr|F6TR44|F6TR44_HORSE                 EKPFYINGQNQIKVTRCSSDITGI---FVCEKGPCPQE-NCII---SKLFKEGCMFIYNS   232
tr|X1WHI8|X1WHI8_DANRE                 EKPFYLS-NGRVEYTRCTTNIEGQI--FEVNG-GSP-Q-SCRINPETFLPSSDCKFFPNK   218
tr|A0A8M9Q5D8|A0A8M9Q5D8_DANRE         NKAFYYS-NGRIEATRCSKNIEGQY--FSVSDGGSL-Q-TCQIDPITLLPDKTCKFFPNK   207
tr|A0A8M3APN3|A0A8M3APN3_DANRE         KTPFYRSASGKIEATRCSKNIEGQL--YDVTA-DPL-Q-PCQNDPQTSLPTDGCRFIPNK   222
                                       .  **          . *   :                  .         :   * *. .

                                                            250-Zn2+       267-125
sp|A8K7I4|CLCA1_HUMAN                  RQTEKASIMFAQHVDSIVEFCTEQNHNKEAPNKQNQKCNLRSTWEVIRDSE----DFKKT   285
sp|Q6PT52|CLCA1_MACMU                  QQTEKASIMFAQHVDSVVEFCTEQNHNKEAPNMQNTKCNLRSTWEVIRDSE----DFKKT   285
sp|Q2TU62|CLCA1_HORSE                  QQTEKASIMFSQSIDSVVEFCTEENHNREAPNMQNQRCNLRSTWEVIRDSE----DFKKT   285
sp|Q9TUB5|CLCA1_PIG                    QQSEKASIMFAQSIDTVVEFCKEKNHNKEAPNDQNQKCNLRSTWEVIQDSE----DFKKT   285
tr|F1MGZ5|F1MGZ5_BOVIN                 DQREKASIMYSQSIDTVVEFCTEKNHNREAPNEQNKKCNHRSTWEVIQDSE----DFKTT   285
tr|A6HWA3|A6HWA3_RAT                   NQREKASIMFNQNINSVVEFCTEKNHNKEAPNAQNQRCNLRSTWEVIQESE----DFKQT   286
sp|Q9D7Z6|CLCA1_MOUSE                  HQNEKASIMFNQNINSVVEFCTEKNHNQEAPNDQNQRCNLRSTWEVIQESE----DFKQT   286
tr|F6S0F1|F6S0F1_MONDO                 FQTEKASIMFMQSINSVVEFCTEKNHNKDAPNAHNQKCNLRSTWEVIKDSE----DYKKS   230
tr|A0A2U3V5T1|A0A2U3V5T1_TURTR         FQTEKASIMYSQSIRSVVEFCTEKNHNKEAPNEQNKKCNLRSTWEVIQGSE----DFKKT   285
sp|Q9UQC9|CLCA2_HUMAN                  TQNATASIMFMQSLSSVVEFCNASTHNQEAPNLQNQMCSLRSAWDVITDSA----DFHHS   288
sp|Q9Y6N3|CLCA3_HUMAN                  SQTAKESIVFMQNLDSVTEFCTEKTHNKEAPNL---------------------------   262
sp|Q14CN2|CLCA4_HUMAN                  VQTEKASIMFMQSIDSVVEFCNEKTHNQEAPSLQNIKCNFRSTWEVISNSE----DFKNT   285
sp|Q8BG22|CLCA2_MOUSE                  TQNATGSIMFMPSLPSVVEFCNESTHNQEAPNLQNQVCSLRSTWDVITASS----DLNHS   288
sp|Q9QX15|CA3A1_MOUSE                  IQTAGASIMFMQNLNSVVEFCTEKNHNAEAPNLQNKMCNRRSTWDVIKTSA----DFQNA   285
sp|Q6Q473|CLA4A_MOUSE                  VQSEKASIMFMQSIDSVTEFCKKENHNREAPTLHNKKCNYRSTWEVISTSE----DFNSS   286
tr|F6TR44|F6TR44_HORSE                 TQNATASIMFMQSLSSVVEFCNASTHNPEAPNLQNQMCSLRSAWDVITDSA----DFRHS   288
tr|X1WHI8|X1WHI8_DANRE                 DQNTDSSVMFSPSLEAVTTFCRETEHNYEAETQQNIICNNKATWTVIFEDSVDKDALSFL   278
tr|A0A8M9Q5D8|A0A8M9Q5D8_DANRE         NQNTDISLMFLPSLDSVTTFCRENDHNSEAPSQQNIKCGNDATWTTIFETSVDKNHLSSL   267
tr|A0A8M3APN3|A0A8M3APN3_DANRE         YQSTDSSMMSLPSLDSVITFCRESEHNYEAPNLQNAKCSNKATWTVIFEDSVDKDALRSL   282
                                        *    *::     : ::  **     ** :* .

                                                  MH β6     308-SH
sp|A8K7I4|CLCA1_HUMAN                  TPMT-T-QPPNPTFSLLQIGQRIVCLVLDKSGSMATGNRLNRLNQAGQLFLLQTVELGSW   343
sp|Q6PT52|CLCA1_MACMU                  TPTT-T-QPPNPTFSLLQIGQRIVCLVLDKSGSMATGNRLNRLNQAGQLFLLQIIELRSW   343
sp|Q2TU62|CLCA1_HORSE                  TPMT-A-QPPQPTFSLLQIGQRIVCLVLDKSGSMAIGDRLKRLTQAGKLFLLQTVEQGSW   343
sp|Q9TUB5|CLCA1_PIG                    TPMT-T-QPPAPTFSLLQIGQRIVCLVLDKSGSMTVGGRLKRLNQAGKLFLLQTVEQGAW   343
tr|F1MGZ5|F1MGZ5_BOVIN                 TPMT-T-QPPSPTFSLLQIGQRIVCLVLDKSGSMTIGNRLKRLNQAGKLFLLQTVEQGSW   343
tr|A6HWA3|A6HWA3_RAT                   TPMT-A-QPPAPTFSLLQTRQRIVCLVLDKSGSMQSDNRLNRMNQASRLFLLQTVEQGSW   344
sp|Q9D7Z6|CLCA1_MOUSE                  TPMT-A-QPPAPTFSLLQIGQRIVCLVLDKSGSMLNDDRLNRMNQASRLFLLQTVEQGSW   344
tr|F6S0F1|F6S0F1_MONDO                 NPMTAA-QPPKPTFSLKQIRERILVLVLDKSGSMASGDRLNRLNQASKLFLLQIIEKGSW   289
tr|A0A2U3V5T1|A0A2U3V5T1_TURTR         TPMT-T-QPPKPTFSLLQIRQRIVCLVLDKSGSMTTGDRLKRLNQAGKLFLLQTVEQGSW   343
sp|Q9UQC9|CLCA2_HUMAN                  FPMNGTELPPPPTFSLVQAGDKVVCLVLDVSSKMAEADRLLQLQQAAEFYLMQIVEIHTF   348
sp|Q9Y6N3|CLCA3_HUMAN                  ------------------------------------------------------------   262
sp|Q14CN2|CLCA4_HUMAN                  IPMV-T-PPPPPVFSLLKISQRIVCLVLDKSGSMGGKDRLNRMNQAAKHFLLQTVENGSW   343
sp|Q8BG22|CLCA2_MOUSE                  LPVHGVGLPAPPTFSLLQAGDRVVCLVIDVSRKMAEGDRLLRLQQAAELYLMQVVEAHTF   348
sp|Q9QX15|CA3A1_MOUSE                  PPMRGTEAPPPPTFSLLKSRRRVVCLVLDKSGSMDKEDRLIRMNQAAELYLTQIVEKESM   345
sp|Q6Q473|CLA4A_MOUSE                  TPME-T-SPSPPFFSLLRISERIMCLVLDVSGSMTSYDRLNRMNQAAKYFLSQIIENRSW   344
tr|F6TR44|F6TR44_HORSE                 SPMSETKLPPPPTFSLLQAGDKVVCLVLDVSSKMAEADRLLRLQQAAEFYLMQIVEIHTF   348
tr|X1WHI8|X1WHI8_DANRE                 PPL--PSTPSPPIFKLLQRKKRAVCLILDVSGSMATESRILRMRQAATHLLRNYVEEQAS   336
tr|A0A8M9Q5D8|A0A8M9Q5D8_DANRE         PSQD-LTHPPNTTFKVVQRTNRVLCLILDVSGSMH-GNRILLLQQAATLFLRTIIEEKAS   325
tr|A0A8M3APN3|A0A8M3APN3_DANRE         KPM--ASTPPAPNFKVVQRMQRTVCLVLDVSGSMK-GSRILRLQQASAHFLQNIIEDQSS   339
                                                *     *.: :   : : *::* * .*    .*:  : **.    *   :*  :
```

```
                                                                            387-421
sp|A8K7I4|CLCA1_HUMAN             VGMVTFDSAAHVQNELIQINSGSDRDTLAKRLPAAAS--GGTSICSGLRSAFTVIRK-KY 400
sp|Q6PT52|CLCA1_MACMU             VGMVTFDSAAHVQSELIQINSGSDRDTLTKRLPTAAS--GGTSICSGLRLAFTVIKK-KY 400
sp|Q2TU62|CLCA1_HORSE             VGMVTFDSAAYVQSALRQIKGGTDRDALTKSLPTVAS--GGTSICSGLRSAFTVIRK-KY 400
sp|Q9TUB5|CLCA1_PIG               VGMVAFDSAAYVKSELVQINSAAERDALARSLPTAAS--GGTSICSGLRSAFTVIKK-KY 400
tr|F1MGZ5|F1MGZ5_BOVIN            VGMVTFDSAAYVQSELVQINSATERDTLTKSLPTTAS--GGTSICSGLRSAFTVIKK-KY 400
tr|A6HWA3|A6HWA3_RAT              VGMVTFDSTAYVQSELTQLNSGADRDLLIKRLPTVAS--GGTSICSGLQAAFTSIKK-KY 401
sp|Q9D7Z6|CLCA1_MOUSE             VGMVTFDSAAYVQSELKQLNSGADRDLLIKHLPTVSA--GGTSICSGLRTAFTVIKK-KY 401
tr|F6S0F1|F6S0F1_MONDO            AGMVTFDSSATIQSELIQIETDAQRNSLITRLPTVAG--GGTSICSGLRTAFTVIKK-KF 346
tr|A0A2U3V5T1|A0A2U3V5T1_TURTR    VGMVTFDSAAYVKSELVQINSATERDTLARSLPTVAS--GGTSICSGIQSAFTVIRK-KY 400
sp|Q9UQC9|CLCA2_HUMAN             VGIASFDSKGEIRAQLHQINSNDDRKLLVSYLPTTVSAKTDISICSGLKKGFEVVEKLNG 408
sp|Q9Y6N3|CLCA3_HUMAN             ------------------------------------------------------------ 262
sp|Q14CN2|CLCA4_HUMAN             VGMVHFDSTATIVNKLIQIKSSDERNTLMAGLPTYPL--GGTSICSGIKYAFQVIGELHS 401
sp|Q8BG22|CLCA2_MOUSE             VGIVTFDSKGEIRASLQQIYSDDDRKLLVSYLPTAVSTDAETNICAGVKKGFEVVEERNG 408
sp|Q9QX15|CA3A1_MOUSE             VGLVTFDSAAHIQNYLIKITSSSDYQKITANLPQQAS--GGTSICHGLQAGFQAITSSDQ 403
sp|Q6Q473|CLA4A_MOUSE             VGMVHFSSQATIVHELIQINSDIERNQLLQTLPTSAN--GGTSICSGIKAAFQVFKNGEY 402
tr|F6TR44|F6TR44_HORSE            VGIASFNSKGEIRAQLHQINNDDDRKLLVSYLPATVSAEAETSICSGLKKGFEVVEKLNG 408
tr|X1WHI8|X1WHI8_DANRE            VGIVKFSTAASIVSSLTIIESDATRDHLINLLPETPG--GSTNMCNGLRLGLQVLSEDDM 394
tr|A0A8M9Q5D8|A0A8M9Q5D8_DANRE    VGIVTFSSQPNTLSQLTVIDSDDTREHLIRLLPTSAT--GSTEMCLGLQEGLKVLQGDNG 383
tr|A0A8M3APN3|A0A8M3APN3_DANRE    VALVTFSTDASTLSALTTIDKESTRENLVKLLPNVAE--GATNMCKGLNLGLQVLKTDNQ 397
                                  ..:. *.:       *  :      . :   **         .:* *:. .:  .   .
```

```
                                                       421-387
sp|A8K7I4|CLCA1_HUMAN             PTDGSEIVLLTDGED-NTISGCFNEVKQSGAIIHTVALGPSAAQELEELSKMTG---GLQ 456
sp|Q6PT52|CLCA1_MACMU             PTDGSEIVLLTDGED-NTISGCFNEVKQSGAIIHTVALGPSAARELEELSKMTG---GLQ 456
sp|Q2TU62|CLCA1_HORSE             KTDGSEIVLLTDGED-NTISSCFNEVKQSGAIIHTVALGPSAAAELEELSKMTG---GLQ 456
sp|Q9TUB5|CLCA1_PIG               PTDGSEIVLLTDGED-NTISACFPEVKQNGAIIHTVALGPSAAKELEELSQMTG---GLQ 456
tr|F1MGZ5|F1MGZ5_BOVIN            PTDGAEIILLTDGED-NTISACFDEVKQSGAIIHTVALGPSAAQELEQMSKMTG---GLQ 456
tr|A6HWA3|A6HWA3_RAT              PTDGAEIVLLTDGED-NTISSCFDLVKNSGAIIHTVALGPSAAKELEQLSKMTG---GLQ 457
sp|Q9D7Z6|CLCA1_MOUSE             PTDGSEIVLLTDGED-NTISSCFDLVKQSGAIIHTVALGPAAAKELEQLSKMTG---GLQ 457
tr|F6S0F1|F6S0F1_MONDO            STDGSEIVLLTDGED-NTISTCFDEVKQSGAIIHTVALGPSADPGLEKLAEMTG---GMK 402
tr|A0A2U3V5T1|A0A2U3V5T1_TURTR    PTDGSEIVLLTDGED-STISSCFNEVKQSGVIIHTVALGPSAARELEELSKMTG---GLQ 456
sp|Q9UQC9|CLCA2_HUMAN             KAYGSVMILVTSGDD-KLLGNCLPTVLSSGSTIHSIALGSSAAPNLEELSRLTG---GLK 464
sp|Q9Y6N3|CLCA3_HUMAN             ------------------------------------------------------------ 262
sp|Q14CN2|CLCA4_HUMAN             QLDGSEVLLLTDGED-NTASSCIDEVKQSGAIVHFIALGRAADEAVIEMSKITG---GSH 457
sp|Q8BG22|CLCA2_MOUSE             RADGSVLILVTSGAD-EHIANCLLTSMNSGSTIHSMALGSSAARKVGELSRLTG---GLK 464
sp|Q9QX15|CA3A1_MOUSE             STSGSEIVLLTDGED-NGIRSCFEAVSRSGAIIHTIALGPSAARELETLSDMTG---GLR 459
sp|Q6Q473|CLA4A_MOUSE             QTDGTEILLLSDGED-STAKDCIDEVKDSGSIVHFIALGPLADLAVTNMSILTG---GNH 458
tr|F6TR44|F6TR44_HORSE            KAHGSVMVLVTSGDD-EHVANCLLTVQSSGSTIHTIDLGSSAVENLEELSHLTGKYRGLK 467
tr|X1WHI8|X1WHI8_DANRE            DAIGDEIIFLTDGQATDDVTLCIPDAINSGAIIHTIALSDSAHNALQEMADKTG---GIF 451
tr|A0A8M9Q5D8|A0A8M9Q5D8_DANRE    DALGDEIIFLTDGQATDNIDACVPSAIQSGAIISTLAFSKDAVKALLVMAEQTG---GKF 440
tr|A0A8M3APN3|A0A8M3APN3_DANRE    DVLGDEIIFLTDGEATDTLSNCVPAAINSGAIIHTIALGDKADKALREMADKTG---GKF 454
                                     *  :::::.*   .    *.     .*  :  : :.  *   :  ::  **   *
```

```
                                                                                 $
sp|A8K7I4|CLCA1_HUMAN             TYASDQVQNNGLIDAFGALSSGNGAVSQRSIQLESKGLTLQNSQWMNGTVIVDSTVGKDT 516
sp|Q6PT52|CLCA1_MACMU             TYASDQVQNNGLIDAFGALSSGNGVVSERSIQLESKGSTLQNSQWMNGTVIVDSTVGKDT 516
sp|Q2TU62|CLCA1_HORSE             TYASDQAQNNGLIDAFGALSSGNGAVSQHSIQLVSRGSTLQNSQWMNGTVIVDSTVGNDT 516
sp|Q9TUB5|CLCA1_PIG               TYASDQAENNGLIDAFGALSSGNRAASQRSIQLESQGLTLQNNEWMNGTVVVDSTVGKDT 516
tr|F1MGZ5|F1MGZ5_BOVIN            TYASDQVQNNGLVDAFAALSSGNKAVSQRSIQLESRGLTLQNSQWMNGTVTVDSTVGKDT 516
tr|A6HWA3|A6HWA3_RAT              TYSSDQIQNNGLVDAFAALSSGNAAISQHSIQLESRGVNLQNKQWMNGSVIVDSTVGKDT 517
sp|Q9D7Z6|CLCA1_MOUSE             TYSSDQVQNNGLVDAFAALSSGNAAIAQHSIQLESRGVNLQNNQWMNGSVIVDSSVGKDT 517
tr|F6S0F1|F6S0F1_MONDO            TTATDNAQNNGLIDAFSALSSGNGAITQRSIQLVSKGATVKNGEWMNGTVIVDSTIGNDT 462
tr|A0A2U3V5T1|A0A2U3V5T1_TURTR    TYASDQVQNNGLMDAFGALSSGNGAVSQRSIQLESKGLTLQNSQWLNGTVIVDSTVGKDT 516
sp|Q9UQC9|CLCA2_HUMAN             FFVPDISNSNSMIDAFSRISSGTGDIFQQHIQLESTGENVKPHHQLKNTVTVDNTVGNDT 524
sp|Q9Y6N3|CLCA3_HUMAN             ------------------------------------------------------------ 262
sp|Q14CN2|CLCA4_HUMAN             FYVSDEAQNNGLIDAFGALTSGNTDLSQKSLQLESKGLTLNSNAWMNDTVIIDSTVGKDT 517
sp|Q8BG22|CLCA2_MOUSE             FFIPDKFTSNGMTEAFVRISSGTGDIFQQSLQVESVCETVQPQHQLADTMTVDSAVGNDT 524
sp|Q9QX15|CA3A1_MOUSE             FYANKD--LNSLIDAFSRISSTSGSVSQQALQLESKAFDVRAGAWINGTVPLDSTVGNDT 517
sp|Q6Q473|CLA4A_MOUSE             KLATDEAQNNGLIDAFGALASENADITQKSLQLESKGAILNNSLWLNDTVVIDSTLGRDT 518
tr|F6TR44|F6TR44_HORSE            FFVPDKSNSNSMIDAFSRISSGTGDIFQQHIQLESTGENVQPHYQLKNTVTMDNSVGNDT 527
tr|X1WHI8|X1WHI8_DANRE            FYSKDDFTSNQLMDAFASLTLSTGDHSNEPVQLESVGTR--TSDWFNGTVSVDQTVGNKT 509
tr|A0A8M9Q5D8|A0A8M9Q5D8_DANRE    ILSNDVISSNQLMDAFASLTLSTGDYTKEPVQLESAGTT--TSNWFNGTVSVDQTIGTKT 498
tr|A0A8M3APN3|A0A8M3APN3_DANRE    ITASDDVLSNQLLNGFSSLIMPTGNQTKDPVQIDSVGIK--TSDWFNGTVSVDQTIGTKT 512
                                      .    * : :.*  :   .    :  :*: *          : .:: :*.::* .*
```

```
sp|A8K7I4|CLCA1_HUMAN              LFLITWTMQP--PQILLWDPSGQKQGGFVVD---KNTKMAYLQIPGIAKVGTWKYSLQA-   570
sp|Q6PT52|CLCA1_MACMU              LFLVTWTTQP--PQILLWDPSGQKQDGFVVD---KNTKMAFLQIPGIAKVGTWKYSLQA-   570
sp|Q2TU62|CLCA1_HORSE              LFLITWTSQP--PQILLWDPSGKKQDGFVVD---TNTKMAYLQVPGTAKVGTWTYSLQA-   570
sp|Q9TUB5|CLCA1_PIG                LFLITLERKFLSPIPFFGVPSGRSQDSFLVG---KHNKMAYFQVPGTAKVGMWKYSLQA-   572
tr|F1MGZ5|F1MGZ5_BOVIN             LFLITWTTDL--PQILLWDPSGNKQDGFIVD---KNTKMAYLQIPDIAKIGVWKYSLQA-   570
tr|A6HWA3|A6HWA3_RAT               LFLVTWTTNS--PSIFIWDPSGVQQSGFVLD---TNTKVAYLQVPGIAKVGFWKYSIQA-   571
sp|Q9D7Z6|CLCA1_MOUSE              LFLITWTTHP--PTIFIWDPSGVEQNGFILD---TTTKVAYLQVPGTAKVGFWKYSIQA-   571
tr|F6S0F1|F6S0F1_MONDO             LFLVTWTAQQ--PQIFLSDPSGKTYNTFSVD---ANSKMAHLQIPNTAKVGMWRYSLKS-   516
tr|A0A2U3V5T1|A0A2U3V5T1_TURTR     LFLITWTTDI--PQILLWDPSGKKQDGFLVD---TYIKMAYLQIPGIAKVGIWKYSLQA-   570
sp|Q9UQC9|CLCA2_HUMAN              MFLVTWQASG-PPEIILFDPDGRKYYTNNFIT-NLTFRTASLWIPGTAKPGHWTYTLNNT   582
sp|Q9Y6N3|CLCA3_HUMAN              ------------------------------------------------------------   262
sp|Q14CN2|CLCA4_HUMAN              FFLITWNSLP--PSISLWDPSGTIMENFTVD---ATSKMAYLSIPGTAKVGTWAYNLQAK   572
sp|Q8BG22|CLCA2_MOUSE              LFLVTWQTGG-PPEIALLDPSGRKYNTGDFII-NLAFRTASLKIPGTAKHGHWTYTLNNT   582
sp|Q9QX15|CA3A1_MOUSE              FFVITWMVKK--PEIILQDPKGKKYTTSDFQDDKLNIRSARLQIPGTAETGTWTYSITGT   575
sp|Q6Q473|CLA4A_MOUSE              FFLVTWSKQA--PAIYLRDPKGTQTTNFTMD---SASKMAYLSIPGTAQVGVWTYNLEAK   573
tr|F6TR44|F6TR44_HORSE             AFLVTWQTSG-PPEIVLADPSGRKYYTDDFVT-NLTLQTARLRIPGTAKPGLWMYTLNNT   585
tr|X1WHI8|X1WHI8_DANRE             SFVIIYERIF--PAVYIQSPSGIVYTEAQMNHD-NRLKTVTLKVAETAEPGDWAYSIKTS   566
tr|A0A8M9Q5D8|A0A8M9Q5D8_DANRE     SFTITYETKL--PRISIQSPSGLIYSQTQMHHD-EPAKTVTLKVPGTAQTGDWKYSILSP   555
tr|A0A8M3APN3|A0A8M3APN3_DANRE     SFTITYETKL--PRISIQSPSGLTYSQTQMRHD-EPAKTVTLKVPGTAQTGDWKYSILSP   569
                                    * :        *    :   *.*         .         : . : :    *: * * *.:

                                                  $
sp|A8K7I4|CLCA1_HUMAN              --SSQTLTLTVTSRASNATLPPITVTSKTNKDTSKFPSPLVVYANIRQGASPILRASVTA   628
sp|Q6PT52|CLCA1_MACMU              --SSQTLTLTVTSRASSATLPPITVTSKMNKDTGKFPSPMIVYANIRQGASPILRASVTA   628
sp|Q2TU62|CLCA1_HORSE              --SSQTLTLTVTSRASSATLPPVTVTSKVNKDTGKFPSPVVVYAKIHQGGLPILRATVTA   628
sp|Q9TUB5|CLCA1_PIG                --SSQTLTLTVSSRRSSATLPPVTVTSKMNKDTGKFPSPMVVYTKIHQGTLPILRAKVTA   630
tr|F1MGZ5|F1MGZ5_BOVIN             --SSQTLTLTVTSRASSATLPPVTVTSKMNKDTGKFPSPMVVYVKIHQGTKPILRSKVTA   628
tr|A6HWA3|A6HWA3_RAT               --SSQTLTLTVTSRAASATLPPITVTPVVNKNTGKFPSPVTVYASIRQGASPILRASVTA   629
sp|Q9D7Z6|CLCA1_MOUSE              --SSQTLTLTVTSRAASATLPPITVTPVVNKNTGKFPSPVTVYASIRQGASPILRASVTA   629
tr|F6S0F1|F6S0F1_MONDO             --NAQTLTLTVTSRAANPTVPPITVDSKMNKDTSSFPSPMIVYAEVRQGSLPIIGADVTA   574
tr|A0A2U3V5T1|A0A2U3V5T1_TURTR     --SSQTLTLTVTSRASSGTLPPVTVTSKMNKDTGKFPSPMVVYAKIHQGSLPILRAKVTA   628
sp|Q9UQC9|CLCA2_HUMAN              HHSLQALKVTVTSRASNSAVPPATVEAFVERDSLHFPHPVMIYANVKQGFYPILNATVTA   642
sp|Q9Y6N3|CLCA3_HUMAN              ------------------------------------------------------------   262
sp|Q14CN2|CLCA4_HUMAN              -ANPETLTITVTSRAANSSVPPITVNAKMNKDVNSFPSPMIVYAEILQGYVPVLGANVTA   631
sp|Q8BG22|CLCA2_MOUSE              HHSPQALKVTVASRASSLAMSPATLEAFVERDSTYFPQPVIIYANVRKGLHPILNATVVA   642
sp|Q9QX15|CA3A1_MOUSE              --KSQLITMTVTTRARSPTMEPLLATAHMSQSTAQYPSRMIVYARVSQGFLPVLGANVTA   633
sp|Q6Q473|CLA4A_MOUSE              -ENSEILTITVTSRAANSSVPPITVNAKVNTDTNTFPSPMIVYAEVLQGYTPIIGARVTA   632
tr|F6TR44|F6TR44_HORSE             HDSLQALKVTVTSRASRSAVPPATVEAFVERDSTRFPHPMMIYANVRKGFYPILNATVTA   645
tr|X1WHI8|X1WHI8_DANRE             --TDQAFTITVTSQASHEDVPPIIVKTRMNQQVSDGTKPMTVFAEVSQNYRPVINVEVWA   624
tr|A0A8M9Q5D8|A0A8M9Q5D8_DANRE     --TLQSLTVTATSQAARANVPPITVKAQMNQQFSDGSKPMIVFAEVVQNYKPLIKAEVWA   613
tr|A0A8M3APN3|A0A8M3APN3_DANRE     --TLQSLTVTATSQAARANVPPITVKAQMNQQFSDGSKPMIVFAEVVQNYKPLIKAEVWA   627
                                      . : :.:*.:::      : *         . .        : ::. : :.   *::    * *

sp|A8K7I4|CLCA1_HUMAN              LIESVNGKTVTLELLDNGAGADATKDDGVYSRYFTTYDTNGRYSVKVRALGGVNAARRRV   688
sp|Q6PT52|CLCA1_MACMU              LIESENGKTVTLELLDNGAGADAAKDDGVYSRYFTTYDTNGRYSVKVRALGGVNAVRRRA   688
sp|Q2TU62|CLCA1_HORSE              LIESVDGKTVTLELLDNGAGADATKDDGIYSRYFTAYNTNGRYSIKVWALGGVNAARQMG   688
sp|Q9TUB5|CLCA1_PIG                LIESENGKTVTLELLDNGAGADATKNDGIYSRYFTAYDANGRYSVKVWALGGVNTPRRRA   690
tr|F1MGZ5|F1MGZ5_BOVIN             LIESVDGKTVTLELLDNGAGADATKDDGIYSRYFTAYDTNGRYSAKVWALGGVNTASQNA   688
tr|A6HWA3|A6HWA3_RAT               LIESVNGKTVTLELLDNGAGADATKNDGVYSRFFTAFDANGRYSIKIWALGGVTADRQRM   689
sp|Q9D7Z6|CLCA1_MOUSE              LIESVNGKTVTLELLDNGAGADATKNDGVYSRFFTAFDANGRYSVKIWALGGVTSDRQRA   689
tr|F6S0F1|F6S0F1_MONDO             LIESADGTTVTLELLDNGAGADTSKNDGVYSRYFTAYKINGRYSLKVTALGGTNT--QRS   632
tr|A0A2U3V5T1|A0A2U3V5T1_TURTR     LIESVDGKTVTLELLDNGAGADATKDDGVYSRYFTAYDTNGRYSVKVWALGGVDTATQKG   688
sp|Q9UQC9|CLCA2_HUMAN              TVEPETGDPVTLRLLDDGAGADVIKNDGIYSRYFFSFAANGRYSLKVHVNHSPSISTPAH   702
sp|Q9Y6N3|CLCA3_HUMAN              ------------------------------------------------------------   262
sp|Q14CN2|CLCA4_HUMAN              FIESQNGHTEVLELLDNGAGADSFKNDGVYSRYFTAYTENGRYSLKVRAHGGANTARLKL   691
sp|Q8BG22|CLCA2_MOUSE              TVEPEAGDPVVLQLLDGGAGADVIRNDGIYSRYFSSFAVSGSYSLTVHVRHSPSTSTLAL   702
sp|Q9QX15|CA3A1_MOUSE              LIEAEHGHQVTLELWDNGAGADTVKNDGIYTRYFTDYHGNGRYSLKVRVQAQRNKTRLS-   692
sp|Q6Q473|CLA4A_MOUSE              TIESNSGKTEELVLLDNGAGADAFKDDGVYSRFFTAYSVNGRYSLKVRADGGRNSARRSL   692
tr|F6TR44|F6TR44_HORSE             TIEPEAGDPVMLKLFDDGAGADVIKNDGIYSRYFFSFAVNGRYSLKVHVRHSPSMSSLTH   705
tr|X1WHI8|X1WHI8_DANRE             TLEPATGPIQSLQLLDNGAGADVMANDGIYSRYFTKM-VNGRCSLKVRAKNQDGKARFA-   682
tr|A0A8M9Q5D8|A0A8M9Q5D8_DANRE     TLESESGTVHELQLLDNGAGADTFKDDGVYSRYFTRM-KKGRSSLKVRVKNKDGQARFT-   671
tr|A0A8M3APN3|A0A8M3APN3_DANRE     TLESESGTVHELQLLDNGAGADTFKDDGVYSRYFTRM-KKGRSSLKVRVKNKDGQARFT-   685
                                    :*    *     * * *.*****    :**:*:*:*       .*   *  .: .
```

```
                                      Autocatalytic
                                      cleavage site
                                            ▼                              727-SH
sp|A8K7I4|CLCA1_HUMAN           IPQQSGALYIPGWIE-NDEIQWNPPRPEINKDDVQHKQVCFSRTSSGGSFVASDVPN-AP 746
sp|Q6PT52|CLCA1_MACMU           IPQQSGVMYIPGWIE-NDEIQWNPPRPEIE-DDVQRKQVCFSRTSSGGSFVASGVPN-AP 745
sp|Q2TU62|CLCA1_HORSE           ISQQNGAMYRAGRMK-NGEIIWNPPRPKINIDDLLSKQVPFSRTSSGGSFVASNVPN-AP 746
sp|Q9TUB5|CLCA1_PIG             PPLWSGAMYIRGWIE-NGEIKWNPPRPDINKDDLQGKQVCFSRTASGGSFVASDVPK-SP 748
tr|F1MGZ5|F1MGZ5_BOVIN          SPQRNGAMYIPGWIE-NGEVKWNPPRPEINKD--QGKQVCFSRTSSGGSFVATDVPK-AP 744
tr|A6HWA3|A6HWA3_RAT            APQRNGVMYIDGWIEDDGEIKMNPPHPETG--NVQDSQVCFSRTSSGGSFVATNVPA-AP 746
sp|Q9D7Z6|CLCA1_MOUSE           APPKNRAMYIDGWIE-DGEVRMNPPRPETS--YVQDKQLCFSRTSSGGSFVATNVPAAAP 746
tr|F6S0F1|F6S0F1_MONDO          SPQVNGAFYIPGWVE-NGDIKMNPPKPDFI-SDLQGKQVCFSRTSSGGSFVLSNIPS-GS 689
tr|A0A2U3V5T1|A0A2U3V5T1_TURTR  IPQQTRAMYIPGWIE-NGQVKWNPPRPEINKDDLQGKQVCFSRTSSGGSFVASDVPK-AP 746
sp|Q9UQC9|CLCA2_HUMAN           SIPGSHAMYVPGYTA-NGNIQMNAPRKSVGRNE-EERKWGFSRVSSGGSFSVLGVPA-GP 759
sp|Q9Y6N3|CLCA3_HUMAN           ------------------------------------------------------------ 262
sp|Q14CN2|CLCA4_HUMAN           RPPLNRAAYIPGWVV-NGEIEANPPRPEIDE-DTQTTLEDFSRTASGGAFVVSQVPS-LP 748
sp|Q8BG22|CLCA2_MOUSE           PVPGNHAMYVPGYIT-NDNIQMNAPKN-LGHRP-VKERWGFSRVSSGGSFSVLGVPD-GP 758
sp|Q9QX15|CA3A1_MOUSE           LRQKNKSLYIPGYVE-NGKIVLNPPRPDVQEEAIEATVEDFNRVTSGGSFTVSGAPPDGD 751
sp|Q6Q473|CLA4A_MOUSE           RHPSSRAAYIPGWVV-DGEIQGNPPRPETTE-ATQPVLENFSRTASGGAFVVSNVPS-GP 749
tr|F6TR44|F6TR44_HORSE          SVPGSHAMYVPGYIA-NGNIQMNAPRTSVGRSE-EEQKWGFSRISSGGSFSVLGVPA-GP 762
tr|X1WHI8|X1WHI8_DANRE          VQKRSGAPYVPGYVV-DGVVELNPPNPSVSDE--PIEVGSFTRTATGESFEVTLT---GS 736
tr|A0A8M9Q5D8|A0A8M9Q5D8_DANRE  LQKRGGAPYVPGYVV-NGVVELNPPKLSVSEE--LPEVGSFTRTATGESFEVSLT---SS 725
tr|A0A8M3APN3|A0A8M3APN3_DANRE  LQKRGGAPYVPGYVV-NGVVELNPPKLSVSEE--LPEVGSFTRTATGESFEVSLT---SS 739
                                        *  *    :. :  * *.              *.* ::* :*

                                                       $                                 $
sp|A8K7I4|CLCA1_HUMAN           IPDLFPPGQITDLKAEIHGGSLINLTWTAPGDDYDHGTAHKYIIRISTSILDLRDKFNES 806
sp|Q6PT52|CLCA1_MACMU           IPDLFPPCQITDLKAEIHGHSLINLTWTAPGDDYDHGTAHKYIIRISTSILDLRDKFNES 805
sp|Q2TU62|CLCA1_HORSE           IPDLFPPCQITDLKAKIQGDSLINLTWTAPGDDYDHGRADRYIIRISTNILDLRDKFNDS 806
sp|Q9TUB5|CLCA1_PIG             IPDLFPPCKITDLKAGIQGDNLINLTWTAPGDDYDHGRADRYIIRISTNILDLRDKFNDS 808
tr|F1MGZ5|F1MGZ5_BOVIN          IPDLFPPCKITDLKAKIEGDNLINLTWTAPGDDYDHGRADKYIIRISTNILELRDKFNES 804
tr|A6HWA3|A6HWA3_RAT            IPDLFPPCQITDLKASIQGQNLVNLTWTAPGDDYDHGRASSYIIRISTSIVDLRNNFSTS 806
sp|Q9D7Z6|CLCA1_MOUSE           IPDLFPPCQITDLKASIQGQNLVNLTWTAPGDDYDHGRASNYIIRMSTSIVDLRDHFNTS 806
tr|F6S0F1|F6S0F1_MONDO          IPDVFPPCQITDLKAQMDGN-HINLTWTAPGDDYDNGQAEQYIIKMSENILDLRDKFDDA 748
tr|A0A2U3V5T1|A0A2U3V5T1_TURTR  IPDLFPPCKITDLKAKIQGDSLINLTWTAPGDDYDHGRAHKYIIGISTNILDLRDKFNDS 806
sp|Q9UQC9|CLCA2_HUMAN           HPDVFPPCKIIDLEAVKVEE-ELTLSWTAPGEDFDQGQATSYEIRMSKSLQNIQDDFNNA 818
sp|Q9Y6N3|CLCA3_HUMAN           ------------------------------------------------------------ 262
sp|Q14CN2|CLCA4_HUMAN           LPDQYPPSQITDLDATVHED-KIILTWTAPGDNFDVGKVQRYIIRISASILDLRDSFDDA 807
sp|Q8BG22|CLCA2_MOUSE           HPDMFPPCKITDLEAMKVED-DVVLSWTAPGEDFDQGQTTSYEIRMSRSLWNIRDDFDNA 817
sp|Q9QX15|CA3A1_MOUSE           HARVFPPSKVTDLEAEFIGD-YIHLTWTAPGKVLDNGRAHRYIIRMSQHPLDLQEDFNNA 810
sp|Q6Q473|CLA4A_MOUSE           LPDLYPPNQITDLQATLDGE-EISLTWTAPGDDYDVGRVQQYIIRTSKNIIELRDNFNNS 808
tr|F6TR44|F6TR44_HORSE          HTDVFPPCKIIDLEAVKVED-QVTLSWTAPGEDFDEGQANSYEIRMSKSLQNIQDDFNNA 821
tr|X1WHI8|X1WHI8_DANRE          TPPNFPPSKVTDLGAKILEE-AVLLSWTAPGEDLDQGRAKSYEIRWSFDLDMLRQDFSNG 795
tr|A0A8M9Q5D8|A0A8M9Q5D8_DANRE  KPPKFPPCKIIDLSAEIQED-AVLLSWTAPGEDLDEGSAKSYDIRWSFEVKDLLFNFSNA 784
tr|A0A8M3APN3|A0A8M3APN3_DANRE  KPPKFPPCKIIDLSAEIQED-AVLLSWTAPGEDLDEGSAKSYDIRWSFEVKDLLFNFSNA 798
                                    :** :: ** *       : *:*****.  * * .  * *  *     :   *. .

                                   $                    $              $
sp|A8K7I4|CLCA1_HUMAN           LQVNTTALIPKEANSEEVFLFKPENITFE----------NGTDLFIAIQAVDKVDLKSEI 856
sp|Q6PT52|CLCA1_MACMU           LQVNTTALIPKEANSEEVFLFKPENITFE----------NGTDLFIAIQAVDKVDLKSEI 855
sp|Q2TU62|CLCA1_HORSE           LQVNTTDLIPKEATSEEVFVFKPENIAFE----------NGTDLFIAIQAVDEVDLESEI 856
sp|Q9TUB5|CLCA1_PIG             VQVNTTDLIPKEANSEEVFVFKPEGIPFT----------NGTDLFIAVQAVDKTNLKSEI 858
tr|F1MGZ5|F1MGZ5_BOVIN          LQVNTTDLIPNEANSEEVFVFKPETITFT----------NGTDLFIAIQAVDEVSLKSEI 854
tr|A6HWA3|A6HWA3_RAT            LEVNTTDLIPKEASSEEMFEFELD-NSFG----------NGTDVFIAIQAVDKSNLKSEI 855
sp|Q9D7Z6|CLCA1_MOUSE           LQVNTTGLIPKEASSEEIFEFELGGNTFG----------NGTDIFIAIQAVDKSNLKSEI 856
tr|F6S0F1|F6S0F1_MONDO          LQVNTSDLIPEKANSQEFFIFEPKDINLV----------NGTNIYIAIQAVDKANLKSSI 798
tr|A0A2U3V5T1|A0A2U3V5T1_TURTR  LQVNTTHLSPKEANSEEVFVFKPENITFT----------NGTDLFIAVQAVDKVNLKSEI 856
sp|Q9UQC9|CLCA2_HUMAN           ILVNTSKRNPQQAGIREIFTFSPQISTNGPEHQPNGETHESHRIYVAIRAMDRNSLQSAV 878
sp|Q9Y6N3|CLCA3_HUMAN           ------------------------------------------------------------ 262
sp|Q14CN2|CLCA4_HUMAN           LQVNTTDLSPKEANSKESFAFKPENISEE----------NATHIFIAIKSIDKSNLTSKV 857
sp|Q8BG22|CLCA2_MOUSE           ILVNSSELVPQHAGTRETFTFSPKLVTHELDHELAEDAQEPYIVYVALRAMDRSSLRSAV 877
sp|Q9QX15|CA3A1_MOUSE           TLVNASSLIPKEAGSKETFKFKPETFKIA----------NGIQLYIAIQADNEASLTSEV 860
sp|Q6Q473|CLA4A_MOUSE           PRVDTTNLTPKEANSEETFAFKPENITEE----------NATYIFIAIESVDKSSLSSGP 858
tr|F6TR44|F6TR44_HORSE          ILVNTSKLTPQQAGTKEIFTFSPKLFTEGPKYQPEGEAQENHRIYVAIRAIDRNSLKSAV 881
tr|X1WHI8|X1WHI8_DANRE          HVVDTATVSPQEAGSVEQHAFNLSL-----------PIQNGTTLFFAVQTLDEHDAKSDT 844
tr|A0A8M9Q5D8|A0A8M9Q5D8_DANRE  HVVNTSVISPQVFGSVEQHLFNLSF-----------PIQNGTILFFAIQSEDNEKLKSEM 833
tr|A0A8M3APN3|A0A8M3APN3_DANRE  HVVNTSVISPQVFGSVEQHLFNLSF-----------PIQNGTILFFAIQSEDNEKLKSEM 847
                                  *:::   *:     * . *.                 :   ::.*:.: :. .  *
```

```
                                                                   "PTS"        884-884 $      Hydrophobic
                                                                                               α-helix
sp|A8K7I4|CLCA1_HUMAN                 SNIARVSLFIPPQTPPETPSP-DETSAPCPNIHINSTIPGIHILKIMWKWIGELQLSIA-   914
sp|Q6PT52|CLCA1_MACMU                 SNIARVSLFIPPQTPPETPSP-DETSAPCPNISINSTIPGIHILKIMWKWIGELQLSIG-   913
sp|Q2TU62|CLCA1_HORSE                 SNIAQVSLFLPPQT----PPETPSPSLPCPDSNITSAIPGIHILKVMWKWLGELQLSVAL   912
sp|Q9TUB5|CLCA1_PIG                   SNIAQVSLFLPPEAPPETPPETPAPSLPCPEIQVNSTIPGIHILKIMWKWLGELQLSIA-   917
tr|F1MGZ5|F1MGZ5_BOVIN                SNIAQVSLFIPPEI----PPEKPSPSLPCPDISINSTIPGIHILKIMWKWLGELQISLG-   909
tr|A6HWA3|A6HWA3_RAT                  SNIARVSLFIPAQEPP------EDSTPSYPEVSINSTIPGIHVLNIVWKWLGEMHVTLGL   909
sp|Q9D7Z6|CLCA1_MOUSE                 SNIARVSVFIPAQEPPIP----EDSTPPCPDISINSTIPGIHVLKIMWKWLGEMQVTLGL   912
tr|F6S0F1|F6S0F1_MONDO                SNIAQVSVFIPPEETP-------------------CAGNNIFKIVLKWLGELQLTIPK   837
tr|A0A2U3V5T1|A0A2U3V5T1_TURTR        SNIAQVSLFIPPET----PPETPSPSLPCPDINTNSTIPGIHILKIMWKWLGELQLSVAL   912
sp|Q9UQC9|CLCA2_HUMAN                 SNIAQAPLFIPPNSDPVP-------------------ARDYLILKGVLTAMGLIGIICLI   919
sp|Q9Y6N3|CLCA3_HUMAN                 ------------------------------------------------------------   262
sp|Q14CN2|CLCA4_HUMAN                 SNIAQVTLFIPQANPDDIDPTPTPTPTPDKSH---NSGVNISTLVLSVIGSVVIVNFI   914
sp|Q8BG22|CLCA2_MOUSE                 SNIALVSMSLPPNSSPVV-------------------SRDDLILKGVLTTVGLIAILCLI   918
sp|Q9QX15|CA3A1_MOUSE                 SNIAQAVKLTSLEDSIS--------------------ALGDDISAISMTIWGLTVIFNSI   900
sp|Q6Q473|CLA4A_MOUSE                 SNIAQVALFTPQAEPDPDE------------SPS---SSGVSVATIVLSVLGALVLVCII   903
tr|F6TR44|F6TR44_HORSE                SNIAQASLFIPPNSAPVL-------------------ARDRLILKGILTAMGLIGIICLT   922
tr|X1WHI8|X1WHI8_DANRE                SNIASASMILPNPKPPGIS------------------NPGLNLTVLVSSLCAVTIVICII   886
tr|A0A8M9Q5D8|A0A8M9Q5D8_DANRE        SNVASGLQIIPNPNTGDL--------------------DLKLIFIAISVCLASVVVIGI   872
tr|A0A8M3APN3|A0A8M3APN3_DANRE        SNVASGLQIIPNPNTGDL--------------------DLKLIFIAISVCLASVVVIGI   886
                                      **:*                                  .  :    .      :
                                                     "PTS"                    Transmembrane helix

sp|A8K7I4|CLCA1_HUMAN                 ------------------------- 914
sp|Q6PT52|CLCA1_MACMU                 ------------------------- 913
sp|Q2TU62|CLCA1_HORSE                 G------------------------ 913
sp|Q9TUB5|CLCA1_PIG                   ------------------------- 917
tr|F1MGZ5|F1MGZ5_BOVIN                ------------------------- 909
tr|A6HWA3|A6HWA3_RAT                  H------------------------ 910
sp|Q9D7Z6|CLCA1_MOUSE                 H------------------------ 913
tr|F6S0F1|F6S0F1_MONDO                ESTT--------------------- 841
tr|A0A2U3V5T1|A0A2U3V5T1_TURTR        G------------------------ 913
sp|Q9UQC9|CLCA2_HUMAN                 IVVTHHTLSRKKRADKKENGTKLL 943
sp|Q9Y6N3|CLCA3_HUMAN                 ------------------------- 262
sp|Q14CN2|CLCA4_HUMAN                 LSTTI-------------------- 919
sp|Q8BG22|CLCA2_MOUSE                 MVVAHCIFNRKKRPSRKENETKFL 942
sp|Q9QX15|CA3A1_MOUSE                 LN----------------------- 902
sp|Q6Q473|CLA4A_MOUSE                 VGTTICILKNKRSSSAAITK--F- 924
tr|F6TR44|F6TR44_HORSE                IVVTHCILNKKKTADNKENGMKLL 946
tr|X1WHI8|X1WHI8_DANRE                AGVTTWAVRRRRPALRI-------- 903
tr|A0A8M9Q5D8|A0A8M9Q5D8_DANRE        FGITTWALTRKKLSDS--------- 888
tr|A0A8M3APN3|A0A8M3APN3_DANRE        FGITTWALTRKKLSDS--------- 902

                                        Transmembrane helix
```

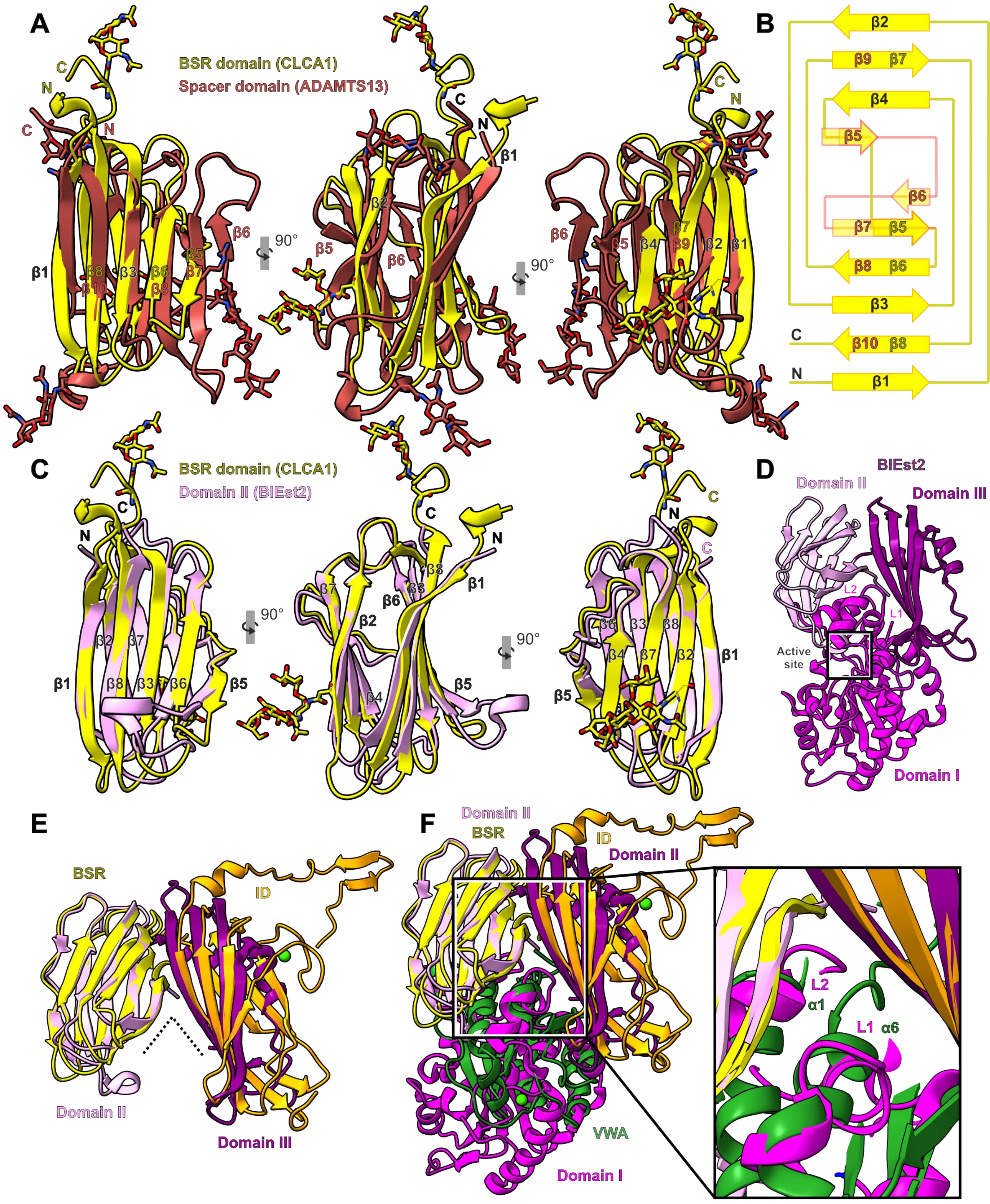
A
BSR domain (CLCA1)
Spacer domain (ADAMTS13)
90°
B
C
N
BSR domain (CLCA1)
Domain II (BlEst2)
D
BlEst2
Domain II
Domain III
L2
L1
Active site
Domain I
E
BSR
ID
Domain II
Domain III
F
Domain II
BSR
ID
Domain II
VWA
Domain I
L2
α1
L1
α6

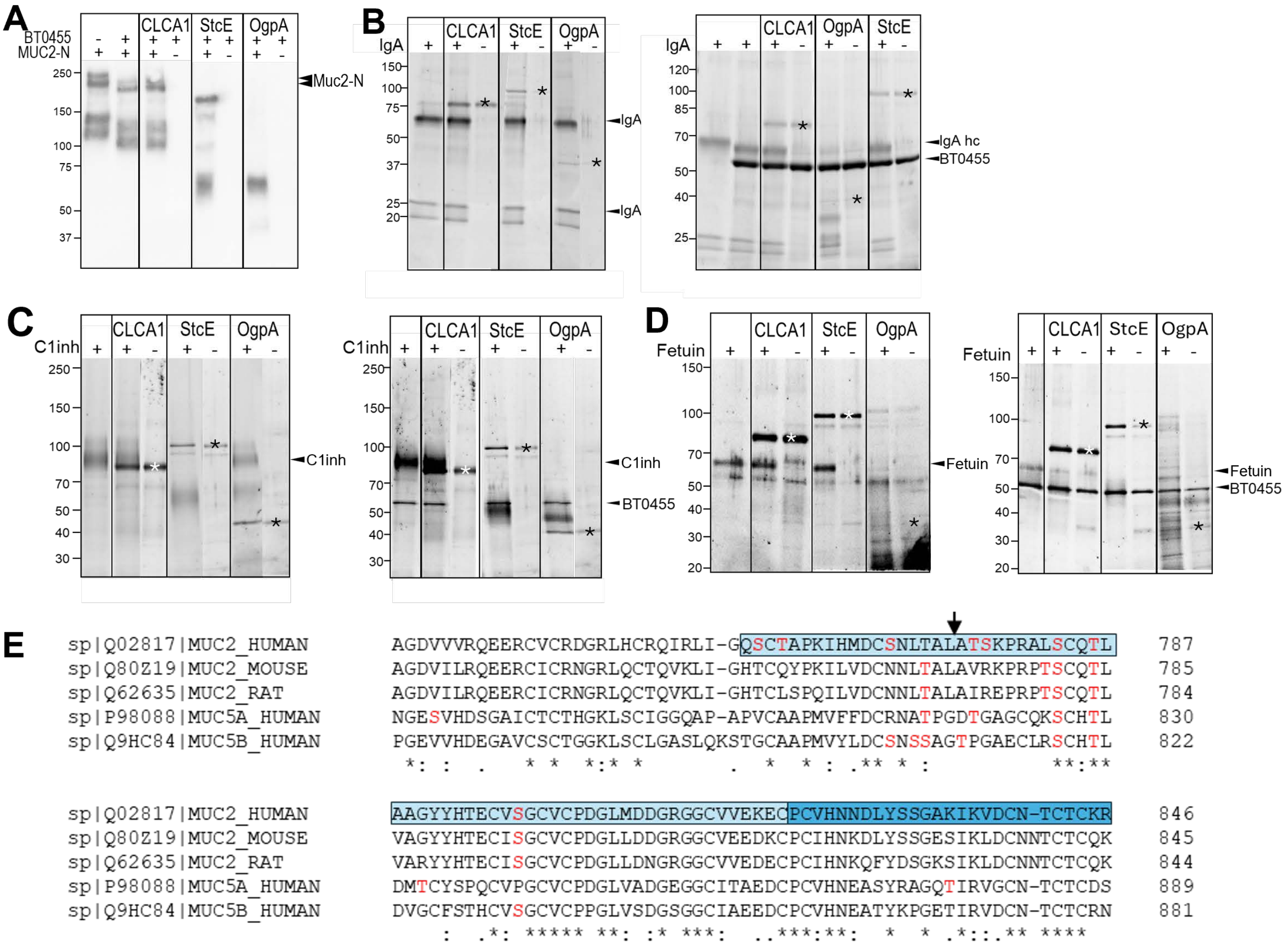

A
BT0455
MUC2-N
CLCA1
StcE
OgpA
Muc2-N
250
150
100
75
50
37
B
IgA
CLCA1
StcE
OgpA
IgA
IgA hc
BT0455
150
120
100
85
75
70
50
40
37
25
20
C
C1inh
CLCA1
StcE
OgpA
C1inh
BT0455
250
150
100
70
50
40
30
D
Fetuin
CLCA1
StcE
OgpA
Fetuin
BT0455
150
100
70
50
40
30
20
E
sp|Q02817|MUC2_HUMAN AGDVVVRQEERCVCRDGRLHCRQIRLI-GQSCTAPKIHMDCSNLTALATSKPRALSCQTL 787
sp|Q80Z19|MUC2_MOUSE AGDVILRQEERCICRNGRLQCTQVKLI-GHTCQYPKILVDCNNLTALAVRKPRPTSCQTL 785
sp|Q62635|MUC2_RAT AGDVILRQEERCICRNGRLQCTQVKLI-GHTCLSPQILVDCNNLTALAIREPRPTSCQTL 784
sp|P98088|MUC5A_HUMAN NGESVHDSGAICTCTHGKLSCIGGQAP-APVCAAPMVFFDCRNATPGDTGAGCQKSCHTL 830
sp|Q9HC84|MUC5B_HUMAN PGEVVHDEGAVCSCTGGKLSCLGASLQKSTGCAAPMVYLDCSNSSAGTPGAECLRSCHTL 822
sp|Q02817|MUC2_HUMAN AAGYYHTECVSGCVCPDGLMDDGRGGCVVEKECPCVHNNDLYSSGAKIKVDCN-TCTCKR 846
sp|Q80Z19|MUC2_MOUSE VAGYYHTECISGCVCPDGLLDDGRGGCVEEDKCPCIHNKDLYSSGESIKLDCNNTCTCQK 845
sp|Q62635|MUC2_RAT VARYYHTECISGCVCPDGLLDNGRGGCVVEDECPCIHNKQFYDSGKSIKLDCNNTCTCQK 844
sp|P98088|MUC5A_HUMAN DMTCYSPQCVPGCVCPDGLVADGEGGCITAEDCPCVHNEASYRAGQTIRVGCN-TCTCDS 889
sp|Q9HC84|MUC5B_HUMAN DVGCFSTHCVSGCVCPPGLVSDGSGGCIAEEDCPCVHNEATYKPGETIRVDCN-TCTCRN 881
