## Supplementary Table S1 for "Cryo-EM structure of CLCA1 identifies CLCA1 as a founding member of a novel metzincin family"

**Supplementary Table S1.** CLCA1 cryo-electron microscopy parameters.

| | |
|---|---|
| **Data Accession** | |
| PDB | 9R2T |
| EMDB | EMD-53539 |
| **Data Collection** | |
| Microscope | TFS KRIOS |
| Voltage (kV) | 300 |
| Detector | GATAN K3 |
| Pixel Size (Å) | 0.86 |
| Electron exposure ($e^-$/Å$^2$) | 50 |
| Defocus range (µm) | -0.5 to -2.5 |
| Micrographs | 11,634 |
| **Reconstruction** | |
| Software | CryoSPARC (v3.2) |
| Micrographs used | 8,282 |
| Particles used in refinement | 337,181 |
| Symmetry imposed | C1 |
| Overall resol. (Å) FSC=0.143 (masked) | 2.99 |
| Map sharpening B-factor (Å$^2$) | -120.8 |
| Local resol. rang.(Å) | 2.016-18.416 |
| **Model Refinement** | |
| Software | Phenix (v1.21.2-5419) |
| Non-hydrogen atoms | 6,676 |
| Protein residues | 834 |
| Ligands | 23 |
| *Av. B factors (Å$^2$)* | |
| Protein | 147.01 |
| Ligands | 236.57 |
| *R.M.S. deviations* | |
| Bond length (Å) | 0.002 |
| Bond angle (°) | 0.517 |
| *Ramachandran statistics (%)* | |
| Outliers | 0.00 |
| Allowed | 1.57 |
| Favored | 98.43 |
| MolProbity score | 1.01 |
| *Model vs. Map FSC* | |
| FSC=0.5 (masked,Å) | 3.2 |
