## Supplementary Table S2 for "Cryo-EM structure of CLCA1 identifies CLCA1 as a founding member of a novel metzincin family"

**Supplementary Table S2.** Identified *N*-glycopeptides by mass spectrometry

| Column name | Description |
|---|---|
| **Spectrum** | MS/MS spectrum identifier, follows the format (file name).(scan #).(scan #).(charge) |
| **Peptide** | Peptide amino acid sequence, any modifications not included ('stripped' peptide sequence) |
| **Modified Peptide** | peptide sequence including modifications, modified residues are followed by brackets containing the integer mass (in Da) of the residue plus the modification; blank if peptide is unmodified |
| **Prev AA** | Residue preceding the identified peptide within the mapped protein sequence; - if none |
| **Next AA** | Residue following the identified peptide within the mapped protein sequence; - if none |
| **Peptide Length** | Number of residues in the peptide sequence |
| **Charge** | Charge state of the identified peptide |
| **Retention** | MS2 scan's precursor retention time (in seconds) |
| **Observed Mass** | Mass of the identified peptide (in Da) |
| **Calibrated Observed Mass** | Mass of the identified peptide after m/z calibration (in Da) |
| **Observed M/Z** | Mass-to-charge ratio of the peptide ion |
| **Calibrated Observed M/Z** | Mass-to-charge ratio of the peptide ion after m/z calibration |
| **Calculated Peptide Mass** | Theoretical peptide mass based on identified sequence and modifications |
| **Calculated M/Z** | Theoretical peptide mass-to-charge ratio based on identified sequence and modifications |
| **Delta Mass** | Difference between calibrated observed peptide mass and calculated peptide mass (in Da) |
| **Expectation** | Expectation value from statistical modeling with PeptideProphet, lower values indicate higher likelihood |
| **Hyperscore** | Similarity score between observed and theoretical spectra, higher values indicate greater similarity |
| **Nextscore** | Similarity score (hyperscore) of second-highest scoring match for the spectrum |
| **PeptideProphet Probability** | Confidence score determined by PeptideProphet, higher values indicate greater confidence |
| **Number of Enzymatic Termini** | 2 = fully-enzymatic, 1 = semi-enzymatic, 0 = non-enzymatic |
| **Number of Missed Cleavages** | Number of potential enzymatic cleavage sites within the identified sequence |
| **Protein Start** | Starting position of the identified peptide within the protein sequence |
| **Protein End** | Ending position of the identified peptide within the protein sequence |
| **Intensity** | Precursor abundance (area under the curve) for each PSM if IonQuant is used; or maximum MS1 peak intensity within the retention time tolerance if Philosopher freequant is used (not recommanded). |
| **Ion Mobility** | TIMS transit time of the precursor ion ($1/K_0$) |
| **Assigned Modifications** | Variable modifications (listed by mass in Da) with modified residue and location within the peptide |
| **MSFragger Localization** | MSFragger-determined localization for open/offset searches, if using localize_delta_mass. Lower case letter(s) indicate localized site(s). |
| | More than one lower case letter indicates ambiguous localization. If all letters are upper case, the unlocalized candidate got a higher score and no localization information is known. |
| **Best Score with Delta Mass** | Highest observed hyperscore when the Delta Mass is placed on the theoretical spectrum (from open/offset search) |
| **Best Score without Delta Mass** | Highest hyperscore observed without placing the Delta Mass on the theoretical spectrum (from open/offset search) |
| **Glycan Score** | Score assigned to the glycan composition. Higher is better. |
| **Glycan q-value** | Q-value for the glycan composition assignment from the glycan FDR calculation. NOTE: all PSMs that pass peptide FDR are reported, even if the glycan FDR is not passed. Filter this column to q-value < 0.01 for a 1% glycan FDR (for example). |
| **Purity** | Proportion of total ion abundance in the inclusion window from the precursor (including precursor isotopic peaks), from Philosopher freequant |
| **Is Unique** | Whether the identified sequence maps to a single identified protein (FALSE if shared between multiple proteins identified in the experiment) |
| **Protein** | Protein sequence header corresponding to the identified peptide sequence; this will be the selected razor protein if the peptide maps to multiple proteins (in this case, other mapped proteins are listed in the 'Mapped Proteins' column). |
| **Protein ID** | Protein ID identifier (primary accession number) for the selected protein |
| **Entry Name** | Entry Name for the selected protein |
| **Gene** | Gene name for the selected protein |
| **Protein Description** | Name of the selected protein |
| **Mapped Genes** | Additional genes the identified peptide may originate from |
| **Mapped Proteins** | Additional proteins the identified peptide maps to (including any arising from I/L substitutions) |
