## Supplementary Table S3 for "Cryo-EM structure of CLCA1 identifies CLCA1 as a founding member of a novel metzincin family"

**Supplementary Table S3.** DALI search. Top hits of DALI PDB25 server search using CLCA1-MH (amino acid 22-302) as input structure.

| | Chain | Z | rmsd | lali | nres | %id | Description | Clan |
|---|---|---|---|---|---|---|---|---|
| 1 | 6o38-A | 7.3 | 3.5 | 138 | 573 | 9 | ACINETOBACTER SECRETED PROTEASE CPAA | M72 |
| 2 | 8snl-A | 7.3 | 3.8 | 161 | 656 | 11 | ADAM17 | M12 |
| 3 | 6qig-A | 7.1 | 3.9 | 140 | 581 | 11 | ADAMTS-13 | M12 |
| 4 | 6r7u-B | 7.0 | 3.7 | 156 | 308 | 12 | MIROLYSIN | M42 |
| 5 | 2dw1-B | 7.0 | 4.0 | 144 | 415 | 13 | CATROCOLLASTATIN | M12 |
| 6 | 8esv-A | 7.0 | 4.2 | 163 | 446 | 15 | ADAM10 | M12 |
| 7 | 3g5c-A | 6.6 | 4.4 | 145 | 486 | 13 | ADAM 22 | M12 |
| 8 | 7ufg-A | 6.5 | 4.0 | 149 | 1151 | 12 | PAPPALYSIN-1 | M43 |
| 9 | 6z2p-A | 6.1 | 3.9 | 153 | 356 | 5 | OGPA FROM AKKERMANSIA MUCINIPHILA | M11 |
| 10 | 4hx3-A | 6.1 | 3.6 | 115 | 134 | 10 | SNAPALYSIN | M07 |
| 11 | 2x7m-A | 5.8 | 3.8 | 129 | 180 | 13 | ARCHAEMETZINCIN | M54 |
| 12 | 4yu6-A | 5.7 | 3.5 | 146 | 754 | 11 | IMMUNE INHIBITOR 2A | M6 |
| 13 | 8dek-A | 5.6 | 3.9 | 158 | 440 | 11 | AMUC_1438 | unclassified |
| 14 | 8a7e-C | 5.2 | 4.1 | 149 | 1523 | 13 | PAPPALYSIN-1 | M43 |
| 15 | 1su3-B | 5.2 | 4.1 | 126 | 416 | 12 | MMP1 | M10 |
| … | | | | | | | | |
| 19 | 3p1v-A | 4.9 | 3.8 | 142 | 406 | 15 | BACOVA_00662 | M64 |
| … | | | | | | | | |
| 113 | 3ujz-A | 2.5 | 5.1 | 131 | 604 | 9 | StcE | M66 |
