## Supplementary Table S4 for "Cryo-EM structure of CLCA1 identifies CLCA1 as a founding member of a novel metzincin family"

**Supplementary Table S4.** Mucins *O*-glycosylation sites.

| Human MUC2 | | | Mouse Muc2 | | | Rat Muc2 | | | Human MUC5AC | | | Human MUC5B | | |
|---|---|---|---|---|---|---|---|---|---|---|---|---|---|---|
| Position | Score | Sub-domain | Position | Score | Sub-domain | Position | Score | Sub-domain | Position | Score | Sub-domain | Position | Score | Sub-domain |
| 25 | 0,5832 | VWD1 | 215 | 0,5884 | C8-1 | 214 | 0,6070 | C8-1 | 24 | 0,5066 | VWD1 | 25 | 0,6108 | VWD1 |
| 217 | 0,5991 | C8-1 | 229 | 0,7239 | C8-1 | 228 | 0,7488 | C8-1 | 31 | 0,6077 | VWD1 | 46 | 0,8929 | VWD1 |
| 276 | 0,5528 | C8-1 | 285 | 0,5246 | C8-1 | 273 | 0,5183 | C8-1 | 32 | 0,5428 | VWD1 | 47 | 0,7331 | VWD1 |
| 287 | 0,6296 | C8-1 | 316 | 0,6480 | TIL1 | 284 | 0,7098 | C8-1 | 43 | 0,7016 | VWD1 | 48 | 0,8403 | VWD1 |
| 289 | 0,5593 | C8-1 | 317 | 0,6676 | TIL1 | 286 | 0,6814 | C8-1 | 50 | 0,9028 | VWD1 | 50 | 0,7692 | VWD1 |
| 304 | 0,6547 | TIL1 | 547 | 0,6533 | VWD2 | 301 | 0,6323 | TIL1 | 58 | 0,8483 | VWD1 | 54 | 0,9565 | VWD1 |
| 318 | 0,6485 | TIL1 | 573 | 0,5318 | C8-2 | 315 | 0,5000 | TIL1 | 62 | 0,8762 | VWD1 | 60 | 0,8434 | VWD1 |
| 319 | 0,7043 | TIL1 | 592 | 0,7475 | C8-2 | 316 | 0,6577 | TIL1 | 65 | 0,7151 | VWD1 | 64 | 0,7990 | VWD1 |
| 549 | 0,8077 | VWD2 | 606 | 0,5000 | C8-2 | 539 | 0,5000 | VWD2 | 72 | 0,6691 | VWD1 | 265 | 0,5440 | C8-1 |
| 556 | 0,5369 | VWD2 | 661 | 0,6010 | TIL2 | 546 | 0,7452 | VWD2 | 319 | 0,5569 | C81 | 340 | 0,5476 | TIL1 |
| 575 | 0,5437 | C8-2 | 662 | 0,6982 | TIL2 | 591 | 0,6928 | C8-2 | 361 | 0,6040 | TIL1 | 347 | 0,5582 | TIL1 |
| 594 | 0,5737 | C8-2 | 671 | 0,6241 | TIL2 | 674 | 0,6849 | TIL2 | 386 | 0,5432 | TIL1 | 388 | 0,6853 | TIL1 |
| 603 | 0,6548 | C8-2 | 675 | 0,6944 | TIL2 | 677 | 0,8094 | TIL2 | 408 | 0,5450 | E1 | 393 | 0,8014 | E1 |
| 677 | 0,5761 | TIL2 | 678 | 0,8675 | TIL2 | 679 | 0,8138 | TIL2 | 584 | 0,6072 | C8-2 | 395 | 0,8977 | E1 |
| 680 | 0,8145 | TIL2 | 680 | 0,7732 | TIL2 | 683 | 0,5545 | TIL2 | 586 | 0,5000 | C8-2 | 398 | 0,6524 | E1 |
| 682 | 0,6291 | TIL2 | 684 | 0,5821 | TIL2 | 769 | 0,5963 | TIL´ | 592 | 0,8902 | C8-2 | 399 | 0,8608 | E1 |
| 686 | 0,5293 | TIL2 | 770 | 0,6687 | TIL´ | 779 | 0,8860 | TIL´ | 599 | 0,5761 | C8-2 | 583 | 0,8531 | C8-2 |
| 758 | 0,5964 | TIL´ | 780 | 0,9626 | TIL´ | 780 | 0,9348 | TIL´ | 602 | 0,6635 | C8-2 | 590 | 0,6239 | C8-2 |
| 760 | 0,5585 | TIL´ | 781 | 0,9489 | TIL´ | 783 | 0,9254 | TIL´ | 612 | 0,5320 | C8-2 | 711 | 0,6489 | TIL2 |
| 769 | 0,7073 | TIL´ | 784 | 0,9357 | TIL´ | 1001 | 0,5620 | VWD3 | 698 | 0,6162 | TIL2 | 716 | 0,6872 | TIL2 |
| 776 | 0,8760 | TIL´ | 1002 | 0,5830 | VWD3 | 1010 | 0,5528 | VWD3 | 702 | 0,8907 | TIL2 | 721 | 0,6034 | TIL2 |
| 777 | 0,8955 | TIL´ | 1011 | 0,6384 | VWD3 | 1022 | 0,6862 | VWD3 | 703 | 0,8134 | TIL2 | 723 | 0,7536 | TIL2 |
| 783 | 0,9293 | TIL´ | 1023 | 0,7539 | VWD3 | 1023 | 0,7008 | VWD3 | 707 | 0,6424 | TIL2 | 725 | 0,7586 | TIL2 |
| 786 | 0,9454 | TIL´ | 1024 | 0,7923 | VWD3 | 1028 | 0,8895 | VWD3 | 709 | 0,8168 | TIL2 | 733 | 0,5647 | TIL2 |
| 798 | 0,5000 | TIL´ | 1029 | 0,9177 | VWD3 | 1035 | 0,8553 | C8-3 | 715 | 0,8959 | TIL2 | 791 | 0,6389 | E2 |
| 831 | 0,6436 | E´ | 1036 | 0,8651 | C8-3 | 1042 | 0,7265 | C8-3 | 716 | 0,8811 | TIL2 | 804 | 0,5499 | TIL´ |
| 999 | 0,5448 | VWD3 | 1043 | 0,7287 | C8-3 | 1141 | 0,5501 | TIL3 | 720 | 0,7503 | TIL2 | 806 | 0,5474 | TIL´ |
| 1004 | 0,5532 | VWD3 | 1054 | 0,5000 | C8-3 | 1144 | 0,5000 | TIL3 | 723 | 0,8233 | TIL2 | 807 | 0,7283 | TIL´ |
| 1013 | 0,6516 | VWD3 | 1142 | 0,5535 | TIL3 | 1150 | 0,5252 | TIL3 | 725 | 0,6799 | TIL2 | 810 | 0,8344 | TIL´ |
| 1026 | 0,8806 | VWD3 | 1145 | 0,5330 | TIL3 | 1153 | 0,7488 | TIL3 | 732 | 0,6390 | TIL2 | 818 | 0,7527 | TIL´ |
| 1031 | 0,9397 | VWD3 | 1151 | 0,5303 | TIL3 | 1198 | 0,5429 | E3 | 759 | 0,6220 | TIL2 | 821 | 0,6702 | TIL´ |
| 1032 | 0,5612 | C8-3 | 1154 | 0,7669 | TIL3 | | | | 768 | 0,6451 | E2 | 833 | 0,6693 | TIL´ |
| 1038 | 0,9343 | C8-3 | 1199 | 0,6148 | E3 | | | | 775 | 0,7064 | E2 | 863 | 0,5754 | E´ |
| 1045 | 0,5837 | C8-3 | | | | | | | 815 | 0,8479 | TIL´ | 869 | 0,7550 | E |
| 1064 | 0,5311 | C8-3 | | | | | | | 819 | 0,7651 | TIL´ | 887 | 0,5502 | E |
| 1144 | 0,5883 | TIL3 | | | | | | | 826 | 0,8546 | TIL´ | 1041 | 0,5499 | VWD3 |
| 1153 | 0,5313 | TIL3 | | | | | | | 829 | 0,6210 | TIL´ | 1058 | 0,6169 | VWD3 |
| 1156 | 0,7636 | TIL3 | | | | | | | 833 | 0,6097 | TIL´ | 1060 | 0,6717 | VWD3 |
| 1194 | 0,5148 | E3 | | | | | | | 877 | 0,7630 | E´ | 1072 | 0,7243 | C8-3 |
| 1201 | 0,8169 | E3 | | | | | | | 895 | 0,5901 | E´ | 1079 | 0,6796 | C8-3 |
| | | | | | | | | | 948 | 0,5374 | VWD3 | 1092 | 0,5665 | C8-3 |
| | | | | | | | | | 1051 | 0,5524 | VWD3 | 1183 | 0,5718 | TIL3 |
| | | | | | | | | | 1053 | 0,5262 | VWD3 | 1203 | 0,5599 | TIL3 |
| | | | | | | | | | 1068 | 0,5260 | VWD3 | | | |
| | | | | | | | | | 1070 | 0,6885 | VWD3 | | | |
| | | | | | | | | | 1082 | 0,6816 | C8-3 | | | |
| | | | | | | | | | 1089 | 0,7176 | C8-3 | | | |
| | | | | | | | | | 1102 | 0,6708 | C8-3 | | | |
| | | | | | | | | | 1188 | 0,5970 | TIL3 | | | |

Amino acids in the N-terminal region (amino acid 1-1200) of MUC2 (human, mouse and rat), MUC5AC and MUC5B (both human) with positive (>0.5) score in NetOGlyc 4.0 server prediction of *O*-glycosylation sites. Scores are conditionally formatted with red being the highest and blue the lowest score in the list. The corresponding sub domain is included, based on (Javitt et al. 2020).
